# Random *in vitro* Protein single-Loop Engineering (RiPLE) by mRNA display

**DOI:** 10.64898/2026.09.21.753128

**Authors:** Kilian Colas, Tongil Ko, Grégoire Philippe, Hiroaki Suga

## Abstract

Peptide lasso-grafting strategies enable the transfer of established pharmacophores into compact protein scaffolds but often require optimization to preserve target binding and biological activity. Here, we report Random in-vitro Protein single-Loop Engineering (RiPLE), an mRNA-display strategy for the direct discovery of engineered ubiquitin binders, termed U-bodies, from a library containing a fully randomized surface loop of variable length. Using receptor tyrosine kinase-like orphan receptor 1 (ROR1) as a model target, we generated U-bodies through RiPLE and compared them with lasso-grafted U-bodies derived from a previously identified ROR1-binding macrocyclic peptide. Both approaches yielded U-bodies bearing single-digit nanomolar affinity to ROR1; however, RiPLE converged on *de novo* binding sequences that were distinct from the parental peptide. Notably, most selected U-bodies exhibited strong selectivity for ROR1 over the closely related receptor ROR2, whereas the parental peptide bound both receptors with high affinity. Representative ROR1-binding U-bodies also inhibited wound closure in ROR1-expressing MDA-MB-231 cells without detectable cytotoxicity. Applying RiPLE to the heterotrimeric G-protein subunit (GαS) similarly yielded high-affinity binders and revealed sequence convergence patterns distinct from lasso-grafting. Together, these results establish RiPLE as a powerful and complementary alternative to peptide-guided lasso-grafting, enabling *de novo* evolution of functional binding surfaces directly within their final protein scaffold.

## Introduction

Designer proteins and macrocyclic peptides have emerged as a promising class of molecules that occupy the chemical space between small molecules and conventional biologics. By displaying target-binding surfaces on compact and structurally stable scaffolds, these molecules can achieve antibody-like affinity and specificity while retaining advantages associated with their smaller size.^1-4^ Such advantages include improved tissue and tumor penetration and, for scaffolds that lack disulfide bonds or multidomain architectures, straightforward production in prokaryotic expression systems.^5, 6^ Their relative structural simplicity also enables extensive sequence diversification, making them attractive starting points for the development of new therapeutic modalities.^7, 8^

A common strategy for generating target specificity is to diversify solvent-exposed surface residues to create semi-randomized libraries of protein variants, and subsequently identify binders using display-based selection. This principle is well established in antibody and nanobody engineering through diversification of complementarity-determining regions (CDRs), particularly CDR3.^9, 10^ The same principle has been extended to non-antibody scaffolds including FN3 domains, affibodies, and DARPins using phage, ribosome display.^11-13^ Although mRNA display has also been used on these scaffolds, it has largely been limited to evolving epitope-binding regions rather than engineering entirely new loops.^14,15^ Ubiquitin has likewise been adapted as a binding scaffold, giving rise to Affilin proteins and ubiquitin variants with high affinity for targets including HER3, Grb2, and the extradomain B of fibronectin.^16, 17^ More extensive modifications such as loop insertion, domain insertion, and circular permutation have also been used to alter protein topology, function, and ligand recognition.^18^

Although these approaches have enabled the discovery and optimization of many functional binders, display-based screening imposes practical constraints on the sequence space that can be explored. Cell-based platforms are limited by transformation efficiency, protein expression, and display efficiency, restricting library diversity and introducing sequence-dependent biases.^19^ Moreover, scaffold libraries are typically constructed by predefining a fixed set of positions for diversification, constraining the binding sequences that can emerge.^11, 13, 16^ Computational and rational design provide an alternative route, and recent structure-based and deep-learning approaches have enabled the generation of high-affinity de novo binders and increasingly diverse protein architectures.^20, 21^ Nevertheless, these methods generally depend on structural information or accurate models of the target, and designed candidates often require experimental screening and iterative optimization to achieve the desired affinity and biophysical properties.^22, 23^

In contrast to these approaches, mRNA display provides a fully *in vitro* platform capable of screening substantially larger libraries.^24, 25^ The technology has been particularly successful in the discovery of macrocyclic peptides through the Random nonstandard Peptides Integrated Discovery (RaPID) system, in which genetic code reprogramming provides access to nonproteinogenic amino acids and diverse cyclization chemistries.^26^ Despite their favorable target-recognition properties, macrocyclic peptides can exhibit poor serum stability, solubility, and bioavailability.^27, 28^ Protein-grafting strategies offer one way to address these limitations by transferring an established peptide pharmacophore onto a protein scaffold.^29^ In particular, lasso-grafting (LG) emerged as a versatile platform and has been applied to a wide range of protein architectures, including immunoglobulin (IgG) domains, serum albumin, and artificial protein capsids.^30, 31^ The resulting constructs can retain the binding ability of the grafted peptide pharmacophore while acquiring scaffold-derived advantages, such as improved stability, solubility, recombinant production, multivalency, or higher-order assembly.^32^

Nevertheless, the LG approach relies on the availability of a pre-existing peptide pharmacophore that must remain in its active conformation following incorporation into the protein scaffold. Achieving this often requires optimization of the spacer residues connecting the grafting site within the protein to the “pseudo-cyclic” peptide motif,^31, 33^ as the thioether bond present in the original macrocycle must also be replaced with proteinogenic structural or by the grafting scaffold itself. Although LG can frequently be implemented in large protein scaffolds, such as loops within the Fc domain of IgG, without substantial loss of the parental macrocycle’s binding ability, not all RaPID-derived peptides can be successfully converted into functional grafted proteins.^29^ Moreover, each peptide pharmacophore requires the design and construction of a bespoke grafting library, with no guarantee that the chosen library design will yield a functional graft. An alternative strategy is to evolve binding sequences directly within the protein scaffold itself, thereby eliminating peptide-specific graft-design requirements and allowing target recognition to emerge within its functional structural context.

Herein we report Random *in-vitro* Protein single-Loop Engineering (RiPLE), an mRNA-display strategy in which a solvent-exposed loop of ubiquitin is replaced with a fully randomized, variable-length sequence. This universal library enables *de novo* selection of target-binding U-bodies without a pre-existing peptide pharmacophore or subsequent peptide-to-scaffold graft optimization. We evaluated this strategy using receptor tyrosine kinase-like orphan receptor 1 (ROR1), a cell-surface receptor associated with cancer-cell survival, proliferation, and migration. Our group previously identified a ROR1-binding macrocyclic peptide by RaPID selection, enabling a direct comparison between peptide-guided LG and *de novo* discovery within the ubiquitin scaffold.^34^ Both strategies yielded single-digit nanomolar ROR1 binders; however, RiPLE converged on binding sequences distinct from the parental peptide. Most selected U-bodies also exhibited preferential recognition of ROR1 over the closely related receptor ROR2. Furthermore, representative compounds inhibited wound closure in ROR1-expressing cells without detectable cytotoxicity. We also compared RiPLE and LG against the heterotrimeric G-protein subunit (GαS), for which high-affinity U-bodies were identified without a pre-existing peptide starting point. Collectively, these results establish RiPLE and LG as complementary approaches for evolving engineered protein binders by mRNA display.

## Results

### RiPLE enables de novo discovery of ROR1-binding U-bodies

We designed an mRNA-display library based on the ubiquitin scaffold, whose compact structure, high solubility, recombinant expression, and tolerance of sequence modification in solvent-exposed loop at the β-1/ β -2 turn (loop 1) make it suitable for protein engineering. In the RiPLE library, native ubiquitin residues Thr7– Gly10 were replaced with fully randomized sequences ranging from 8 to 15 amino acids in length (Fig. 1a and SI Fig. S1). The diversified region was encoded using NNK degenerate codons, providing access to all 20 proteinogenic amino acids. Following transcription, the mRNA library was ligated to a puromycin-containing linker, translated in vitro, and reverse-transcribed to generate covalently linked phenotype/genotype complexes for iterative affinity selection. Unlike LG (Fig. 1b), in which an established peptide pharmacophore is incorporated into a protein scaffold and the surrounding sequence is subsequently optimized, RiPLE directly diversifies the replacement loop without specifying a starting recognition sequence. This design provides a theoretical library diversity of 10^13^ protein-mRNA fusions per milliliter of translation reaction.^35, 36^

**Figure 1.**
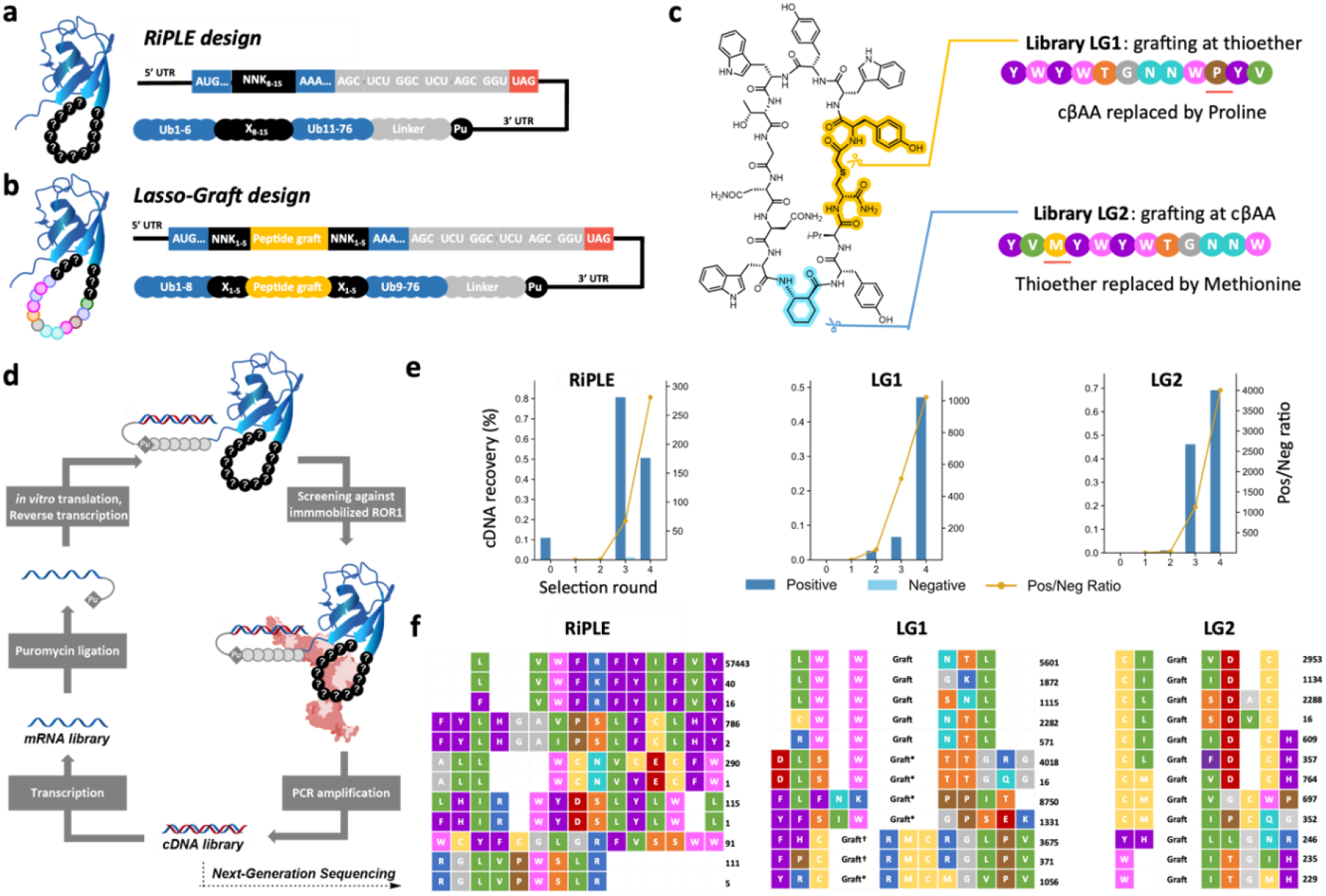
RiPLE and lasso-graft library design and selection against ROR1. a) Design of the RiPLE ubiquitin library. Native residues in ubiquitin loop 1 were replaced with fully randomized sequences 8–15 amino acids in length encoded by NNK codons, generating a variable-length binding surface without a predefined recognition sequence. b) General design of the lasso-graft ubiquitin library, in which a peptide pharmacophore is incorporated into loop 1 and flanked by 1–5 randomized residues on either side. c) Design of the two ROR1 lasso-graft libraries derived from the previously identified ROR1-binding macrocyclic peptide. LG1 was opened at the thioether linkage, with the nonproteinogenic cyclic β-amino acid (cβAA) replaced by proline, whereas LG2 was opened at the cβAA, with the thioether-forming residue replaced by methionine. d) Schematic of the mRNA-display workflow for affinity selection against immobilized ROR1. e) Enrichment of the RiPLE, LG1 and LG2 libraries during selection. Bars indicate cDNA recovery from positive and negative selections, and lines indicate the ratio of positive-to-negative recovery. f) Sequence distributions of enriched variants from the round 5 of RiPLE, LG1, and LG2 selections determined by next-generation sequencing. Sequence counts are shown at right; ‘graft’ denotes residues derived from the parental ROR1-binding peptide; ‘graft*’ contains a proline to leucine mutation; ‘graft^†^’ contains a deletion mutation of the last three residues.

We first evaluated RiPLE against receptor tyrosine kinase-like orphan receptor 1 (ROR1), for which we had previously identified a macrocyclic peptide B2 by RaPID selection.^34^ This enabled a direct comparison with peptide-guided LG. While source peptide B2 showed potent biological activity, it suffered from poor solubility and serum stability, making it a typical candidate for the LG alternative. However, the presence of three non-proteinogenic residues within the sequence, including a crucial cyclic-beta amino-acid (cβAA), complicates the design of the grafting library. Therefore, two LG libraries were designed using different opening points within the parental macrocycle (Fig. 1c). In the LG1 library, the peptide was opened at the thioether linkage, following the strategy used in our previous peptide-grafting studies.^29^ In the LG2 library, the peptide was instead opened at the cβAA, which we reasoned might favor a hairpin-like geometry more compatible with presentation from ubiquitin loop 1. To permit recombinant expression, nonproteinogenic residues were replaced with structurally related proteinogenic residues, including substitution of the cβAA with proline and replacement of the thioether-forming residue with methionine. The sequences flanking the grafted pharmacophore were diversified with 1–5 NNK codons to permit further optimization during selection.

The RiPLE, LG1, and LG2 libraries were independently subjected to iterative affinity selection against the Fc-tagged ROR1 extracellular domain immobilized on protein G magnetic beads (Fig. 1d). Negative selections against Fc-loaded protein G beads were used to reduce enrichment of nonspecific binders. All three libraries showed similar progressive enrichment with both the RiPLE and LG1/2 libraries showing a marked increase in positive recovery and the positive-to-negative recovery ratio after four rounds (Fig. 1e). Recovered DNA pools from rounds 3-5 were analyzed by next-generation sequencing to monitor sequence evolution and convergence. Round 5 populations are summarized in Fig. 1f, with the complete sequencing results provided in the Supplementary Table 1.

Sequencing revealed distinct outcomes among the three libraries. The round 5 RiPLE population was dominated by a single sequence, which accounted for 95.6% of the counts, alongside several lower-frequency families. Although unrelated in primary sequence to the original RaPID peptide, the selected loops were similarly enriched in aromatic and hydrophobic residues, particularly Trp, Phe, and Tyr. In contrast, the LG selections generally retained recognizable elements of the parental pharmacophore while accumulating changes within both the graft and its flanking regions. Notably, the proline introduced in LG1 as a proteinogenic replacement for the parental cβAA was frequently replaced by leucine, and enriched variants contained Trp-rich linker regions surrounding the retained pharmacophore. Sequencing of the LG2 population revealed several enriched variants containing paired cysteine residues in the two linker regions. Although engineered disulfide bridges have previously been used to stabilize grafted peptide conformations in LG engineering,^32^ the cysteine pairs observed here arose spontaneously during the selection. This pattern raises the possibility that an intramolecular disulfide bond generates an alternative constrained architecture that partially restores the conformational restriction of the parental macrocycle. Disulfide-constrained sequences also emerged in our previous U-body work, although they were not superior to other sequences lacking this feature.^31^ Thus, the two lasso-graft designs underwent distinct sequence adaptations despite originating from the same parental pharmacophore.

Representative variants from each library were recombinantly expressed (SI Fig. S3), and their binding to ROR1 was quantified by surface plasmon resonance (SPR, Table 1 and SI Fig. S5). Tested RiPLE variants bound ROR1 with low-nanomolar affinity (*K*_*D*_ = 1.1–11.7 nM). Several variants were characterized by particularly slow dissociation, including RL3 and RL4 with *k*_*on*_ values of approximately 0.9 × 10^−3^ s^−1^. LG1-derived U-bodies showed similarly strong binding, including the sub-nanomolar variant LG1-3 (*K*_*D*_ = 0.94 nM), which exhibited an even slower dissociation rate (*k*_*on*_ = 0.52 × 10^−3^ s^-1^). By contrast, the expressed LG2 variants displayed generally weaker affinities and dissociated more rapidly from ROR1. The lower affinities of the LG2 were associated primarily with reduced stability of the bound complex rather than a uniform loss of target association. It has been reported before that LG-U-bodies exhibit good serum and thermal stability as long as the core Ubiquitin residues are not mutated.^31^ We likewise found no detectable decomposition in any of these new U-bodies during thermal-shift assay. The serum stability of model RiPLE U-body RL1-5 was also measured at 58 h, in line with our previous reports (SI Fig. S4).

**Table 1.**
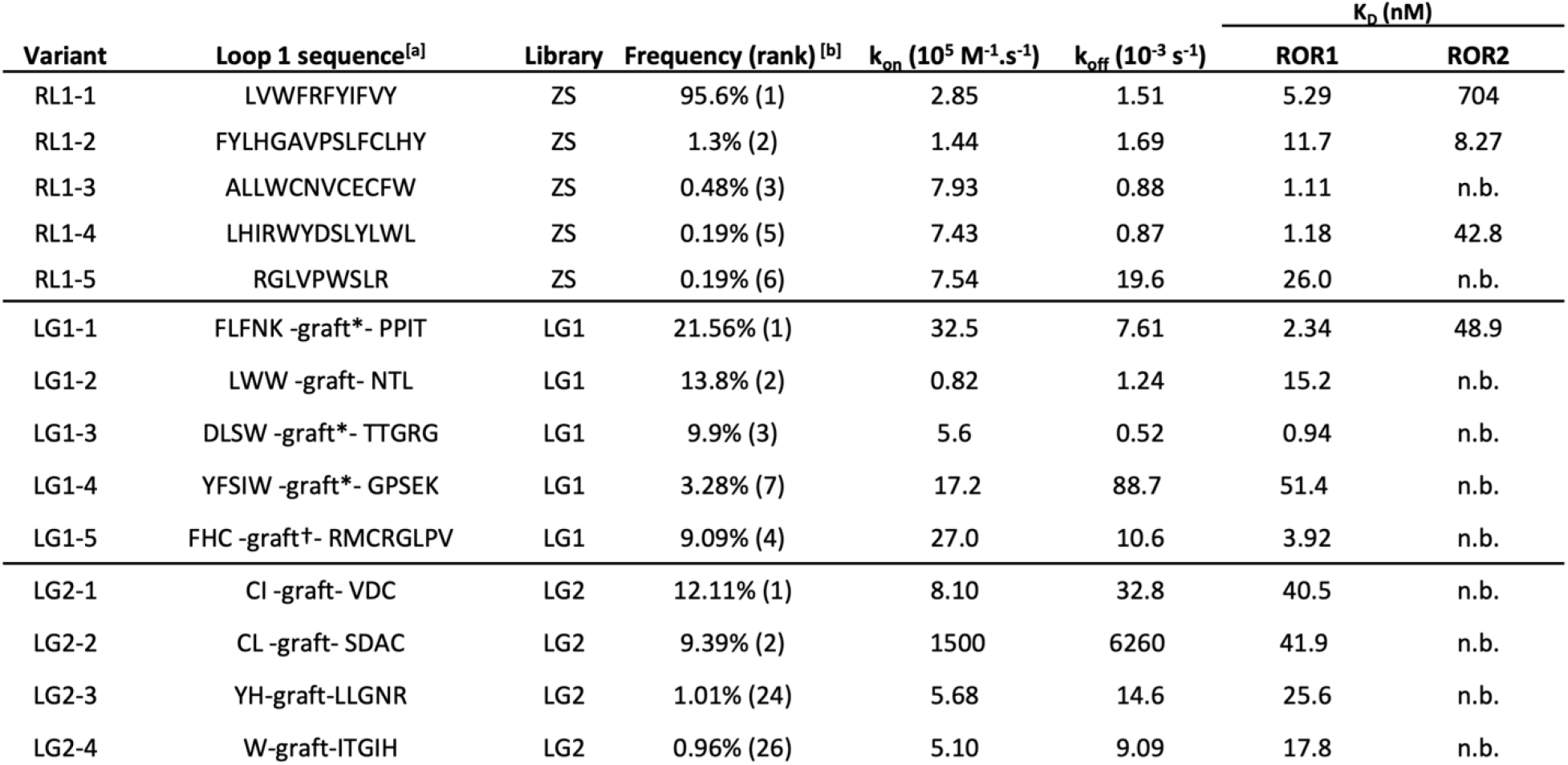
Sequence and binding properties of ROR1 U-bodies identified by RiPLE and lasso-grafting. Sequence frequencies and ranks correspond to the round-5 selection populations. SPR-derived *k*_*on*_ and *k*_*off*_ values refer to ROR1 binding; *K*_*D*_ values are shown for ROR1 and ROR2. n.b., no binding detected under the conditions tested.

We next assessed whether the selected U-bodies preferentially recognized ROR1 over the closely related receptor ROR2 by SPR (Table 1 and SI Fig. S6). Most tested U-bodies showed substantially weaker or undetectable binding to ROR2. This was notable because the parental macrocyclic peptide used for lasso-grafting bound both receptors with high affinity (ROR1 *K*_*D*_ = 1.2 nM; ROR2 *K*_*D*_ = 5.0 nM). This is consistent with our prior observation that the activity and selectivity of the source peptide was sensitive to subtle sequence changes. For variants in which ROR2 binding could be quantified, the reduced affinity was accompanied by markedly faster dissociation, particularly among the RiPLE-derived U-bodies. Thus, the selected U-bodies not only retained high-affinity recognition of ROR1 but also showed kinetic discrimination between the two closely related receptors. The structural basis for this selectivity is examined in a subsequent section.

### RiPLE is transferable across targets and complements lasso-grafting

To determine whether the behavior of RiPLE observed against ROR1 extended to other protein targets, we next applied the strategy to GαS, for which a macrocyclic peptide ligand had also been previously identified by RaPID selection.^37^ Both selections were performed against GppNHp-bound GαS(R201C), which carries an activating mutation that impairs GTP hydrolysis and mimics GTP-like active state. An LG library (LG3) was generated by incorporating a proteinogenic analogue of the original GαS-binding peptide GN13 into ubiquitin loop 1 (SI Fig. S2), while the same general RiPLE library design used for ROR1 was independently selected against GαS.

Both selections enriched and had converged by round 4 (Fig. 2a), but their sequence distributions differed from those observed for ROR1. Whereas the ROR1 RiPLE selection became dominated by a single sequence and the corresponding lasso-graft libraries retained several abundant variants, the GαS selections showed the opposite pattern. The RiPLE selection yielded a broader collection of enriched loop sequences, while the LG3 population converged more strongly towards one major family (Fig. 2b). Thus, the degree of convergence observed with RiPLE differed between the ROR1 and GαS selections, indicating that strong convergence is not an inherent feature of the library design.

**Figure 2.**
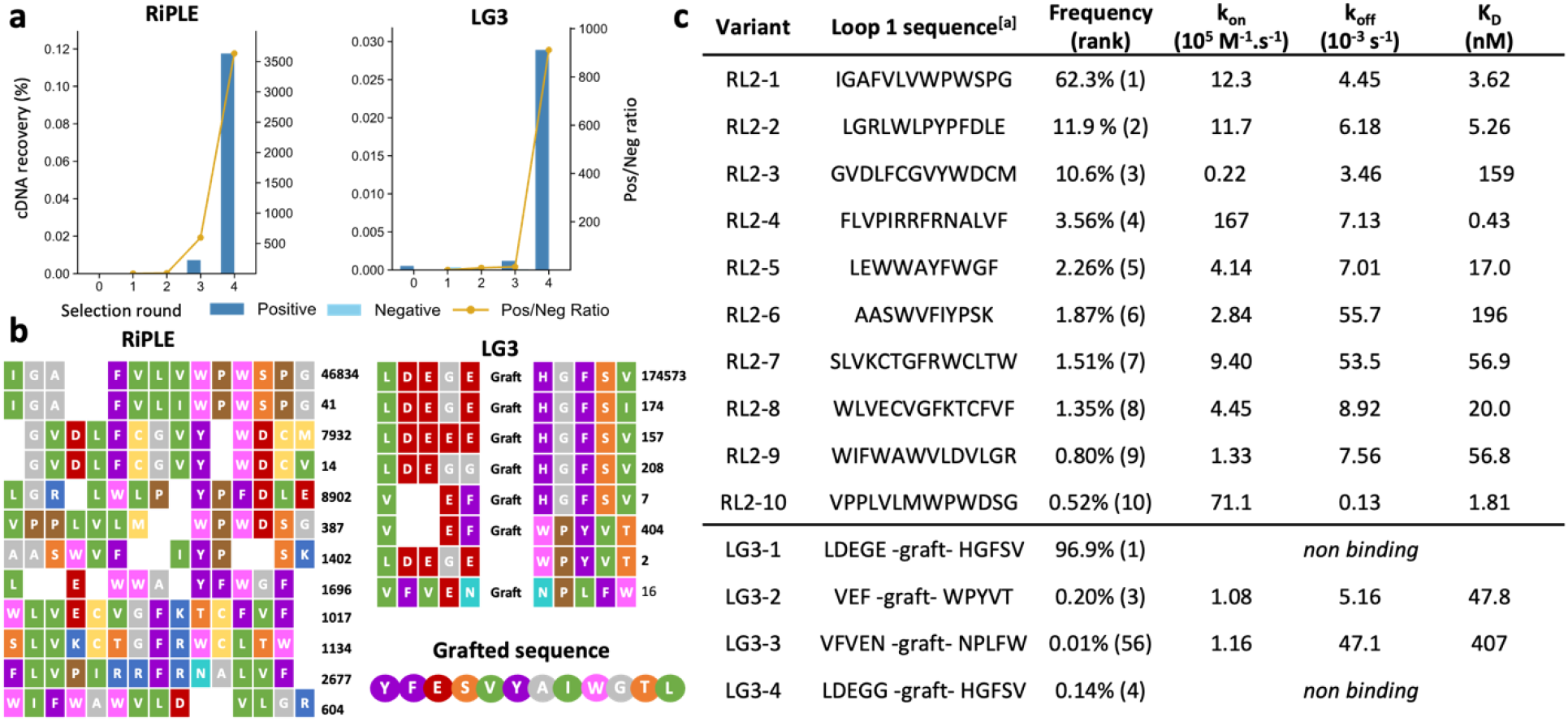
RiPLE generates high-affinity U-bodies against GppNHp-bound GαS. a) Enrichment of independently selected RiPLE and lasso-graft (LG3) libraries against GppNHp-bound GαS. Bars indicate cDNA recovery from positive and negative selections, and lines indicate the positive-to-negative recovery ratio. b) Sequence distributions of enriched RiPLE and LG3 variants after round 4 determined by next-generation sequencing. Sequence counts are shown at right; ‘Graft’ denotes residues derived from the parental GαS-binding macrocyclic peptide. c) SPR characterization of representative RiPLE- and LG3-derived U-bodies binding to GppNHp-bound GαS. Association rate constants (*k*_*on*_), dissociation rate constants *(k*_*off*_) and equilibrium dissociation constants (*K*_*D*_) are shown. Variants for which no binding was detected under the conditions tested are indicated as non-binding (n.b.).

Representative variants from both GαS selections were recombinantly expressed and characterized by SPR (Fig. 2c and SI Fig. S7). The RiPLE selection yielded several high-affinity binders, including RL2-4 (*K*_*D*_ = 0.43 nM), RL2-10 (*K*_*D*_ = 1.81 nM), and RL2-1 (*K*_*D*_ = 3.62 nM), together with additional variants spanning the low-to tens-of-nanomolar range. By comparison, the characterized LG3 variants generally displayed significantly weaker binding: LG3-2 and LG3-3 bound with *K*_*D*_ values of 47.8 and 407 nM, respectively, while LG3-1 and LG3-4 showed no detectable binding. The lack of binding for LG3-1 is particularly notable given its abundance in the sequence pool, perhaps reflecting the unsuitability of the source peptide for grafting. These results show that RiPLE can generate high-affinity binders even when lasso-grafting a previously validated peptide yields suboptimal results.

### Structural modeling suggests a basis for ROR1/ROR2 selectivity

The preferential binding of the selected U-bodies to ROR1 over ROR2 prompted us to investigate a possible structural basis for this selectivity. Although the structure of the ROR1 cysteine-rich domain (CRD) has not been experimentally determined, the corresponding ROR2 CRD has recently been crystallized.^38^ Comparison of the ROR2 structure with an AlphaFold3 model of ROR1 revealed a highly similar overall fold, including an additional α5 helix positioned over the hydrophobic cavity corresponding to the Wnt binding interface of Frizzled receptors (pseudo-site 1; Fig. 3a).

**Figure 3.**
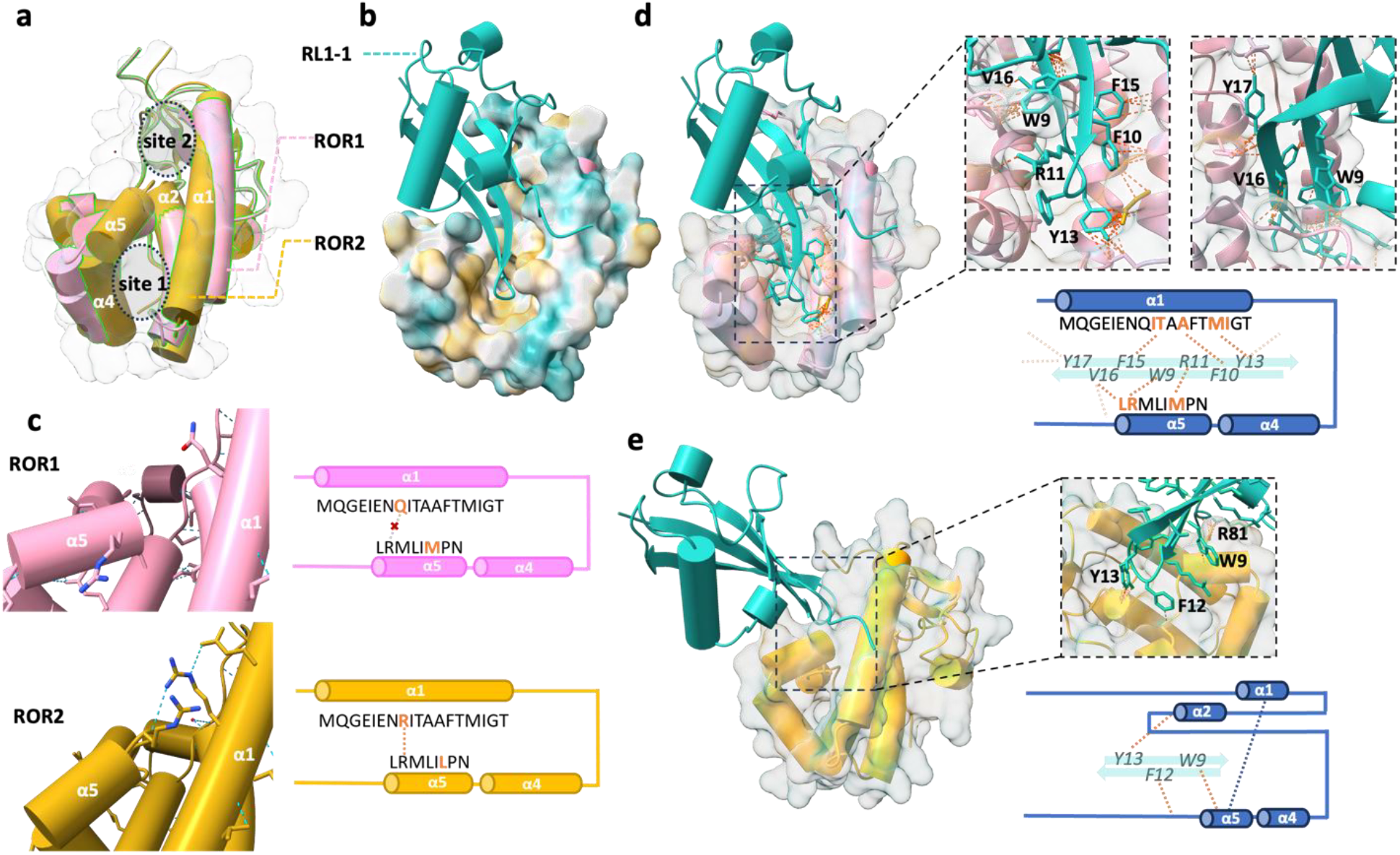
Structural modeling suggests a basis for discrimination between ROR1 and ROR2. a) AlphaFold3-predicted structure of the ROR1 cysteine-rich domain (CRD; pink) superimposed with the experimentally determined ROR2 CRD structure (orange). Pseudo-sites 1 and 2 indicate surfaces corresponding to the canonical Wnt-binding interfaces of Frizzled receptors. b) AlphaFold3 model of RL1-1 (teal) in complex with ROR1, predicting displacement of the α5 helix and exposure of the underlying hydrophobic pseudo-site 1. c) Comparison of the α1, α4 and α5 regions of ROR1 and ROR2. Sequence differences are highlighted, including replacement of the ROR2 α1 arginine involved in an α1– α5 interaction by glutamine in ROR1. d) Predicted interaction of RL1-1 with ROR1 pseudo-site 1. Enlarged views show contacts involving residues within the evolved ubiquitin loop; predicted interactions are indicated by dashed lines. e) Predicted RL1-1–ROR2 complex, in which the evolved loop instead engages a shallower surface near pseudo-site 2. The enlarged view highlights predicted contacts involving aromatic residues of the evolved loop.

Modeling of RL1-1in complex with ROR1 predicted displacement of α5 and opening of the underlying hydrophobic cavity, allowing the engineered ubiquitin loop to engage the exposed surface (Fig. 3b and 3d). Several residues within the evolved loop, including W9 and V16, were positioned at the predicted interface with α5, while additional aromatic and hydrophobic residues contacted the α1/α2 surface. Subtle sequence differences between ROR1 and ROR2 were observed in this region (Fig. 3c). Notably, α1 Arg205 in ROR2, which forms a hydrogen bond with the α5 unit closing pseudo-site 1, is replaced by a glutamine and potentially disrupts this interaction.

Modeling of RL1-1 with ROR2 produced a distinct predicted interaction in which the engineered loop engaged a shallower surface in the region of Wnt binding (pseudo-site 2; Fig. 3a), without equivalent opening of the α5-occluded cavity (Fig. 3e). This alternative model is consistent with the weaker and more rapidly dissociating ROR2 interactions observed by SPR and suggests that differential accessibility of the α5-gated hydrophobic surface may contribute to ROR1 preference.

### ROR1-binding U-bodies suppress wound closure in ROR1-expressing cells

ROR1 has been implicated in the regulation of tumor-cell motility through noncanonical Wnt signaling, providing a means for evaluating the cellular activity of the selected U-bodies using a wound-healing assay (Fig. 4a).^39, 40^ ROR1-expressing MDA-MB-231 cells were serum-starved and maintained in 1% FBS during the assay to limit the contribution of proliferation. Representative U-bodies were selected for testing based on a combination of ROR1-binding affinity and recombinant expression yield. Cells were then treated with representative RiPLE-derived ROR1 U-bodies at 1 µM or with PBS vehicle control, and wound closure was monitored over 24 h (Fig. 4b). Representative images showed progressive wound closure in vehicle-treated cells, whereas treatment with RL1-1, RL1-2, RL1-4, or RL1-5 visibly delayed closure over the same period. Quantification of the wound closure area confirmed this effect, with each of the tested RiPLE U-bodies reducing normalized wound closure to approximately half that of the vehicle-treated control at 24 h (Fig. 4c).

**Figure 4.**
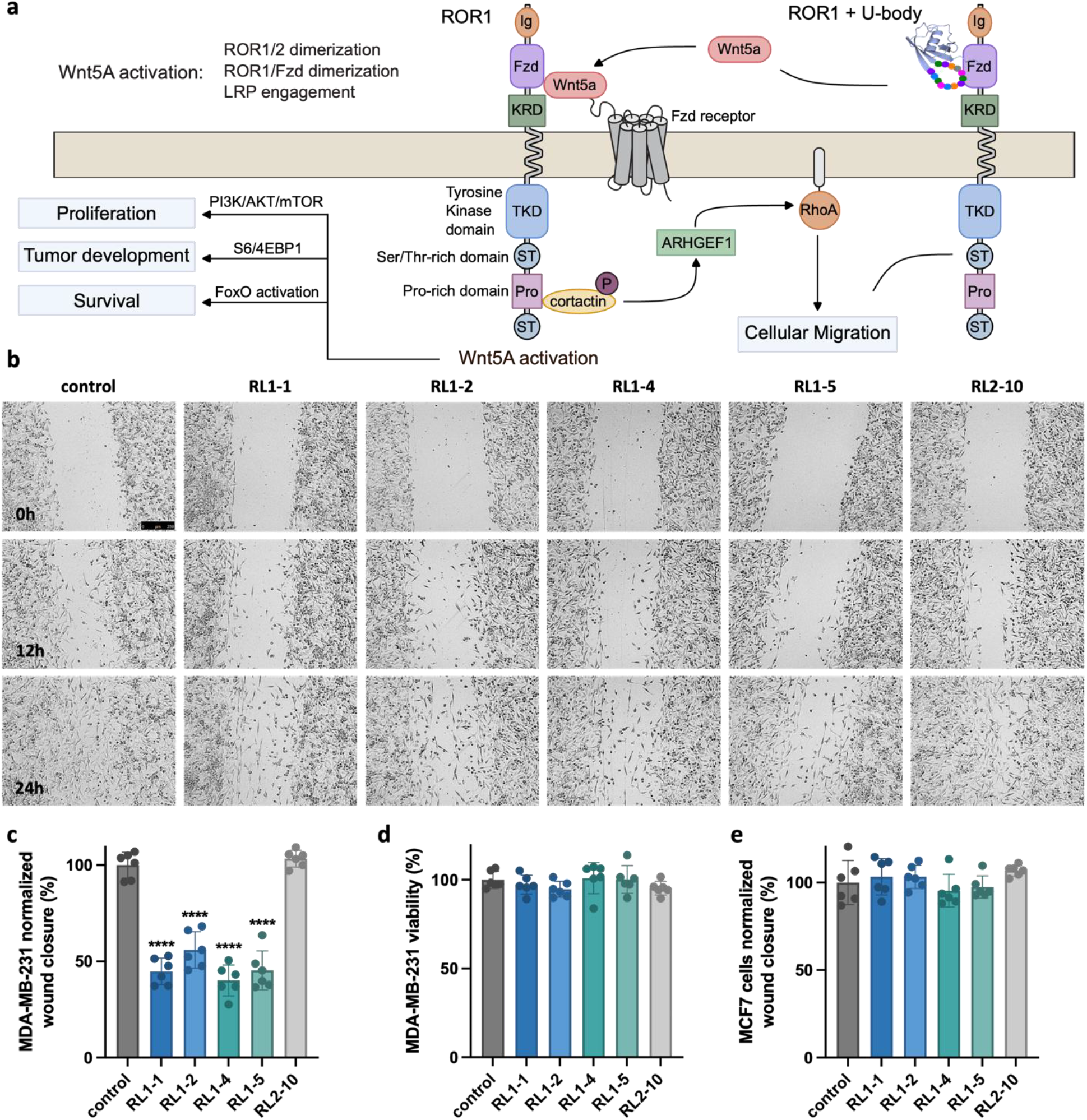
ROR1-binding U-bodies suppress wound closure in ROR1-expressing cells. a) Schematic representation of ROR1-associated Wnt5a signaling implicated in tumor-cell motility and the rationale for evaluating ROR1-binding U-bodies in a wound-healing assay. b) Representative bright-field images of MDA-MB-231 cells at 0, 12 and 24 h following treatment with PBS vehicle control, the ROR1-binding RiPLE U-bodies RL1-1, RL1-2, RL1-4 or RL1-5, or the unrelated GαS-binding U-body RL2-10. U-bodies were used at 1 µM and cells were maintained in 1% FBS during the assay. Contrast and sharpness were adjusted uniformly across images for visual clarity. c) Quantification of MDA-MB-231 wound closure at 24 h, normalized to the vehicle-treated control. d) Relative viability of MDA-MB-231 cells measured at the end of the 24-h assay. e) Quantification of wound closure in MCF7 cells after 48 h under the same treatment conditions, normalized to the vehicle-treated control. Data are presented as mean ± s.d. from n = 6 wells. Statistical significance was assessed by one-way ANOVA followed by Dunnett’s multiple-comparisons test, with each treatment compared with the vehicle control; ****(P < 0.0001). Scale bar, 250 µm.

To determine whether this response was associated with ROR1 recognition rather than a nonspecific effect of the ubiquitin scaffold, the same ROR1-binding U-bodies were evaluated in MCF7 cells, which express little ROR1.^41^ No significant reduction in wound closure was observed across all compounds when compared to the vehicle-control (Fig. 4e). The unrelated GαS-binding U-body, Z2-10, was included as an additional control and produced no detectable effect on wound closure in either MDA-MB-231 or MCF7 cells. Cell viability measured at the end of the assay was also not significantly reduced by any of the tested compounds, indicating that the impaired wound closure observed in MDA-MB-231 cells was not attributable to cytotoxicity (Fig. 4d).

Lasso-grafted ROR1 U-bodies showed a similar phenotype, reducing wound closure in MDA-MB-231 cells to a comparable extend, while having no significant effect in MCF7 cells compared to controls (SI Fig. S8-S9). Together, these results show that ROR1-binding U-bodies discovered by both RiPLE and lasso-grafting suppress wound closure under low-serum conditions preferentially in ROR1-expressing cells, without detectable effects on cell viability.

## Discussion and Conclusion

This study establishes RiPLE as a complementary approach to peptide-guided lasso-grafting (LG) for engineering protein binders by mRNA display. Where LG begins with an established peptide pharmacophore and adapts it to a protein scaffold, RiPLE allows the binding sequence itself to emerge directly within the scaffold during selection. The identification of binders against ROR1 and GαS demonstrates that high-affinity recognition can be achieved without requiring prior peptide discovery and provides a direct comparison between these two modes of protein evolution.

The ROR1 and GαS selections further show that the evolutionary behavior of RiPLE is target dependent. RiPLE converged strongly toward a dominant sequence against ROR1 but retained substantially greater sequence diversity against GαS, whereas the corresponding lasso-graft selection converged to only a few families. Thus, the strong convergence observed for ROR1 does not appear to reflect an intrinsic limitation of the randomized-loop library. The LG selections against ROR1 also underwent substantial sequence adaptation after the parental peptide was incorporated into ubiquitin. Changes within the graft and linker regions suggest that selection can compensate for structural constraints introduced during peptide grafting. Notably, a proline residue used as a replacement for the source peptide cβAA was overwhelmingly replaced by a leucine. While engineered disulfide bridges have previously been used deliberately to stabilize engineered protein loops, the cysteine pairs observed here arose naturally during selection. Consistent with our previously reported U-body work, disulfide-constrained structures were not superior to other sequences.^29^ Together, these observations highlight a fundamental distinction between the two strategies: lasso-grafting evolves a scaffold around an imposed recognition element, while RiPLE allows the recognition element itself to evolve in its final binding protein surface.

This distinction also has practical implications for binder discovery. RiPLE removes the requirement for both an initial peptide-discovery campaign and subsequent graft optimization in favor of a straightforward selection process. As a single library can be used for any new protein target, it also alleviates the costs and uncertainty associated with bespoke library designs. LG remains attractive when a validated peptide ligand is available and its functional properties can be successfully transferred to a protein scaffold. In the case of ROR1 for instance, LG considerably enhanced the target-specificity of the source peptide, whereas RiPLE generally yielded stronger *de novo* binders. These approaches should therefore be considered complementary rather than competing, and where possible, pursued in parallel to maximize the likelihood of identifying potent U-bodies.

The ROR1 selections also revealed substantial discrimination between the closely related receptors ROR1 and ROR2. This preference was unexpected because previously identified ROR1-binding peptides, including the parental peptide used for lasso-grafting, bound ROR1 and ROR2 with comparable affinities. The U-bodies formed relatively long-lived complexes with ROR1 but dissociated much more rapidly from ROR2, indicating that selectivity can emerge through differences in complex stability despite the high similarity of the two receptors. The structural models suggest a possible explanation involving the α5-blocked hydrophobic cavity within the ROR cysteine-rich domain. In the predicted ROR1 complexes, displacement of α5 permits engagement of the hydrophobic pseudo-site 1 by the engineered ubiquitin loop, whereas the ROR2 models favor a shallower interaction into pseudo-site 2. This raises the possibility that the U-bodies exploit subtle conformational differences between the receptors that are not apparent from sequence conservation alone. The proposed binding mode remains to be validated experimentally, but it also suggests that these molecules may provide useful probes for investigating the mechanistic basis of how ROR cysteine-rich domains engage Wnt ligands, as well as other interaction partners.

The ROR1-binding U-bodies were also active in a cellular context, reducing wound closure in ROR1-expressing MDA-MB-231 cells while showing little effect in MCF7 cells with low ROR1 expression. The inactivity of an unrelated GαS-binding U-body and the absence of detectable cytotoxicity further support a target-associated effect rather than a nonspecific consequence of the ubiquitin scaffold. Nevertheless, scratch closure can reflect contributions from both migration and proliferation, and the present experiments do not define the downstream signaling events responsible for the phenotype. Structural and mechanistic characterization of the selected ROR1 binders could therefore provide useful tool compounds for dissecting ROR1/Wnt recognition and downstream signaling. In parallel, extending RiPLE to alternative scaffolds and loop architectures, potentially together with functional or state-selective selection strategies, may enable access to distinct binding modes, improved therapeutic properties, and new cellular functionalities

In conclusion, peptide-guided LG and RiPLE provide complementary routes for evolving engineered protein binders by mRNA display. By allowing a recognition surface to emerge de novo within the final protein scaffold, RiPLE expands and streamlines the use of ultralarge genetically encoded libraries beyond the optimization of pre-existing peptide pharmacophores.

## Methods

### mRNA Library preparation

DNA templates were assembled as previously described^29^ by overlap extension PCR from oligonucleotide primers (Eurofins Genomics) using KOD One polymerase mastermix (Toyobo, KMM-101) at 100 µL reaction scale. The full sequence design is shown in SI figure S1. The mixtures were denatured at 98 °C for 10s, followed by five cycles of annealing (65 °C for 5s) and extension (68 °C for 5s). In a second step, the extension products were mixed and subjected to another PCR amplification with KOD One (98 °C for 10s, 61 °C for 5 s, 68 °C for 5 s, seven cycles) The list of all oligonucleotides and mixing ratios can be found in Supplementary Table 2. PCR products were purified by phenol/chloroform extraction and ethanol precipitation. Transcription was carried out using the following mixtures: 40 mM Tris-HCl buffer (pH 8.0), 22.5 mM MgCl_2_, 10 mM dithiothreitol (DTT), 1mM spermidine, 0.01% Triton X-100, 3.75 mM NTP mix, 0,04 UµL^-1^ RNase inhibitor (Promega, N2615) and 120 mM T7 RNA polymerase at 37 °C for 16 h. Then, the DNA was digested using 30 µL of 1 UµL^-1^ RQ1 RNase-free DNAse (Promega) for 1 h. at 37 °C. The reaction was quenched using EDTA (f.c. 68 mM) and NaCl (f.c. 270 mM) after which the RNA was precipitated with isopropanol and purified using 8% denaturing polyacrylamide gel electrophoresis (PAGE) containing 6 M urea. The list of oligonucleotides and primers used is available in supplementary table 2.

### mRNA display affinity selections

Selections were carried out as previously described following the general protocol below, and adapted for each protein target as required. The random mRNA library was ligated at the 3’ end with a puromycin linker and precipitated with EtOH. The resuspended ligated library was then subjected to following *in vitro* translation mixture: 50mM HEPES-KOH (pH 7.6), 100 mM potassium acetate, 12.3 mM magnesium acetate, 2mM ATP, 2mM GTP, 1mM CTP, 1mM UTP, 20 mM creatine phosphate, 0.1 mM 10-formyl-5,6,7,8-tetrahydrofolic acid, 2mM spermidine, 1 mM DTT, 1.5 mg/mL *E. coli* total tRNA, 1.2 µM *E*.*coli* ribosomes, 0.6 µM methionyl-tRNA formyltransferase, 2.7 µM IF1, 0.4 µM IF2, 1.5 µM IF3, 0.26 µM EF-G, 10 µM EF-Tu, 0.66 µM EF-Ts, 0.25 µM RF2, 0.17 µM RF3, 0.5 µM RRF, 4 µg/mL creatine kinase, 3 µg/mL myokinase, 0.1 µM inorganic pyrophosphatase, 0.1 µM nucleotide diphosphate kinase, 0.1 µM T7 RNA polymerase, 0.73 µM AlaRS, 0.03 µM ArgRS, 0.38 µM AsnRS, 0.13 µM AspRS, 0.02 µM CysRS, 0.06 µM GlnRS, 0.23 µM GluRS, 0.09 µM GlyRS, 0.02 µM HisRS, 0.40 µM IleRS, 0.04 µM LeuRS, 0.11 µM LysRS, 0.03 µM MetRS, 0.068 µM PheRS, 0.16 µM ProRS, 0.04 µM SerRS, 0.09 µM ThrRS, 0.03 µM TrpRS, 0.02 µM TyrRS, 0.02 µM ValRS, 0.5 mM each of the 20 proteinogenic amino-acids, and 1.5 µM of the puromycin-ligated mRNA library. Translation was performed for 45 min at 37 °C in 50 µL of translation system (first round) or 5 µL (round 2 onwards). The translated library was then incubated at 25 °C for 7 min to allow for conjugation of the U-bodies with the corresponding mRNA-puromycin, followed by addition of 0.5 µL of 500 mM EDTA (pH 8.0) and further incubation at 37 °C for 5 min to dissociate ribosomes from the mRNA-U-body conjugates. Reverse transcription was carried out using RT-primer (see Supplementary Table 2) and M-MLV reverse transcriptase lacking RNase H activity (Promega, M3682). The resulting cDNA/mRNA/U-body conjugates were subjected to negative selections with a 1:1 mixture of naked protein G Dynabeads and Fc-covered beads. For hINSR, the negative selections were performed instead with a 1:1 mixture of naked streptavidin Dynabeads and biotin-covered beads.. The supernatant from the negative selection was then transferred to a new sample of negative beads. Note that negative selections were not conducted in the first round to preserve library diversity. The supernatant from the last negative sample was then transferred to Dynabeads with immobilized target protein. The beads were washed thrice with ice cold selection buffer, then suspended in 100 µL water and eluted at 95 °C for 5 min. Then, 100 uL of 2X KOD One polymerase mastermix was added along with 0.25 µM each of forward and reverse primers (T7G10MUb6.F63 and SSG2an13.R36 respectively). The recovery rate of cDNA was measured by qPCR using 1 µL of eluted library and 19 µL of 1X PCR buffer containing SYBR Green 1 (Lonza, 50513) and KOD One polymerase. The elute was amplified by PCR to create a cDNA library, which was then transcribed and submitted to another round of selection.

#### Selection buffers and protein immobilization

ROR1 selections were performed with TBS-T (Tris-HCl pH 8.0, 50 mM,150 mM NaCl, 0.05% (v/v) Tween-20). GαS selections were performed with HBSMgT (25 mM HEPES pH 7.5, 150 mM NaCl, 1mM MgCl2, 0.05% (v/v) Tween-20) supplemented with 0.5 mM GppNHP. Fc-tagged ROR1 (Abcam, ab270585) was immobilized on Dynabeads Protein G (Invitrogen, 10009D). Biotinylated GαS (gift from Prof. K. Shokat) was immobilized on Dynabeads Streptavidin M280 (Invitrogen, 11206D).

#### Next generation sequencing

The pooled cDNA libraries were PCR-amplified with KOD One PCR Master Mix (using 0.5 µM Rd1T7g10M.F70 and an13Rd2.R49 as primers according to the manufacturer’s instructions. The products were carried forward to the second PCR step, using KOD One PCR Master Mix and Nextera XT v2 Set (sequences from Illumina) primers to install sequencing barcodes on each sample. Then, PCR products were combined, extracted by phenol/chloroform, and column-purified using a NucleoSpin kit (TaKaRa) according to the manufacturer’s instructions. The concentration of the combined cDNA sample was measured with Qubit (Thermo Fisher) using the dsDNA HS kit. The resulting DNA was appropriately diluted, denatured, and combined with PhiX Control v3 (Illumina). The denatured DNA was sequenced on an Illumina MiSeq instrument. The collected fastq data was analyzed using Prof. A. Vinogradov’s combinatorial library analysis suite (https://github.com/avngrdv/clibas). The top 100 sequences from each sequenced pool are shown in Supplementary Table 1.

### Protein Expression and Purification

U-body constructs were prepared as previously desribed.^29^ U-body sequences were cloned into the pET22b(+) vector using InFusion Snap Assembly Master Mix (Takara Bio, 638947) and transformed in *E. coli* NEB Turbo (New England Biolabs, C2984). The purified plasmids were sent to FASMAC for validation of the cloned sequence using Sanger sequencing and primer NT0071. U-bodies were expressed with a N-terminal His6-tag. from the prepared plasmids in *E. coli* BL21-gold (DE3) (Agilent, 230132), see Supplementary Figure 3 for the detailed expression construct. Cultures were grown to OD_600_ 0.4 at 37 °C in Lysogeny Broth medium (LB Miller), containing 100 µg/mL ampicillin, and expression was induced with 0.4 mM IPTG at 16 °C for 24 h. Cells were lysed by sonication in lysis buffer (50 mM Tris-HCl, 150 mM NaCl, 20 mM imidazole), and purified by Ni^2+^ Sepharose^TM^ 6 Fast Flow (Cytiva, 17531802). The U-bodies were eluted from the Sepharose bead with an elution buffer (50 mM Tris-HCl, 150 mM NaCl, 500 mM imidazole) and concentrated by ultra-filtration. The U-bodies were used without further purification for SPR analysis. For additional assays, selected target-binding U-bodies were further purified by size exclusion chromatography using Superdex 75 increase 10/300 GL (Cytiva) or HiLoad 16/600 Superdex 200 pg (Cytiva) column on ÄKTA avant 25 (Cytiva) or ÄKTA pure (Cytiva) system (running buffer: 50 mM sodium citrate pH 5.5, 50 mM NaCl). Correct expression and purity was assessed by tricine SDS-PAGE before & after SEC (SI Fig. S3).

### Surface-Plasmon Resonance

The dissociation constant and the binding kinetic constants of each U-body construct was determined by SPR using a Biacore 8K instrument (Cytiva) at 25 °C. ROR1 and ROR2 were immobilized on a protein G sensor chip, and the running buffer was PBS-T. GαS was immobilized on a Biotin CAPture chip and the running buffer was HBSMg-T. The obtained binding sensorgrams were analyzed using Biacore insight evaluation software and fitted by the standard 1:1 binding model (See Supplementary Figures S5-S7).

### Serum stability

The U-body sample (6 μM) and a serum-resistant internal standard peptide consisting only of D-amino acids (NH2-PEG5-wstndwstnd-PEG5-CONH2, 6 μM) were mixed and incubated in human serum (Cosmo Bio, 12181201) at 37 °C for up to 72h. At each time point, 10 μL of the mixture was removed and quenched by adding 23 μL of methanol, followed by incubation on ice for 15 min. Following centrifugation (15,000 × g, 25 °C, 5 min), the supernatant was analyzed using LC/MS employing a reverse-phase column (ACQUITY UPLC BEH C4, 2.1 × 150 mm; Waters) and a Xevo G2-XS QTof system (Waters) with a linear gradient from 1% B to 61% B. Buffer A: water with 0.1% (v/v) formic acid; buffer B: acetonitrile with 0.1% (v/v) formic acid. The percentages of remaining peptides were determined by using the peak area integration of the chromatograms. The obtained LC/MS data were analyzed using a MassLynx 4.1 (Waters).

### Thermal shift assay

The thermal stability of U-bodies was measured by differential scanning fluorimetry as described previously.^29^ U-bodies solutions were prepared at 4 μM in TBS-T in a 96-well PCR plate and mixed with SYPRO Orange at a final concentration of 1:5000 per the manufacturer’s recommendation. The fluorescence intensity was monitored during a temperature scan ranging from 37 °C to 97 °C at a 0.03 °C/s gradient using a real-time PCR instrument. No melting transition could be detected in the measured range, consistent with prior literature.^29^

#### Cell Culture

MDA-MB-231 (ECACC, EC92020424-F0) and MCF-7 (ATCC, HTB-22) cells were cultured in Dulbecco’s Modified Eagle Medium (DMEM; Nacalai Tesque, 08458-45) supplemented with 10% FBS, 100 units/mL penicillin, 0.1 mg/mL streptomycin, and 0.25 µg/mL amphotericin B. All cells were maintained in a humidified incubator at 37°C with 5% CO2 in air.

### Wound healing assay

MDA-MB-231 or MCF7 cells were seeded in 96-well plates at 4.0 x 10^4^ cells per well in 100 µl DMEM supplemented with 10% FBS and 1% penicillin-streptomycin. The following day, the medium was replaced with DMEM containing 1% FBS and 1% penicillin-streptomycin, and cells were serum-starved overnight. Once a confluent monolayer was obtained, a scratch wound was generated using a sterile wooden toothpick. The medium was removed and the cells were washed with PBS to remove detached cells. U-bodies were diluted to a final concentration of 1 µM in DMEM containing 1% FBS and 1% penicillin–streptomycin, with PBS adjusted to 1% (v/v) in all treatment conditions. A total volume of 100 µl was added to each well. Cells were incubated at 37 °C for 1 h before acquisition of the initial image (t=0) using a Leica DMI6000B inverted microscope. Wounds were subsequently imaged at 12 and 24 h for MDA-MB-231 cells and at 24 and 48 h for MCF7 cells. Wound area was quantified using a custom macro in ImageJ (see below). For each time series, the analysis region was adjusted to encompass the wound consistently across images. Wound closure for each well was calculated relative to its corresponding t=0 wound area. For comparisons between experiments, wound closure values were normalized to the corresponding vehicle control within each experiment. Two independent experiments were performed, each with three replicate wells per condition (six wells per condition in total). Data are presented as mean ± s.d. Statistical analysis was performed using GraphPad Prism 10 for macOS. Group differences were evaluated by one-way ANOVA followed by Dunnett’s multiple-comparisons test, with each treatment compared with the vehicle control.

### Macro

~~~
run(“Gaussian Blur…”, “sigma=0.5 stack”);
run(“Find Edges”, “stack”);
setOption(“BlackBackground”, true);
run(“Convert to Mask”, “method=Otsu background=Dark”);
run(“Invert”, “stack”);
run(“Dilate”, “stack”);
run(“Fill Holes”, “stack”);
run(“Measure”);
~~~

### Cell viability assay

Cell viability was assessed immediately following acquisition of the final wound-healing images using a resazurin reduction assay. 10 µL of resazurin was added to each well and cells were incubated for 45 min at 37 °C. Resazurin reduction was quantified by measuring the fluorescence with excitation at 544 nm and emission at 590 nm using a Tecan M1000 Pro microplate reader (Tecan group) with optimal gain settings, 20 µs integration time and 400 Hz flash frequency. Background signal from wells containing medium and resazurin without cells was subtracted. Viability was normalized to the corresponding vehicle-treated control within each experiment. Data are presented as mean ± s.d. Statistical significance was assessed by one-way ANOVA followed by Dunnett’s multiple-comparisons test, with each treatment compared with the vehicle control.

## Data availability

Data supporting the reported findings is available in the manuscript and in supplementary information files. Source data and detailed sequences for plasmids, vectors and mRNA constructs is available upon reasonable request.

## Code availability

The NGS analysis was performed using Prof. A. Vinogradov’s combinatorial library analysis suite, which is available online on Github (https://github.com/avngrdv/clibas).

## Acknowledgements

We thank Michelle Freyer for technical assistance in expressing some U-body samples. T.K. and G.P. received postdoctoral fellowship funding from the Japan Society for the Promotion of Science (JSPS). K.C. received funding from the JSPS Grant-in-Aid for Early Career Researchers (26K17966) and AMED UTOPIA Next-Generation Diagnostic Tool Discovery Program (JP223fa627001). H.S received funding from the JSPS Grant-in-Aid for Specially Promoted Research (JP20H05618).

## Author Contributions

K.C. and H.S. designed the concept and experiments. K.C., T.K. and G.P. performed the selections, protein expression, assays and all downstream analysis. K.C. designed and prepared the mRNA libraries RL, LG1 and LG2; G.P. designed and prepared library LG3. T.K. performed the cell studies. K.C., T.K. and H.S. wrote the manuscript. All authors contributed input to the manuscript.

## Additional information

Supplementary information is available, including detailed NGS data for all selections, a list of primers used, detailed mRNA sequence of the RiPLE library, SPR sensorgrams, electrophoresis gels of expressed U-bodies, and results from thermal shift assays, serum stability assays, and wound healing assays.

