## Supplementary Information for "Random *in vitro* Protein single-Loop Engineering (RiPLE) by mRNA display"

7-3-1 Hongo, Bunkyo-ku, Tokyo 113-0033, Japan

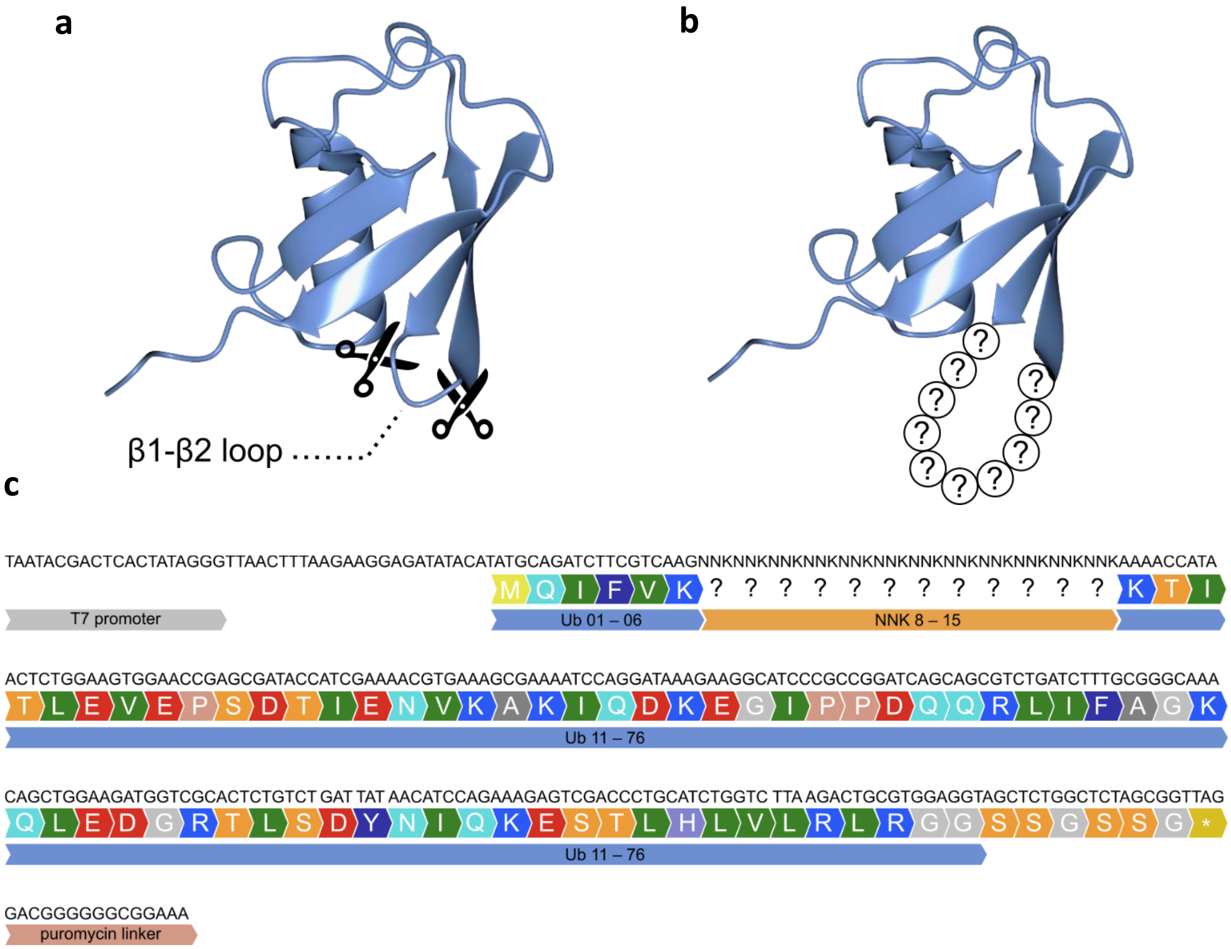


**Figure S1.** Library design for mRNA display selection. a) The grafting is performed at the b1-b2-turn (“loop 1”). B) In the case of RiPLE, the native loop 1 (Ub8-10) is replaced by 8-15 randomized residues. C) Full sequence of the mRNA display library including untranslated regions at the 3’ and 5’ ends.

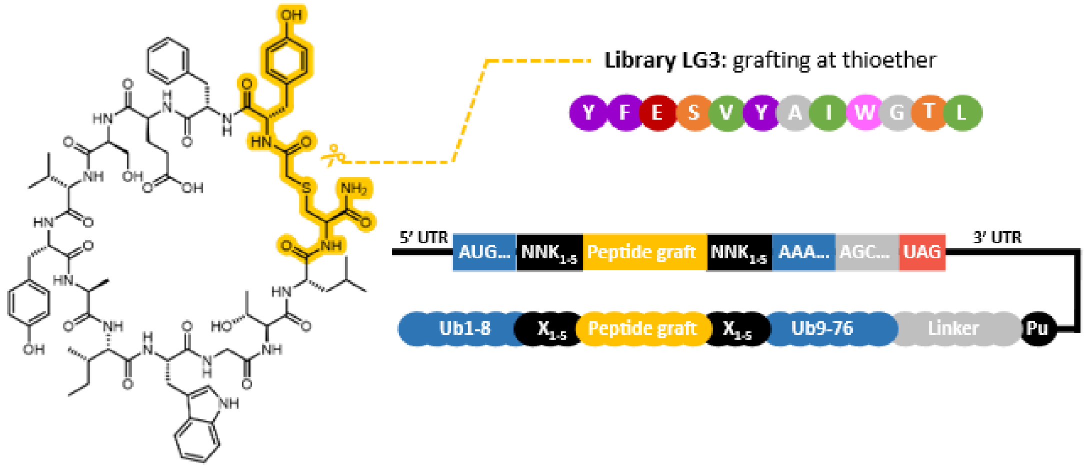


**Figure S2.** left: structure of the source peptide GN13 with the thioether macrocyclization highlighted. Right: Grafting was performed at the thioether junction, replacing *D*-Tyr with *L*-Tyr and deleting Cys.

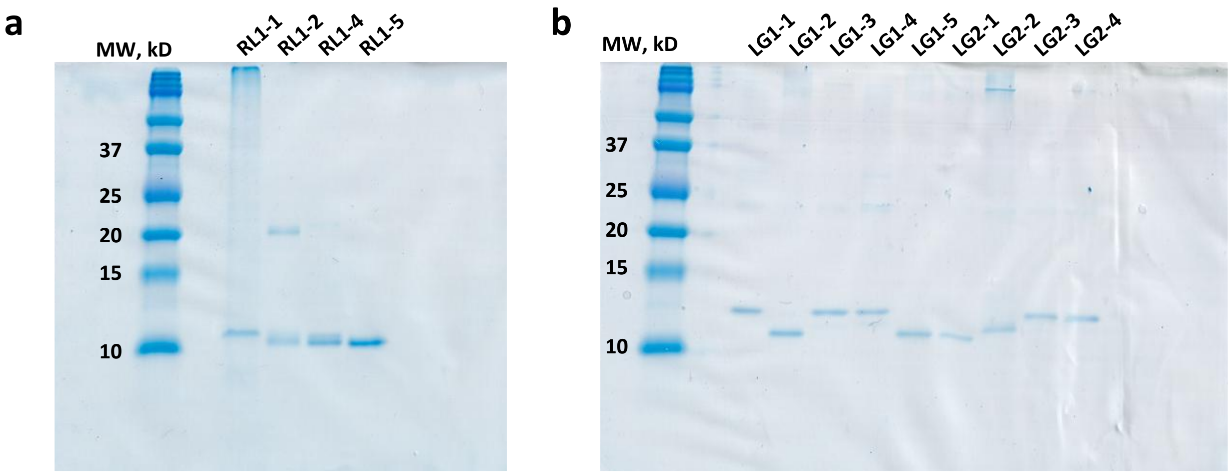


**Figure S3:** Tricine SDS–PAGE analysis of SEC-purified ROR1-binding U-bodies. a) RiPLE-derived variants RL1-1, RL1-2, RL1-4 and RL1-5. b) Lasso-grafted variants LG1-1–LG1-5 and LG2-1–LG2-4. Molecular-weight markers are indicated in kDa. Note that RL1-3 expresses poorly and was used without SEC purification for SPR analysis, and subsequently excluded from cell-assays. Gels were stained with Coomassie Brilliant Blue following size-exclusion chromatography.

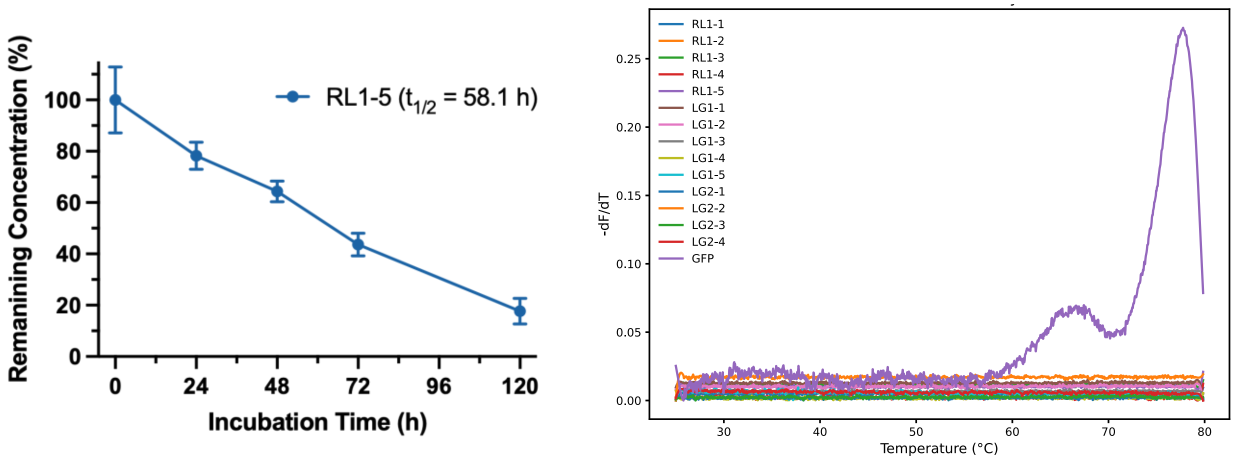


**Figure S4:** Serum stability of RL1-5 (left) and thermal shift-assays of all anti-ROR1 U-bodies (right). A sample of in-house expressed, crude GFP was used as a control.

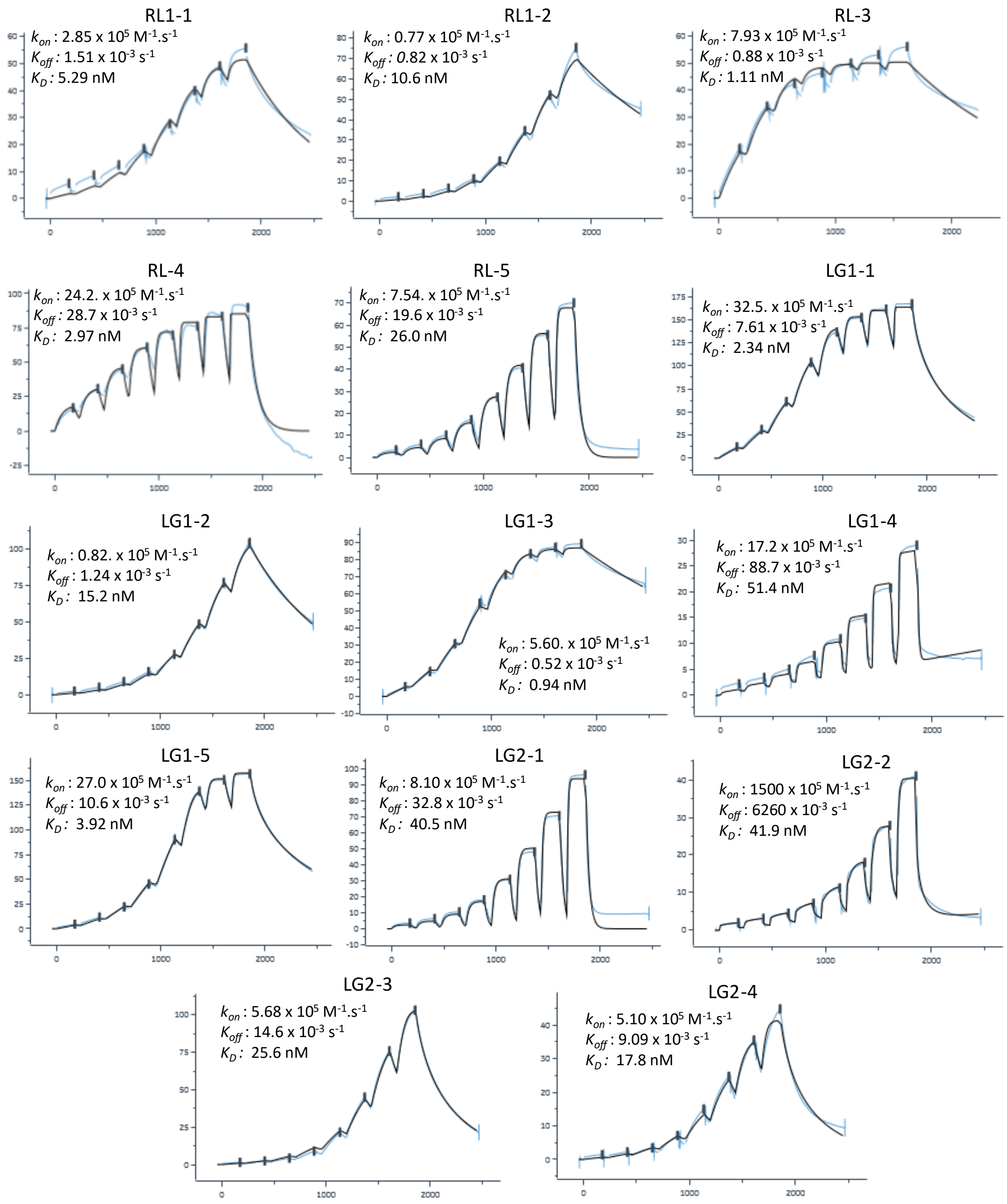


**Figure S5.** Sensorgrams of U-bodies tested against ROR1 with their kinetic data when allowed by response/fitting. Measurements were performed on a series of 2-fold dilution from 400 nM to 3.1 nM.

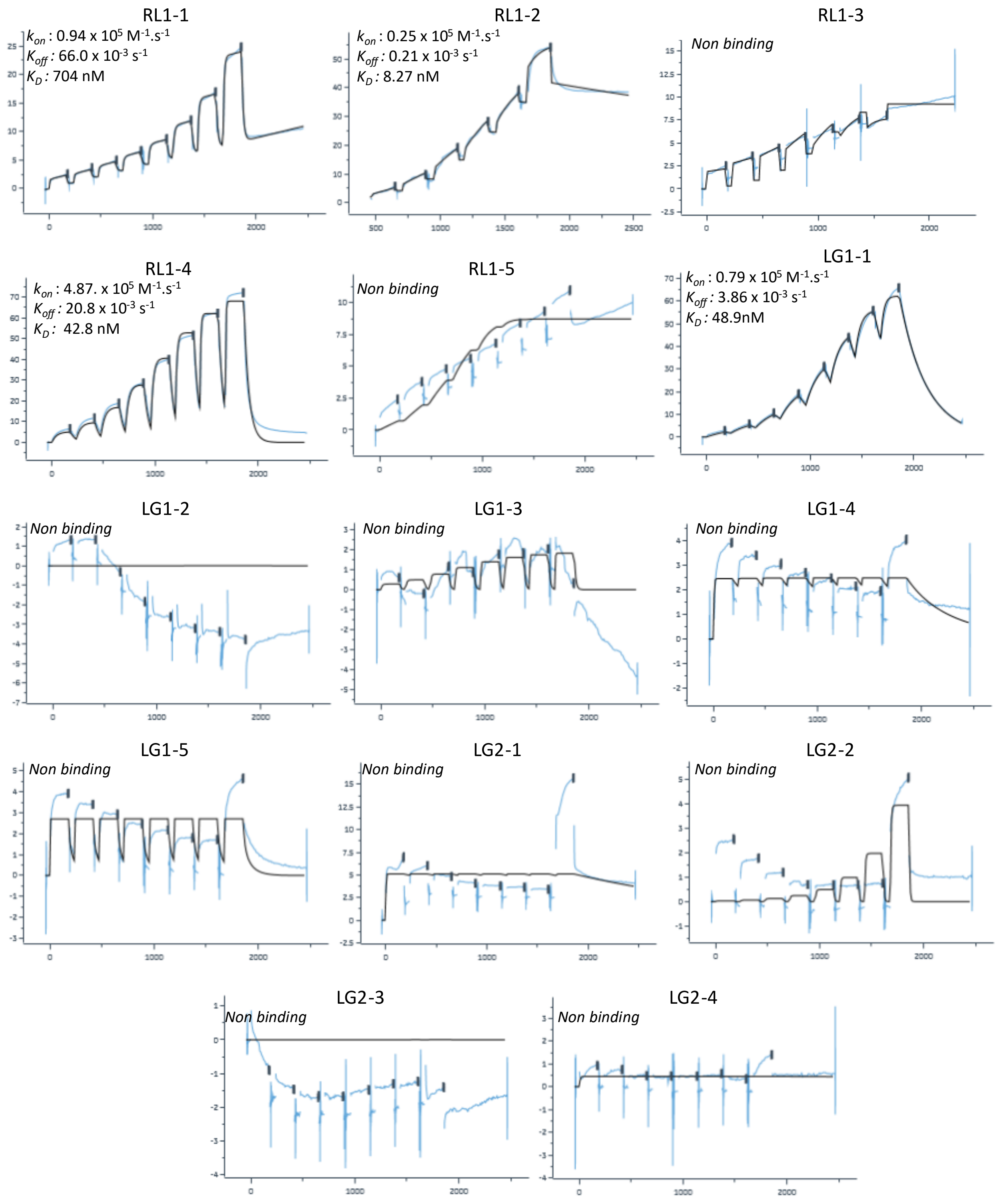


**Figure S6.** Sensorgrams of U-bodies tested against ROR2 with their kinetic data when allowed by response/fitting. Measurements were performed on a series of 2-fold dilution from 400 nM to 3.1 nM.

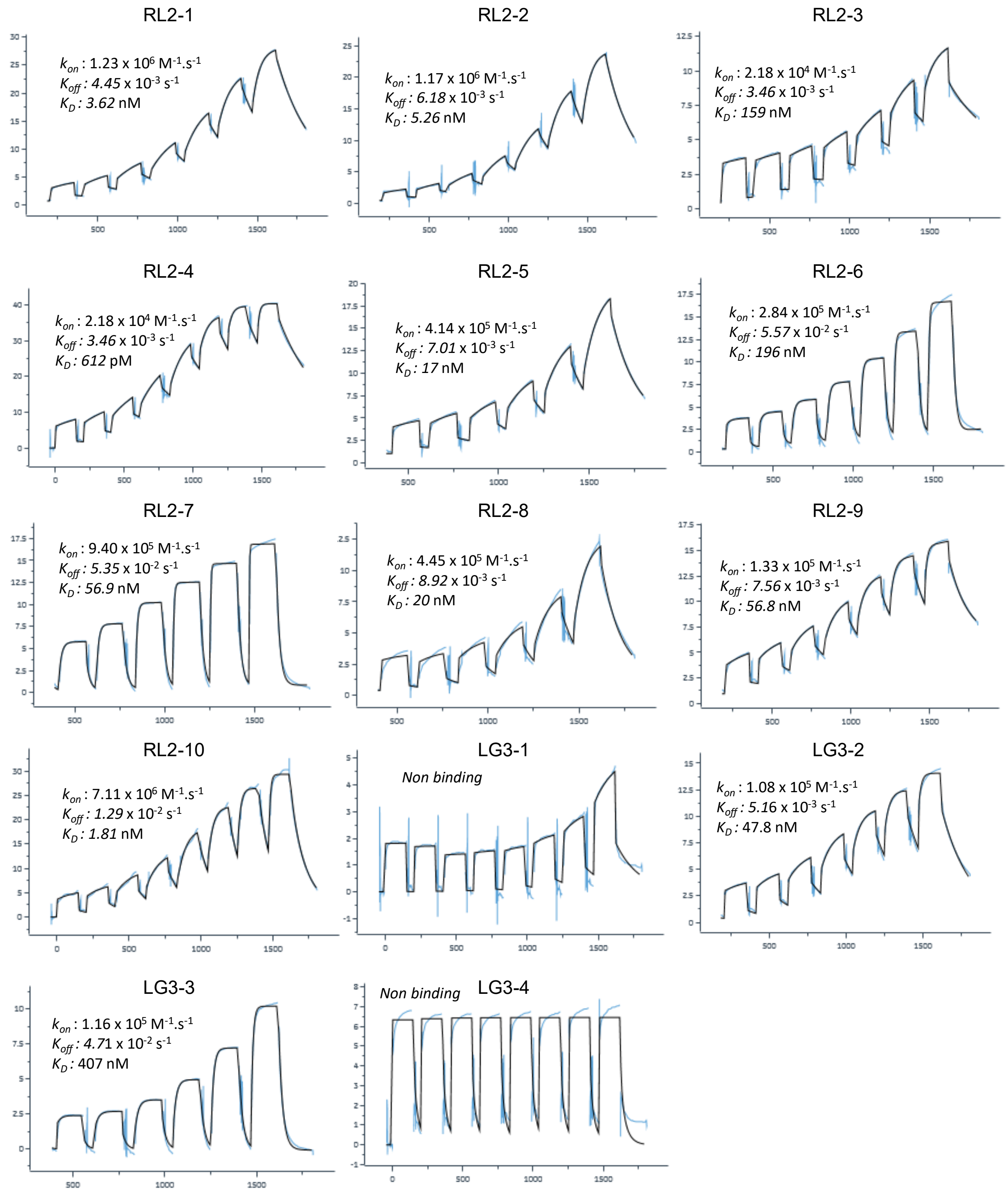


**Figure S7.** Sensorgrams of U-bodies tested against GppNhp-loaded GαS(R201C) with their kinetic data when allowed by response/fitting. Measurements were performed on a series of 2-fold dilution from 500 nM to 3.9 nM, and for strong binders (ZS2-1, ZS2-2, ZS2-4, ZS2-5, ZS2-8 and ZS2-10) on a series of 2-fold dilution from 15 nM to 0.12 nM.

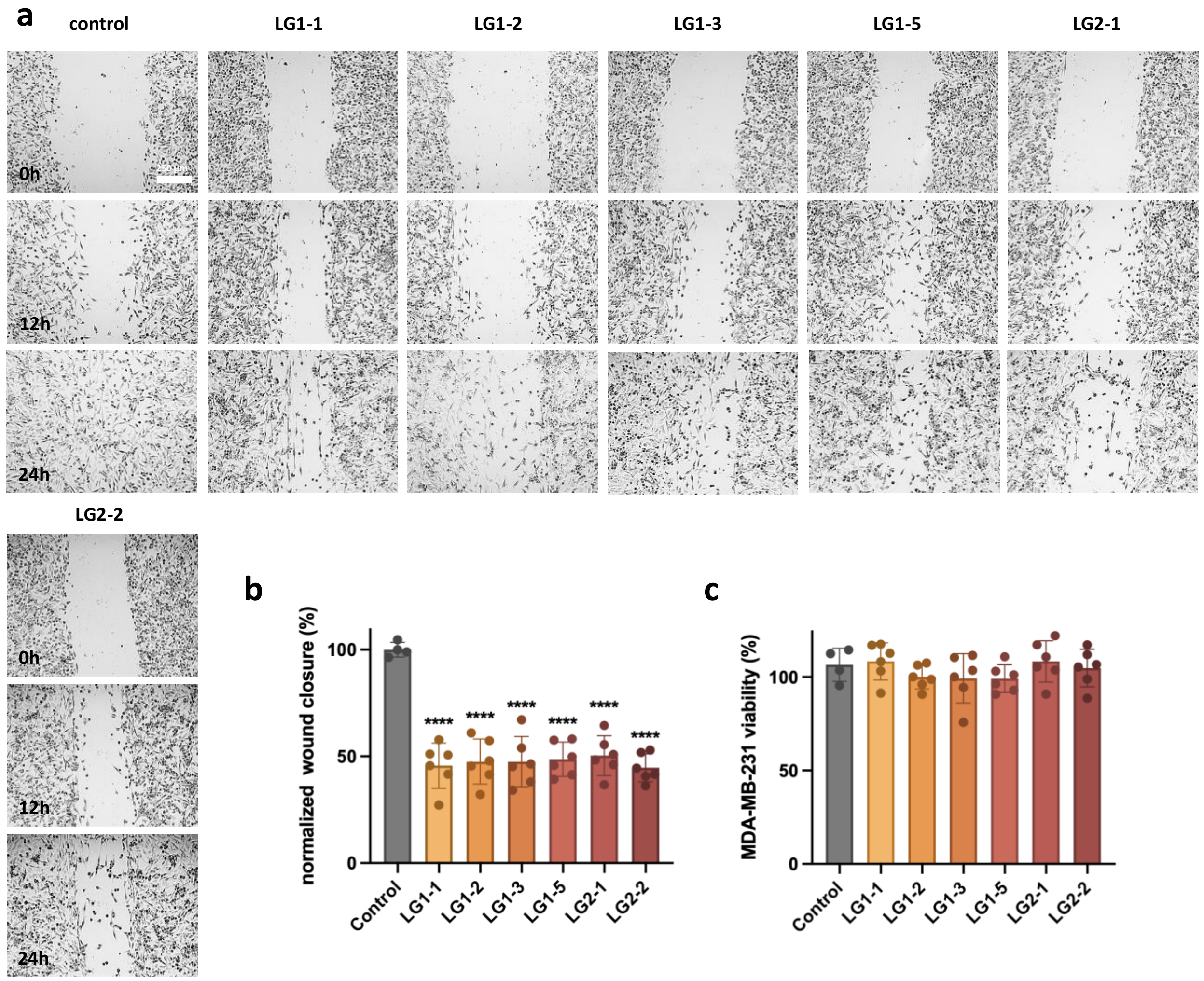


**Figure S8.** Lasso-grafted ROR1 U-bodies suppress wound closure in MDA-MB-231 cells. a) Representative images of MDA-MB-231 cells at 0, 12 and 24 h following treatment with PBS vehicle control or the indicated ROR1-binding lasso-grafted U-bodies (LG1-1, LG1-2, LG1-3, LG1-5, LG2-1 and LG2-2). U-bodies were used at 1 μM and cells were maintained in 1% FBS during the assay. Scale bar, 250 μm. Contrast and sharpness were adjusted uniformly across images for visual clarity. b) Quantification of wound closure at 24 h, normalized to the vehicle-treated control. c) Relative viability of MDA-MB-231 cells measured by resazurin reduction following completion of the 24-h wound-healing assay. Data are presented as mean ± s.d. Statistical significance was assessed by one-way ANOVA followed by Dunnett's multiple-comparisons test, with each treatment compared with the vehicle control; $****P<0.0001$.

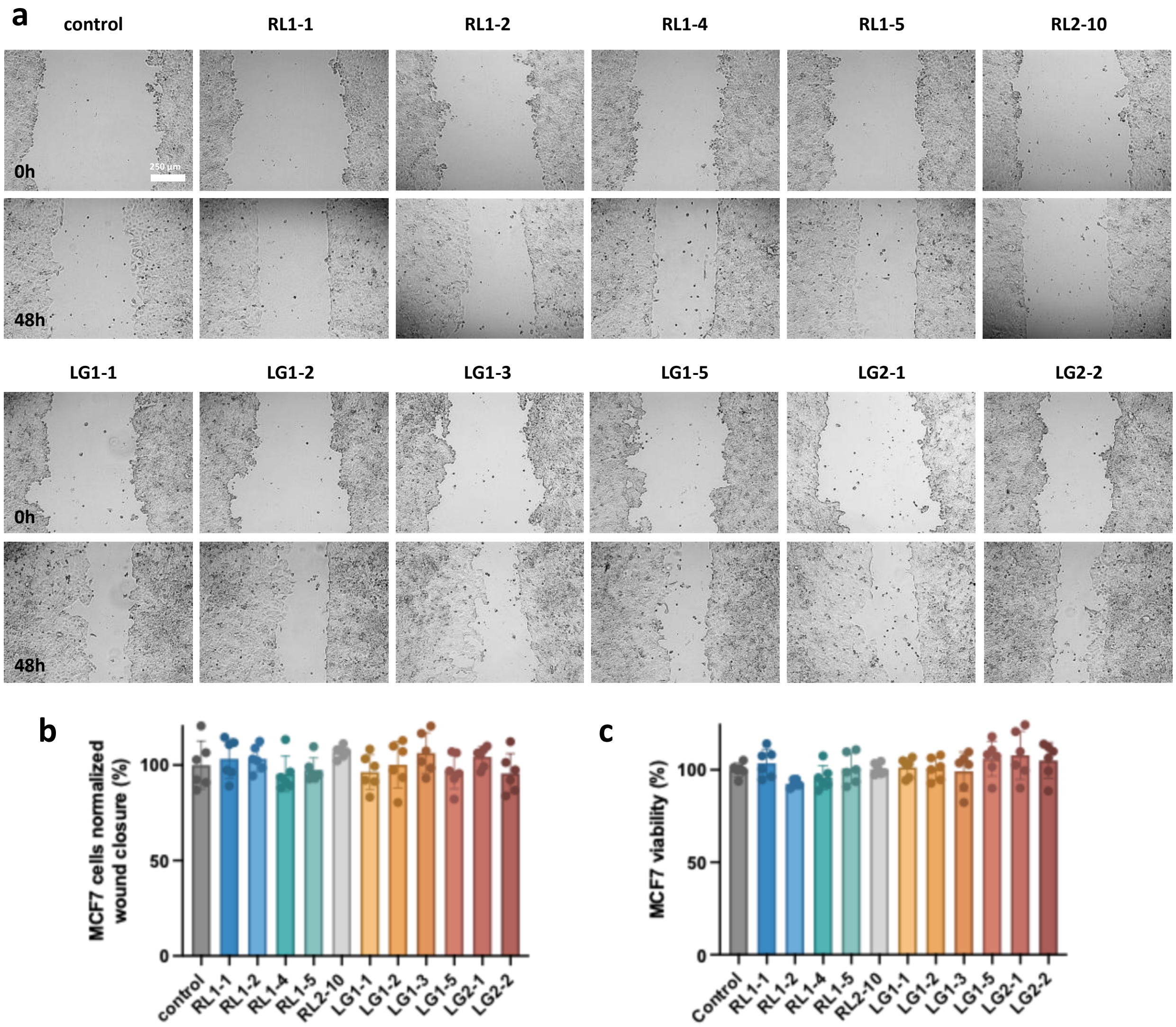


**Figure S9.** ROR1-binding U-bodies show little effect on wound closure in MCF7 cells. a) Representative wound-healing images of MCF7 cells at 0 and 48 h following treatment with PBS vehicle control, the indicated RiPLE-derived ROR1 U-bodies (RL1-1, RL1-2, RL1-4 and RL1-5), lasso-grafted ROR1 U-bodies (LG1-1, LG1-2, LG1-3, LG1-5, LG2-1 and LG2-2), or the unrelated GαS-binding U-body RL2-10. U-bodies were used at 1 µM and cells were maintained in 1% FBS throughout the assay. Scale bar, 250 µm. Contrast and sharpness were adjusted uniformly across images for visual clarity. b) Quantification of wound closure at 48 h, normalized to the vehicle-treated control. c) Relative viability of MCF7 cells measured by resazurin reduction following completion of the 48-h wound-healing assay. Data are presented as mean ± s.d.
