## Supplementary Table 1 for "Random *in vitro* Protein single-Loop Engineering (RiPLE) by mRNA display"

### Supporting Information Table S1

#### Page 1 - ZeroStart library against ROR1

Top 100 sequences from ZeroStart library against ROR 1 in different selection rounds, with NGS read frequency. Bold entries indicate the sequences selected for further analysis.
The Ubiquitin core residues around the randomized loop 1 insert are ommitted for clarity. They span Ub1-6 and Ub11-76

Note that RL1-6 could not be expressed and was not analyzed further

**Round 3**

| Rank | Loop 1 | read count | read frequency | U-body name |
|---|---|---|---|---|
| **1** | **LVWFRFYIFVY** | **36593** | **75.11%** | **RL1-1** |
| **2** | **FYLHGAVPSLFCLHY** | **6003** | **12.32%** | **RL1-2** |
| **3** | **ALLWCNVCECFW** | **1265** | **2.60%** | **RL1-3** |
| **4** | **WCYFCGLGRFVSSWW** | **716** | **1.47%** | **RL1-5** |
| 5 | YILFIWSGVRFLS | 448 | 0.92% | |
| 6 | SLWYFVWLTSRFST | 384 | 0.79% | |
| **7** | **LHIRWYDSLYLWL** | **372** | **0.76%** | **RL1-4** |
| 8 | VCMRWSCRAFSF | 254 | 0.52% | |
| **9** | **RGLVPWSLR** | **246** | **0.50%** | **RL1-6** |
| 10 | RGRSKCVFLLVCL | 228 | 0.47% | |
| 11 | SFVLLPWWGLHQL | 175 | 0.36% | |
| 12 | LYVPWWITFLRL | 162 | 0.33% | |
| 13 | VCVKWTCAILDF | 146 | 0.30% | |
| 14 | LVLFSMWWVVHF | 144 | 0.30% | |
| 15 | CIILWYFLIS | 114 | 0.23% | |
| 16 | LLVQWWLTYLVY | 113 | 0.23% | |
| 17 | IWWFYSYPHLGPSEL | 98 | 0.20% | |
| 18 | RGLLLWSSR | 80 | 0.16% | |
| 19 | RGLVAWSLR | 70 | 0.14% | |
| 20 | LVLSWWMTLLWF | 65 | 0.13% | |
| 21 | KLWCASCLIWW | 55 | 0.11% | |
| 22 | VGFGFFLPFGYFLF | 52 | 0.11% | |
| 23 | VVLYWYAVFFIWY | 47 | 0.10% | |
| 24 | YAIWWLGLFLLFF | 46 | 0.09% | |
| 25 | ASWRWFSSFFLDL | 44 | 0.09% | |
| 26 | IYWPHCAVLFFHCWA | 38 | 0.08% | |
| 27 | ICMLWNCWSVTF | 35 | 0.07% | |
| 28 | LCLLWTCRSYSW | 34 | 0.07% | |
| 29 | LILRWFSFFVILL | 34 | 0.07% | |
| 30 | SVFRWYASSWLHL | 32 | 0.07% | |
| 31 | LVMRWLLELLHHY | 29 | 0.06% | |
| 32 | VCVKWTCAVLDF | 24 | 0.05% | |
| 33 | VCSVWVAFWLWLC | 23 | 0.05% | |
| 34 | WFWFWICVRALCWL | 20 | 0.04% | |
| 35 | YLTSYFTSWCCFLFP | 18 | 0.04% | |
| 36 | YCVWWLSFVLFR | 17 | 0.03% | |
| 37 | FYLHGAIPSLFCLHY | 17 | 0.03% | |
| 38 | LVWFKFYIFVY | 15 | 0.03% | |
| 39 | LVLVVVHWMPLVY | 14 | 0.03% | |
| 40 | LVWVRFYIFVY | 14 | 0.03% | |
| 41 | LIWFRFYIFVY | 13 | 0.03% | |
| 42 | LVWFRFYIFIY | 13 | 0.03% | |
| 43 | SHLLWLLLGMHLV | 12 | 0.02% | |
| 44 | LVLRWYGSRFFPY | 12 | 0.02% | |
| 45 | PVWFRFYIFVY | 11 | 0.02% | |
| 46 | LVWLRFYIFVY | 11 | 0.02% | |
| 47 | VSILCRYYWWPRMS | 11 | 0.02% | |
| 48 | LLGVNQNW | 10 | 0.02% | |
| 49 | NVLMWRLQVDG | 10 | 0.02% | |
| 50 | FWLRWVFFEFLVY | 10 | 0.02% | |
| 51 | LVWFRFYILVY | 10 | 0.02% | |
| 52 | LVWFRFHIFVY | 9 | 0.02% | |
| 53 | LVWFRLYIFVY | 9 | 0.02% | |
| 54 | VMLRWFLAYFIHV | 9 | 0.02% | |
| 55 | CGLPLFLFPYFLF | 9 | 0.02% | |
| 56 | IRRGLGGFGFFRW | 8 | 0.02% | |
| 57 | GLVFFEWFKAVTTTV | 8 | 0.02% | |
| 58 | LVWFRFYIFVF | 8 | 0.02% | |

**Round 4**

| Rank | Loop 1 | read count | read frequency | U-body name |
|---|---|---|---|---|
| **1** | **LVWFRFYIFVY** | **57443** | **96.29%** | **RL1-1** |
| **2** | **FYLHGAVPSLFCLHY** | **786** | **1.32%** | **RL1-2** |
| **3** | **ALLWCNVCECFW** | **290** | **0.49%** | **RL1-3** |
| 4 | YILFIWSGVRFLS | 125 | 0.21% | |
| **5** | **LHIRWYDSLYLWL** | **115** | **0.19%** | **RL1-4** |
| 6 | RGLVPWSLR | 111 | 0.19% | **RL1-6** |
| **7** | **WCYFCGLGRFVSSWW** | **91** | **0.15%** | **RL1-5** |
| 8 | RGRSKCVFLLVCL | 81 | 0.14% | |
| 9 | SLWYFVWLTSRFST | 44 | 0.07% | |
| 10 | VCMRWSCRAFSF | 41 | 0.07% | |
| 11 | LVWFKFYIFVY | 40 | 0.07% | |
| 12 | RGLLLWSSR | 26 | 0.04% | |
| 13 | CIILWYFLIS | 19 | 0.03% | |
| 14 | LVLFSMWWVVHF | 18 | 0.03% | |
| 15 | LVWFRFYIFIY | 16 | 0.03% | |
| 16 | FVWFRFYIFVY | 16 | 0.03% | |
| 17 | LIWFRFYIFVY | 15 | 0.03% | |
| 18 | SFVLLPWWGLHQL | 15 | 0.03% | |
| 19 | LVWLRFYIFVY | 15 | 0.03% | |
| 20 | LVWFRVYIFVY | 15 | 0.03% | |
| 21 | LAWFRFYIFVY | 15 | 0.03% | |
| 22 | LVWFRFHIFVY | 13 | 0.02% | |
| 23 | LVRFRFYIFVY | 11 | 0.02% | |
| 24 | LLVQWWLTYLVY | 11 | 0.02% | |
| 25 | LVWFRFYVFVY | 10 | 0.02% | |
| 26 | LVWFRFYILVY | 10 | 0.02% | |
| 27 | IWWFYSYPHLGPSEL | 10 | 0.02% | |
| 28 | LVWFRFYIFAY | 10 | 0.02% | |
| 29 | LVWFRFYIFVH | 10 | 0.02% | |
| 30 | LYVPWWITFLRL | 9 | 0.02% | |
| 31 | LVWFRFCIFVY | 9 | 0.02% | |
| 32 | LVWFRLYIFVY | 8 | 0.01% | |
| 33 | LVWFRSYIFVY | 8 | 0.01% | |
| 34 | LVWFRFYIFVF | 8 | 0.01% | |
| 35 | LVWFGFYIFVY | 8 | 0.01% | |
| 36 | LVWFRFYI | 8 | 0.01% | |
| 37 | PVWFRFYIFVY | 7 | 0.01% | |
| 38 | LVLFRFYIFVY | 7 | 0.01% | |
| 39 | LVWFRFFIFVY | 7 | 0.01% | |
| 40 | LVLSWWMTLLWF | 7 | 0.01% | |
| 41 | LVWSRFYIFVY | 7 | 0.01% | |
| 42 | LVWFRFYISVY | 7 | 0.01% | |
| 43 | SVFRWYASSWLHL | 7 | 0.01% | |
| 44 | VCVKWTCAILDF | 6 | 0.01% | |
| 45 | LVWFRFYIFVC | 6 | 0.01% | |
| 46 | RGLVAWSLR | 5 | 0.01% | |
| 47 | LVWVRFYIFVY | 5 | 0.01% | |
| 48 | LVWFWFYIFVY | 5 | 0.01% | |
| 49 | IVWFRFYIFVY | 5 | 0.01% | |
| 50 | KLWCASCLIWW | 4 | 0.01% | |
| 51 | ICMLWNCWSVTF | 4 | 0.01% | |
| 52 | LVWFRFYTFVY | 4 | 0.01% | |
| 53 | ASWRWFSSFFLDL | 4 | 0.01% | |
| 54 | LVWFRFYIFDY | 4 | 0.01% | |
| 55 | LCLLWTCRSYSW | 3 | 0.01% | |
| 56 | LGWFRFYIFVY | 3 | 0.01% | |
| 57 | LVWFRFYIIVY | 3 | 0.01% | |
| 58 | LVSFRFYIFVY | 3 | 0.01% | |

**Round 5**

| Rank | Loop 1 | read count | read frequency | U-body name |
|---|---|---|---|---|
| **1** | **LVWFRFYIFVY** | **93450** | **96.94%** | **RL1-1** |
| **2** | **ALLWCNVCECFW** | **581** | **0.60%** | **RL1-3** |
| **3** | **RGLVPWSLR** | **404** | **0.42%** | **RL1-6** |
| 4 | RGRSKCVFLLVCL | 332 | 0.34% | |
| 5 | YILFIWSGVRFLS | 311 | 0.32% | |
| **6** | **FYLHGAVPSLFCLHY** | **286** | **0.30%** | **RL1-2** |
| **7** | **LHIRWYDSLYLWL** | **184** | **0.19%** | **RL1-4** |
| 8 | RGLLLWSSR | 105 | 0.11% | |
| 9 | LVWFKFYIFVY | 68 | 0.07% | |
| 10 | FVWFRFYIFVY | 32 | 0.03% | |
| 11 | LVWFRFYI | 31 | 0.03% | |
| **12** | **WCYFCGLGRFVSSWW** | **30** | **0.03%** | **RL1-5** |
| 13 | LVWFRFYIFIY | 28 | 0.03% | |
| 14 | LVWFRVYIFVY | 25 | 0.03% | |
| 15 | LVWFRFHIFVY | 21 | 0.02% | |
| 16 | VCMRWSCRAFSF | 21 | 0.02% | |
| 17 | LVRFRFYIFVY | 18 | 0.02% | |
| 18 | LIWFRFYIFVY | 17 | 0.02% | |
| 19 | LVLFRFYIFVY | 15 | 0.02% | |
| 20 | LVWLRFYIFVY | 15 | 0.02% | |
| 21 | LVWFRSYIFVY | 15 | 0.02% | |
| 22 | LVWFGFYIFVY | 14 | 0.01% | |
| 23 | LVLFSMWWVVHF | 14 | 0.01% | |
| 24 | IVWFRFYIFVY | 14 | 0.01% | |
| 25 | LVWFRFYILVY | 14 | 0.01% | |
| 26 | LVWFRFCIFVY | 13 | 0.01% | |
| 27 | LVWFRFFIFVY | 13 | 0.01% | |
| 28 | LVWFRFYISVY | 13 | 0.01% | |
| 29 | LVWFRFYIFVC | 12 | 0.01% | |
| 30 | LVWFRFYIFAY | 12 | 0.01% | |
| 31 | LVWSRFYIFVY | 12 | 0.01% | |
| 32 | CIILWYFLIS | 12 | 0.01% | |
| 33 | LVWFRLYIFVY | 12 | 0.01% | |
| 34 | LVWFSFYIFVY | 11 | 0.01% | |
| 35 | LVWFRFYIFV | 11 | 0.01% | |
| 36 | LVWFRFYIFVF | 10 | 0.01% | |
| 37 | LYVPWWITFLRL | 9 | 0.01% | |
| 38 | LVWFRFYIFDY | 9 | 0.01% | |
| 39 | LVWFWFYIFVY | 9 | 0.01% | |
| 40 | SFVLLPWWGLHQL | 9 | 0.01% | |
| 41 | LVWVRFYIFVY | 8 | 0.01% | |
| 42 | SVFRWYASSWLHL | 8 | 0.01% | |
| 43 | LAWFRFYIFVY | 8 | 0.01% | |
| 44 | SLWYFVWLTSRFST | 8 | 0.01% | |
| 45 | LVWFRFYVFVY | 8 | 0.01% | |
| 46 | PVWFRFYIFVY | 7 | 0.01% | |
| 47 | VCVKWTCAILDF | 6 | 0.01% | |
| 48 | LVWFRFYNFVY | 6 | 0.01% | |
| 49 | LDWFRFYIFVY | 6 | 0.01% | |
| 50 | LVWIRFYIFVY | 6 | 0.01% | |
| 51 | LVWYRFYIFVY | 5 | 0.01% | |
| 52 | LVWFRFYIIVY | 5 | 0.01% | |
| 53 | ASWRWFSSFFLDL | 5 | 0.01% | |
| 54 | RGLVAWSLR | 5 | 0.01% | |
| 55 | LVWFRFYIFVH | 4 | 0.00% | |
| 56 | IWWFYSYPHLGPSEL | 4 | 0.00% | |
| 57 | LVCFRFYIFVY | 4 | 0.00% | |
| 58 | LVLSWWMTLLWF | 4 | 0.00% | |

| | | | |
|---|---|---|---|
| 59 | LFVHLGVFRYYHL | 8 | 0.02% |
| 60 | LVRFRFYIFVY | 8 | 0.02% |
| 61 | FFFRWYAHFVLWF | 8 | 0.02% |
| 62 | FWWRWWASVLLEI | 8 | 0.02% |
| 63 | SYGPWVPFWFWLF | 8 | 0.02% |
| 64 | LFLHMFWHRMLLY | 8 | 0.02% |
| 65 | GSSLWSSWVC | 7 | 0.01% |
| 66 | LFLRWYAYLLLLY | 7 | 0.01% |
| 67 | LAWFRFYIFVY | 7 | 0.01% |
| 68 | PLGVYPVGGVGWLLQ | 7 | 0.01% |
| 69 | HIWWGWGRRLVGS | 7 | 0.01% |
| 70 | WLAFVIRDFWWVL | 6 | 0.01% |
| 71 | LRLLSRIVF | 6 | 0.01% |
| 72 | RSQVYLYEVQGRLS | 6 | 0.01% |
| 73 | FYLHGAVPSLFYLHY | 6 | 0.01% |
| 74 | FYLHGAVPSFFCLHY | 6 | 0.01% |
| 75 | WCYFCSLGRFVSSWW | 6 | 0.01% |
| 76 | ISAGLWVWPYLLF | 6 | 0.01% |
| 77 | LVWFRFYVFVY | 6 | 0.01% |
| 78 | LVWFRSYIFVY | 6 | 0.01% |
| 79 | GNLPWLLFL | 6 | 0.01% |
| 80 | LVWFRFYI | 6 | 0.01% |
| 81 | LRWGYVRGVGVVGLL | 5 | 0.01% |
| 82 | LLFTILGPLA | 5 | 0.01% |
| 83 | GAFVMQFVR | 5 | 0.01% |
| 84 | LFWFRFYIFVY | 5 | 0.01% |
| 85 | LVWFRVYIFVY | 5 | 0.01% |
| 86 | VVWWEGARVVVCFRS | 5 | 0.01% |
| 87 | FVVSAQPLCSWMF | 5 | 0.01% |
| 88 | YLDSWLFVRVMWW | 5 | 0.01% |
| 89 | LVWFRFYFFVY | 5 | 0.01% |
| 90 | GMVFLSSVFRVMLIV | 5 | 0.01% |
| 91 | LNDVFWPW | 5 | 0.01% |
| 92 | LVWFRFYIFAY | 5 | 0.01% |
| 93 | LYQRLLESSSS | 5 | 0.01% |
| 94 | LVWFGFYIFVY | 5 | 0.01% |
| 95 | RFTWFQFLLL | 5 | 0.01% |
| 96 | LWVGVIFVGWATYV | 5 | 0.01% |
| 97 | CIRWGRMGGCLQSTA | 5 | 0.01% |
| 98 | LTGSYLNWQSLRS | 5 | 0.01% |
| 99 | GLFLRREWWVGCGGG | 5 | 0.01% |
| 100 | CCIGVIWAFFGGGMS | 5 | 0.01% |
| | *total reads (top 100)* | 48716 | |

| | | | |
|---|---|---|---|
| 59 | LVWFRIYIFVY | 3 | 0.01% |
| 60 | LVCFRFYIFVY | 3 | 0.01% |
| 61 | YAIWWLGLFLLFF | 3 | 0.01% |
| 62 | VGFGFFLPFGYFLF | 3 | 0.01% |
| 63 | LVWFRFYNFVY | 2 | 0.00% |
| 64 | LVWFRFYICVY | 2 | 0.00% |
| 65 | LVWYRFYIFVY | 2 | 0.00% |
| 66 | LVWIRFYIFVY | 2 | 0.00% |
| 67 | LVWFMFYIFVY | 2 | 0.00% |
| 68 | LVWFSFYIFVY | 2 | 0.00% |
| 69 | SLWYFVWLTSRFLT | 2 | 0.00% |
| 70 | VALAMGDF | 2 | 0.00% |
| 71 | LDWFRFYIFVY | 2 | 0.00% |
| 72 | FYLHGAIPSLFCLHY | 2 | 0.00% |
| 73 | LFVHLGVFRYYHL | 2 | 0.00% |
| 74 | IRRGLGGFGFFRW | 2 | 0.00% |
| 75 | LPSWCVCG | 1 | 0.00% |
| 76 | CLSMCLCVYVL | 1 | 0.00% |
| 77 | CRSVSVFNVRLSADG | 1 | 0.00% |
| 78 | CLRTLSWQVLSY | 1 | 0.00% |
| 79 | CLMEFCRK | 1 | 0.00% |
| 80 | CSGFGMGGLASGG | 1 | 0.00% |
| 81 | CSLVLPEVWQHH | 1 | 0.00% |
| 82 | CLLIRFLLC | 1 | 0.00% |
| 83 | CVVYRPFR | 1 | 0.00% |
| 84 | CLKFIRPFNWG | 1 | 0.00% |
| 85 | LMFMLWILDVFFV | 1 | 0.00% |
| 86 | FLHQITGLFGF | 1 | 0.00% |
| 87 | DCMLTLFIPVNF | 1 | 0.00% |
| 88 | CLFREFGSGSLYTV | 1 | 0.00% |
| 89 | LLRLRLCFYSHFV | 1 | 0.00% |
| 90 | LLNRSVKHT | 1 | 0.00% |
| 91 | LLLQVLSVLRSHA | 1 | 0.00% |
| 92 | LLCQLLFGIYL | 1 | 0.00% |
| 93 | DLLMHRVISWVGVYY | 1 | 0.00% |
| 94 | LIQWFFIRLLSPG | 1 | 0.00% |
| 95 | LHREALARLRHNSR | 1 | 0.00% |
| 96 | DLRLAWLV | 1 | 0.00% |
| 97 | LHAWVLVL | 1 | 0.00% |
| 98 | LGWVAVSGAGA | 1 | 0.00% |
| 99 | DLWVMVIVRLIASS | 1 | 0.00% |
| 100 | LGVVGVSVGG | 1 | 0.00% |
| | *total reads (top 100)* | 59655 | |

| | | | |
|---|---|---|---|
| 59 | LVWFRYYIFVY | 4 | 0.00% |
| 60 | LVWFRFYIYVY | 4 | 0.00% |
| 61 | LLVQWWLTYLVY | 3 | 0.00% |
| 62 | LVWFRFYIFFY | 3 | 0.00% |
| 63 | LVWFRFYIFGY | 3 | 0.00% |
| 64 | LGWFRFYIFVY | 3 | 0.00% |
| 65 | LHIRWYDSLYLWF | 3 | 0.00% |
| 66 | VCMRWSCQAFSF | 3 | 0.00% |
| 67 | LVWFRFYFFVY | 3 | 0.00% |
| 68 | LFWFRFYIFVY | 3 | 0.00% |
| 69 | LFLHMFWHRMLLY | 3 | 0.00% |
| 70 | YAIWWLGLFLLFF | 2 | 0.00% |
| 71 | RVWFRFYIFVY | 2 | 0.00% |
| 72 | LVWFMFYIFVY | 2 | 0.00% |
| 73 | SLWYFVWLTSRFLT | 2 | 0.00% |
| 74 | LVSFRFYIFVY | 2 | 0.00% |
| 75 | HVWFRFYIFVY | 2 | 0.00% |
| 76 | LVWFRIYIFVY | 2 | 0.00% |
| 77 | VCVKWTCAVLDF | 2 | 0.00% |
| 78 | KLWCASCLIWW | 2 | 0.00% |
| 79 | LVWFRFYTFVY | 2 | 0.00% |
| 80 | ARLTILLRGV | 1 | 0.00% |
| 81 | ALLWCNVCKCFW | 1 | 0.00% |
| 82 | GTSLVLPLLWLALF | 1 | 0.00% |
| 83 | ICMLWNCWSVTF | 1 | 0.00% |
| 84 | LVCFRFYI | 1 | 0.00% |
| 85 | IVWFRFYIFVN | 1 | 0.00% |
| 86 | LCFVFVRGVCVKDR | 1 | 0.00% |
| 87 | LVGFRFYIFVY | 1 | 0.00% |
| 88 | CWKEFDTHFVD | 1 | 0.00% |
| 89 | LCLYSVSC | 1 | 0.00% |
| 90 | EVIRPIFWGLVLF | 1 | 0.00% |
| 91 | EMSRWVFASVR | 1 | 0.00% |
| 92 | YLGFLWFSSGTWYL | 1 | 0.00% |
| 93 | LVSGIAGAGVR | 1 | 0.00% |
| 94 | RYSNFLASDF | 1 | 0.00% |
| 95 | RGLVLWSLR | 1 | 0.00% |
| 96 | RGLVPWSSR | 1 | 0.00% |
| 97 | RGRSKCVF | 1 | 0.00% |
| 98 | RGRSKCVFLLICL | 1 | 0.00% |
| 99 | RSLVPWSLR | 1 | 0.00% |
| 100 | RVTYIVLETMT | 1 | 0.00% |
| | *total reads (top 100)* | 96395 | |

Supporting Information Table S1

### Page 2 - Lasso-graft library LG1 against ROR1

Top 100 sequences from Lasso-Graft library LG1 against ROR1 in different selection rounds, with NGS read frequency. Bold entries indicate the sequences selected for further analysis.
The grafted sequence is surrounded by dashes for clarity. Red text indicates mutations occurred in the grafted sequence.
The Ubiquitin core residues around the randomized loop 1 insert are ommitted for clarity. They span Ub1-6 and Ub11-76. Ub7-8 (TL) and Ub9-10 (TG) are included in the shown sequences, and are encoded by default in the library design.

**Round 3**

| Rank | Loop 1 | read count | read frequency | U-body name |
|---|---|---|---|---|
| **1** | **TLLWW-----YWYWTGNNWPYV-----NTLTG** | **2399** | **14.32%** | **LG1-1** |
| 2 | TLCWW-----YWYWTGNNWPYV-----NTLTG | 1220 | 7.28% |  |
| 3 | TLLWW-----YWYWTGNNWPYV-----GKLTG | 1094 | 6.53% |  |
| 4 | TLLWW-----YWYWTGNNWPYV-----SNLTG | 783 | 4.67% |  |
| 5 | TLCWW-----YWYWTGNNWPYV-----GTLTG | 665 | 3.97% |  |
| 6 | TLLWW-----YWYWTGNNWPYV-----DTLTG | 652 | 3.89% |  |
| 7 | TLLWW-----YWYWTGNNWPYV-----NSLTG | 589 | 3.52% |  |
| 8 | TLCWW-----YWYWTGNNWPYV-----STLTG | 486 | 2.90% |  |
| 9 | TLLWW-----YWYWTGNNWPYV-----SKLTG | 467 | 2.79% |  |
| 10 | TLLWW-----YWYWTGNNWPYV-----SSLTG | 436 | 2.60% |  |
| 11 | TLLWW-----YWYWTGNNWPYV-----GTLTG | 435 | 2.60% |  |
| 12 | TLLWW-----YWYWTGNNWPYV-----SRLTG | 409 | 2.44% |  |
| 13 | TLMWW-----YWYWTGNNWPYV-----NTLTG | 381 | 2.27% |  |
| 14 | TLLWW-----YWYWTGNNWPYV-----STLTG | 339 | 2.02% |  |
| 15 | TLLWW-----YWYWTGNNWPYV-----GSLTG | 315 | 1.88% |  |
| 16 | TLLWW-----YWYWTGNNWPYV-----NKLTG | 283 | 1.69% |  |
| 17 | TLCWW-----YWYWTGNNWPYV-----GKLTG | 268 | 1.60% |  |
| 18 | TLLWW-----YWYWTGNNWPYV-----SDLTG | 257 | 1.53% |  |
| 19 | TLCWW-----YWYWTGNNWPYV-----DTLTG | 241 | 1.44% |  |
| 20 | TLCWW-----YWYWTGNNWPYV-----NKLTG | 241 | 1.44% |  |
| 21 | TLCWW-----YWYWTGNNWPYV-----NSLTG | 223 | 1.33% |  |
| 22 | TLLWW-----YWYWTGNNWPYV-----NRLTG | 196 | 1.17% |  |
| 23 | TLCWW-----YWYWTGNNWPYV-----GRLTG | 185 | 1.10% |  |
| 24 | TLRWW-----YWYWTGNNWPYV-----GTLTG | 184 | 1.10% |  |
| 25 | TLLWW-----YWYWTGNNWPYV-----GRLTG | 178 | 1.06% |  |
| 26 | TLRWW-----YWYWTGNNWPYV-----NTLTG | 169 | 1.01% |  |
| 27 | TLLWW-----YWYWTGNNWPYV-----SQLTG | 165 | 0.98% |  |
| 28 | TLMWW-----YWYWTGNNWPYV-----STLTG | 149 | 0.89% |  |
| **29** | **TLFLFNKYWYWTGNNWLYVPPITTG** | **142** | **0.85%** | **LG1-3** |
| 30 | TLCWW-----YWYWTGNNWPYV-----SNLTG | 106 | 0.63% |  |
| 31 | TLYRCYWYWTGNNWRMCMGVPV | 106 | 0.63% | |
| 32 | TLLWW-----YWYWTGNNWPYV-----DSLTG | 106 | 0.63% |  |
| 33 | TLLWW-----YWYWTGNNWPYV-----NQLTG | 103 | 0.61% |  |
| 34 | TLMWW-----YWYWTGNNWPYV-----SKLTG | 102 | 0.61% |  |
| 35 | TLCWW-----YWYWTGNNWPYV-----SRLTG | 94 | 0.56% |  |
| 36 | TLMWW-----YWYWTGNNWPYV-----SSLTG | 91 | 0.54% |  |
| 37 | TLCWW-----YWYWTGNNWPYV-----NRLTG | 86 | 0.51% |  |
| 38 | TLCWW-----YWYWTGNNWPYV-----SSLTG | 81 | 0.48% |  |
| 39 | TLMWW-----YWYWTGNNWPYV-----NSLTG | 79 | 0.47% |  |
| 40 | TLCWW-----YWYWTGNNWPYV-----SDLTG | 79 | 0.47% |  |
| **41** | **TLYFSIWYWYWTGNNWLYVGPSEKTG** | **79** | **0.47%** | **LG1-4** |
| 42 | TLLWW-----YWYWTGNNWPYV-----ANLTG | 75 | 0.45% |  |
| 43 | TLCWW-----YWYWTGNNWPYV-----GSLTG | 75 | 0.45% |  |
| 44 | TLLWW-----YWYWTGNNWPYV-----ETLTG | 73 | 0.44% |  |
| **45** | **TLFHCYWYWTGNNWRMCRGLPV** | **72** | **0.43%** | **LG1-5** |
| 46 | TLCWW-----YWYWTGNNWPYV-----ETLTG | 70 | 0.42% |  |
| **47** | **TLDLSWYWYWTGNNWLYVTTGRGTG** | **68** | **0.41%** | **LG1-2** |
| 48 | TLLWW-----YWYWTGNNWPYV-----GQLTG | 68 | 0.41% |  |
| 49 | TLLWW-----YWYWTGNNWPYV-----YNLTG | 65 | 0.39% |  |
| 50 | TLMWW-----YWYWTGNNWPYV-----DTLTG | 64 | 0.38% |  |
| 51 | TLMWW-----YWYWTGNNWPYV-----NKLTG | 63 | 0.38% |  |
| 52 | TLLWW-----YWYWTGNNWPYV-----TTLTG | 61 | 0.36% |  |
| 53 | TLCWW-----YWYWTGNNWPYV-----TTLTG | 59 | 0.35% |  |
| 54 | TLRWW-----YWYWTGNNWPYV-----STLTG | 57 | 0.34% |  |
| 55 | TLLFAWA-----YWYWTGNNWPYV-----AKRPTG | 55 | 0.33% |  |
| 56 | TLLWW-----YWYWTGNNWPYV-----NNLTG | 47 | 0.28% |  |
| 57 | TLMWW-----YWYWTGNNWPYV-----GTLTG | 44 | 0.26% |  |
| 58 | TLVLQP-----YWYWTGNNWPYV-----LWNWKTG | 44 | 0.26% |  |
| 59 | TLLWW-----YWYWTGNNWPYV-----YDLTG | 42 | 0.25% |  |
| 60 | TLLWW-----YWYWTGNNWPYV-----GNLTG | 42 | 0.25% |  |
| 61 | TLVLQP-----YWYWTGNNWPYV-----AWHWVTG | 41 | 0.24% |  |
| 62 | TLCWF-----YWYWTGNNWPYV-----STLTG | 38 | 0.23% |  |
| 63 | TLLWW-----YWYWTGNNWPYV-----QTLTG | 37 | 0.22% |  |
| 64 | TLLWW-----YWYWTGNNWPYV-----GELTG | 37 | 0.22% |  |
| 65 | TLMWW-----YWYWTGNNWPYV-----SRLTG | 35 | 0.21% |  |
| 66 | TLCWW-----YWYWTGNNWPYV-----NQLTG | 35 | 0.21% |  |
| 67 | TLLWW-----YWYWTGNNWPYV-----HTLTG | 31 | 0.19% |  |
| 68 | TLGSI-----YWYWTGNNWPYV-----KTTYWTG | 30 | 0.18% |  |
| 69 | TLCWW-----YWYWTGNNWPYV-----SKLTG | 30 | 0.18% |  |
| 70 | TLCWW-----YWYWTGNNWPYV-----DSLTG | 29 | 0.17% |  |
| 71 | TLLWW-----YWYWTGNNWPYV-----SELTG | 29 | 0.17% |  |
| 72 | TLCWW-----YWYWTGNNWPYV-----HTLTG | 29 | 0.17% |  |

**Round 4**

| Rank | Loop 1 | read count | read frequency | U-body name |
|---|---|---|---|---|
| **1** | **TLLWW-----YWYWTGNNWPYV-----NTLTG** | **2448** | **16.23%** | **LG1-1** |
| 2 | TLCWW-----YWYWTGNNWPYV-----NTLTG | 1312 | 8.70% |  |
| 3 | TLLWW-----YWYWTGNNWPYV-----GKLTG | 920 | 6.10% |  |
| **4** | **TLFLFNKYWYWTGNNWLYVPPITTG** | **696** | **4.62%** | **LG1-3** |
| 5 | TLLWW-----YWYWTGNNWPYV-----SNLTG | 634 | 4.20% |  |
| 6 | TLCWW-----YWYWTGNNWPYV-----GTLTG | 584 | 3.87% |  |
| 7 | TLLWW-----YWYWTGNNWPYV-----NSLTG | 573 | 3.80% |  |
| 8 | TLMWW-----YWYWTGNNWPYV-----NTLTG | 456 | 3.02% |  |
| **9** | **TLFHCYWYWTGNNWRMCRGLPV** | **447** | **2.96%** | **LG1-5** |
| 10 | TLLWW-----YWYWTGNNWPYV-----DTLTG | 368 | 2.44% |  |
| 11 | TLYRCYWYWTGNNWRMCMGVPV | 365 | 2.42% | |
| 12 | TLLWW-----YWYWTGNNWPYV-----SKLTG | 364 | 2.41% |  |
| 13 | TLCWW-----YWYWTGNNWPYV-----STLTG | 352 | 2.33% |  |
| **14** | **TLDLSWYWYWTGNNWLYVTTGRGTG** | **337** | **2.23%** | **LG1-2** |
| 15 | TLLWW-----YWYWTGNNWPYV-----NKLTG | 284 | 1.88% |  |
| 16 | TLLWW-----YWYWTGNNWPYV-----SSLTG | 282 | 1.87% |  |
| 17 | TLLWW-----YWYWTGNNWPYV-----SRLTG | 279 | 1.85% |  |
| 18 | TLLWW-----YWYWTGNNWPYV-----GTLTG | 274 | 1.82% |  |
| 19 | TLLWW-----YWYWTGNNWPYV-----SDLTG | 243 | 1.61% |  |
| 20 | TLCWW-----YWYWTGNNWPYV-----NKLTG | 235 | 1.56% |  |
| 21 | TLLWW-----YWYWTGNNWPYV-----STLTG | 199 | 1.32% |  |
| **22** | **TLYFSIWYWYWTGNNWLYVGPSEKTG** | **195** | **1.29%** | **LG1-4** |
| 23 | TLCWW-----YWYWTGNNWPYV-----DTLTG | 176 | 1.17% |  |
| 24 | TLCWW-----YWYWTGNNWPYV-----GKLTG | 162 | 1.07% |  |
| 25 | TLCWW-----YWYWTGNNWPYV-----NSLTG | 160 | 1.06% |  |
| 26 | TLLWW-----YWYWTGNNWPYV-----GSLTG | 160 | 1.06% |  |
| 27 | TLLWW-----YWYWTGNNWPYV-----NRLTG | 128 | 0.85% |  |
| 28 | TLRWW-----YWYWTGNNWPYV-----NTLTG | 118 | 0.78% |  |
| 29 | TLLWW-----YWYWTGNNWPYV-----SQLTG | 118 | 0.78% |  |
| 30 | TLLWW-----YWYWTGNNWPYV-----GRLTG | 114 | 0.76% |  |
| 31 | TLRWW-----YWYWTGNNWPYV-----GTLTG | 112 | 0.74% |  |
| 32 | TLCWW-----YWYWTGNNWPYV-----GRLTG | 103 | 0.68% |  |
| 33 | TLMWW-----YWYWTGNNWPYV-----STLTG | 81 | 0.54% |  |
| 34 | TLLFAWA-----YWYWTGNNWPYV-----AKRPTG | 76 | 0.50% |  |
| 35 | TLMWW-----YWYWTGNNWPYV-----SKLTG | 71 | 0.47% |  |
| 36 | TLLWW-----YWYWTGNNWPYV-----NQLTG | 65 | 0.43% |  |
| 37 | TLCWW-----YWYWTGNNWPYV-----SDLTG | 63 | 0.42% |  |
| 38 | TLMWW-----YWYWTGNNWPYV-----DTLTG | 61 | 0.40% |  |
| 39 | TLVLQP-----YWYWTGNNWPYV-----AWHWVTG | 53 | 0.35% |  |
| 40 | TLLWW-----YWYWTGNNWPYV-----ETLTG | 52 | 0.34% |  |
| 41 | TLCWW-----YWYWTGNNWPYV-----SNLTG | 51 | 0.34% |  |
| 42 | TLMWW-----YWYWTGNNWPYV-----NSLTG | 50 | 0.33% |  |
| 43 | TLLWW-----YWYWTGNNWPYV-----GQLTG | 50 | 0.33% |  |
| 44 | TLLWW-----YWYWTGNNWPYV-----DSLTG | 50 | 0.33% |  |
| 45 | TLFPCYWYWTGNNWRMCRGLPV | 48 | 0.32% | |
| 46 | TLCWW-----YWYWTGNNWPYV-----SSLTG | 48 | 0.32% |  |
| 47 | TLMWW-----YWYWTGNNWPYV-----SSLTG | 45 | 0.30% |  |
| 48 | TLVLQP-----YWYWTGNNWPYV-----LWNWKTG | 43 | 0.29% |  |
| 49 | TLCWW-----YWYWTGNNWPYV-----GSLTG | 42 | 0.28% |  |
| 50 | TLCWW-----YWYWTGNNWPYV-----SRLTG | 38 | 0.25% |  |
| 51 | TLCWF-----YWYWTGNNWPYV-----STLTG | 38 | 0.25% |  |
| 52 | TLCEYCYWYWTGNNWPYWSVWGPV | 37 | 0.25% | |
| 53 | TLMWW-----YWYWTGNNWPYV-----NKLTG | 37 | 0.25% |  |
| 54 | TLCWW-----YWYWTGNNWPYV-----NRLTG | 35 | 0.23% |  |
| 55 | TLLWW-----YWYWTGNNWPYV-----YNLTG | 32 | 0.21% |  |
| 56 | TLLWW-----YWYWTGNNWPYV-----YDLTG | 32 | 0.21% |  |
| 57 | TLLWW-----YWYWTGNNWPYV-----GNLTG | 29 | 0.19% |  |
| 58 | TLCWW-----YWYWTGNNWPYV-----ETLTG | 29 | 0.19% |  |
| 59 | TLLWW-----YWYWTGNNWPYV-----ANLTG | 28 | 0.19% |  |
| 60 | TLRWW-----YWYWTGNNWPYV-----STLTG | 27 | 0.18% |  |
| 61 | TLMWW-----YWYWTGNNWPYV-----GTLTG | 25 | 0.17% |  |
| 62 | TLLWW-----YWYWTGNNWPYV-----TTLTG | 24 | 0.16% |  |
| 63 | TLLWW-----YWYWTGNNWPYV-----NSLTS | 23 | 0.15% |  |
| 64 | TLCWW-----YWYWTGNNWPYV-----TTLTG | 22 | 0.15% |  |
| 65 | TLLWW-----YWYWTGNNWPYV-----NNLTG | 21 | 0.14% |  |
| 66 | TLFVWV-----YWYWTGNNWPYV-----ENWPWTG | 21 | 0.14% |  |
| 67 | TLYSV-----YWYWTGNNWPYV-----LNWPWTG | 21 | 0.14% |  |
| 68 | TLGSI-----YWYWTGNNWPYV-----KTTYWTG | 20 | 0.13% |  |
| 69 | TLCWW-----YWYWTGNNWPYV-----SKLTG | 20 | 0.13% |  |
| 70 | TLCWW-----YWYWTGNNWPYV-----SQLTG | 19 | 0.13% |  |
| 71 | TLLWW-----YWYWTGNNWPYV-----GELTG | 18 | 0.12% |  |
| 72 | TLCWW-----YWYWTGNNWPYV-----DSLTG | 18 | 0.12% |  |

**Round 5**

| Rank | Loop 1 | read count | read frequency | U-body name |
|---|---|---|---|---|
| **1** | **TLFLFNKYWYWTGNNWLYVPPITTG** | **8750** | **21.56%** | **LG1-3** |
| **2** | **TLLWW-----YWYWTGNNWPYV-----NTLTG** | **5601** | **13.80%** | **LG1-1** |
| **3** | **TLDLSWYWYWTGNNWLYVTTGRGTG** | **4018** | **9.90%** | **LG1-2** |
| **4** | **TLFHCYWYWTGNNWRMCRGLPV** | **3675** | **9.06%** | **LG1-5** |
| 5 | TLCWW-----YWYWTGNNWPYV-----NTLTG | 2282 | 5.62% |  |
| 6 | TLLWW-----YWYWTGNNWPYV-----GKLTG | 1872 | 4.61% |  |
| **7** | **TLYFSIWYWYWTGNNWLYVGPSEKTG** | **1331** | **3.28%** | **LG1-4** |
| 8 | TLLWW-----YWYWTGNNWPYV-----SNLTG | 1115 | 2.75% |  |
| 9 | TLYRCYWYWTGNNWRMCMGVPV | 1056 | 2.60% | |
| 10 | TLLWW-----YWYWTGNNWPYV-----NSLTG | 1009 | 2.49% |  |
| 11 | TLCWW-----YWYWTGNNWPYV-----GTLTG | 742 | 1.83% |  |
| 12 | TLLWW-----YWYWTGNNWPYV-----DTLTG | 647 | 1.59% |  |
| 13 | TLLWW-----YWYWTGNNWPYV-----SKLTG | 605 | 1.49% |  |
| 14 | TLRWW-----YWYWTGNNWPYV-----NTLTG | 571 | 1.41% |  |
| 15 | TLLWW-----YWYWTGNNWPYV-----SSLTG | 436 | 1.07% |  |
| 16 | TLLWW-----YWYWTGNNWPYV-----SDLTG | 418 | 1.03% |  |
| 17 | TLLWW-----YWYWTGNNWPYV-----SRLTG | 416 | 1.03% |  |
| 18 | TLCWW-----YWYWTGNNWPYV-----NKLTG | 387 | 0.95% |  |
| 19 | TLFPCYWYWTGNNWRMCRGLPV | 371 | 0.91% | |
| 20 | TLMWW-----YWYWTGNNWPYV-----NTLTG | 357 | 0.88% |  |
| 21 | TLCWW-----YWYWTGNNWPYV-----STLTG | 323 | 0.80% |  |
| 22 | TLLWW-----YWYWTGNNWPYV-----GTLTG | 321 | 0.79% |  |
| 23 | TLLWW-----YWYWTGNNWPYV-----NKLTG | 314 | 0.77% |  |
| 24 | TLLWW-----YWYWTGNNWPYV-----STLTG | 308 | 0.76% |  |
| 25 | TLCWW-----YWYWTGNNWPYV-----NSLTG | 263 | 0.65% |  |
| 26 | TLRWW-----YWYWTGNNWPYV-----GTLTG | 257 | 0.63% |  |
| 27 | TLLWW-----YWYWTGNNWPYV-----NRLTG | 256 | 0.63% |  |
| 28 | TLCWW-----YWYWTGNNWPYV-----GKLTG | 226 | 0.56% |  |
| 29 | TLLWW-----YWYWTGNNWPYV-----GSLTG | 219 | 0.54% |  |
| 30 | TLCWW-----YWYWTGNNWPYV-----DTLTG | 209 | 0.52% |  |
| 31 | TLLWW-----YWYWTGNNWPYV-----GRLTG | 170 | 0.42% |  |
| 32 | TLLWW-----YWYWTGNNWPYV-----SQLTG | 147 | 0.36% |  |
| 33 | TLLWW-----YWYWTGNNWPYV-----NQLTG | 127 | 0.31% |  |
| 34 | TLCWW-----YWYWTGNNWPYV-----GRLTG | 106 | 0.26% |  |
| 35 | TLVLQP-----YWYWTGNNWPYV-----AWHWVTG | 100 | 0.25% |  |
| 36 | TLLFAWA-----YWYWTGNNWPYV-----AKRPTG | 94 | 0.23% |  |
| 37 | TLRWW-----YWYWTGNNWPYV-----STLTG | 70 | 0.17% |  |
| 38 | TLCWW-----YWYWTGNNWPYV-----SNLTG | 66 | 0.16% |  |
| 39 | TLLWW-----YWYWTGNNWPYV-----ANLTG | 62 | 0.15% |  |
| 40 | TLLWW-----YWYWTGNNWPYV-----GQLTG | 57 | 0.14% |  |
| 41 | TLLWW-----YWYWTGNNWPYV-----ETLTG | 53 | 0.13% |  |
| 42 | TLLWW-----YWYWTGNNWPYV-----DSLTG | 50 | 0.12% |  |
| 43 | TLCWW-----YWYWTGNNWPYV-----NRLTG | 44 | 0.11% |  |
| 44 | TLYSV-----YWYWTGNNWPYV-----MNWPWTG | 43 | 0.11% |  |
| 45 | TLRWW-----YWYWTGNNWPYV-----SSLTG | 43 | 0.11% |  |
| 46 | TLCWW-----YWYWTGNNWPYV-----GSLTG | 43 | 0.11% |  |
| 47 | TLCWW-----YWYWTGNNWPYV-----SSLTG | 41 | 0.10% |  |
| 48 | TLMWW-----YWYWTGNNWPYV-----DTLTG | 39 | 0.10% |  |
| 49 | TLRWW-----YWYWTGNNWPYV-----DTLTG | 39 | 0.10% |  |
| 50 | TLCWW-----YWYWTGNNWPYV-----SRLTG | 37 | 0.09% |  |
| 51 | TLMWW-----YWYWTGNNWPYV-----STLTG | 36 | 0.09% |  |
| 52 | TLRWW-----YWYWTGNNWPYV-----NSLTG | 29 | 0.07% |  |
| 53 | TLCWF-----YWYWTGNNWPYV-----STLTG | 29 | 0.07% |  |
| 54 | TLFDGPYWYWTGNNWLYVTELTG | 27 | 0.07% | |
| 55 | TLLWW-----YWYWTGNNWPYV-----YDLTG | 25 | 0.06% |  |
| 56 | TLMWW-----YWYWTGNNWPYV-----SKLTG | 25 | 0.06% |  |
| 57 | TLLWW-----YWYWTGNNWPYV-----HTLTG | 25 | 0.06% |  |
| 58 | TLLWW-----YWYWTGNNWPYV-----GELTG | 25 | 0.06% |  |
| 59 | TLLWW-----YWYWTGNNWPYV-----YNLTG | 24 | 0.06% |  |
| 60 | RYFCLISIGIGPATTGC | 24 | 0.06% | |
| 61 | TLCWW-----YWYWTGNNWPYV-----SDLTG | 24 | 0.06% |  |
| 62 | TLLWW-----YWYWTGNNWPYV-----TTLTG | 23 | 0.06% |  |
| 63 | TLMWW-----YWYWTGNNWPYV-----GTLTG | 22 | 0.05% |  |
| 64 | TLLWW-----YWYWTGNNWPYV-----NNLTG | 21 | 0.05% |  |
| 65 | TLCWW-----YWYWTGNNWPYV-----SKLTG | 21 | 0.05% |  |
| 66 | TLLWW-----YWYWTGNNWPYV-----GNLTG | 20 | 0.05% |  |
| 67 | TLMWW-----YWYWTGNNWPYV-----NKLTG | 19 | 0.05% |  |
| 68 | TLLWW-----YWYWTGNNWPYV-----NTLTD | 19 | 0.05% |  |
| 69 | TLWSQI-----YWYWTGNNWPYV-----TWPMTG | 19 | 0.05% |  |
| 70 | TLLWW-----YWYWTGNNWPYV-----QTLTG | 17 | 0.04% |  |
| 71 | TLGSI-----YWYWTGNNWPYV-----KTTYWTG | 17 | 0.04% |  |
| 72 | TLCWW-----YWYWTGNNWPYV-----TTLTG | 16 | 0.04% |  |

| | | | |
|---|---|---|---|
| 73 | TLRWW-----YWYWTGNNWPYV-----NSLTG | 28 | 0.17% |
| 74 | TLRWW-----YWYWTGNNWPYV-----SSLTG | 27 | 0.16% |
| 75 | TLMWW-----YWYWTGNNWPYV-----GKLTG | 26 | 0.16% |
| 76 | TLRWW-----YWYWTGNNWPYV-----DTLTG | 24 | 0.14% |
| 77 | TLCWW-----YWYWTGNNWPYV-----YSLDP | 24 | 0.14% |
| 78 | TLLWW-----YWYWTGNNWPYV-----ASLTG | 23 | 0.14% |
| 79 | TLKWW-----YWYWTGNNWPYV-----GTLTG | 22 | 0.13% |
| 80 | TLCWW-----YWYWTGNNWPYV-----SQLTG | 22 | 0.13% |
| 81 | TLVALYYWYWTGNNWLYVTTGADTG | 21 | 0.13% |
| 82 | TLCWW-----YWYWTGNNWPYV-----ARLTG | 21 | 0.13% |
| 83 | TLLWW-----YWYWTGNNWPYV-----STFTG | 21 | 0.13% |
| 84 | TLQWW-----YWYWTGNNWPYV-----NTLTG | 20 | 0.12% |
| 85 | TLCEYCYWYWTGNNWPYWSVWGPV | 19 | 0.11% |
| 86 | TLRSLFEYWYWTGNNWQYVNTTG | 19 | 0.11% |
| 87 | TLGST-----YWYWTGNNWPYV-----RVTYWTG | 18 | 0.11% |
| 88 | TLMWW-----YWYWTGNNWPYV-----SQLTG | 18 | 0.11% |
| 89 | TLMWW-----YWYWTGNNWPYV-----GQLTG | 18 | 0.11% |
| 90 | TLRWW-----YWYWTGNNWPYV-----GRLTG | 18 | 0.11% |
| 91 | TLRWW-----YWYWTGNNWPYV-----TTLTG | 17 | 0.10% |
| 92 | TLCWW-----YWYWTGNNWPYV-----SALTG | 17 | 0.10% |
| 93 | TLCWW-----YWYWTGNNWPYV-----ATLTG | 17 | 0.10% |
| 94 | TLLWW-----YWYWTGNNWPYV-----GDLTG | 16 | 0.10% |
| 95 | TLLWW-----YWYWTGNNWPYV-----NSLTS | 15 | 0.09% |
| 96 | TLCWW-----YWYWTGNNWPYV-----GQLTG | 15 | 0.09% |
| 97 | TLLWW-----YWYWTGNNWPYV-----ESLTG | 15 | 0.09% |
| 98 | TLFVWV-----YWYWTGNNWPYV-----ENWPWTG | 15 | 0.09% |
| 99 | TLLWW-----YWYWTGNNWPYV-----ARLTG | 14 | 0.08% |
| 100 | TLYSV-----YWYWTGNNWPYV-----MNWPWTG | 14 | 0.08% |
| | *total reads (top 100)* | 16752 | |

| | | | |
|---|---|---|---|
| 73 | TLLWW-----YWYWTGNNWPYV-----QTLTG | 18 | 0.12% |
| 74 | TLMWW-----YWYWTGNNWPYV-----GKLTG | 17 | 0.11% |
| 75 | TLLWW-----YWYWTGNNWPYV-----HTLTG | 16 | 0.11% |
| 76 | TLYSV-----YWYWTGNNWPYV-----MNWPWTG | 16 | 0.11% |
| 77 | TLVALYYWYWTGNNWLYVTTGADTG | 15 | 0.10% |
| 78 | TLLWW-----YWYWTGNNWPYV-----SELTG | 15 | 0.10% |
| 79 | TLMWW-----YWYWTGNNWPYV-----SRLTG | 14 | 0.09% |
| 80 | TLCWW-----YWYWTGNNWPYV-----YSLDP | 13 | 0.09% |
| 81 | TLRWW-----YWYWTGNNWPYV-----DTLTG | 13 | 0.09% |
| 82 | TLKWW-----YWYWTGNNWPYV-----NTLTG | 12 | 0.08% |
| 83 | TLRWW-----YWYWTGNNWPYV-----SSLTG | 12 | 0.08% |
| 84 | TLLWW-----YWYWTGNNWPYV-----NTLTD | 12 | 0.08% |
| 85 | TLCWW-----YWYWTGNNWPYV-----SELTG | 12 | 0.08% |
| 86 | TLKWW-----YWYWTGNNWPYV-----GTLTG | 11 | 0.07% |
| 87 | TLCWW-----YWYWTGNNWPYV-----NQLTG | 11 | 0.07% |
| 88 | TLYANPYWYWTGNNWLYVTELPSTG | 11 | 0.07% |
| 89 | TLCWW-----YWYWTGNNWPYV-----ATLTG | 10 | 0.07% |
| 90 | TLLWF-----YWYWTGNNWPYV-----STLTG | 10 | 0.07% |
| 91 | TLCWW-----YWYWTGNNWPYV-----HTLTG | 9 | 0.06% |
| 92 | TLWSQI-----YWYWTGNNWPYV-----TWPMTG | 9 | 0.06% |
| 93 | TLLIY-----YWYWTGNNWPYV-----KYGTG | 9 | 0.06% |
| 94 | TLLWWYWYWTGKNWPYVSHLTG | 8 | 0.05% |
| 95 | TLRWW-----YWYWTGNNWPYV-----NSLTG | 8 | 0.05% |
| 96 | TLRWW-----YWYWTGNNWPYV-----GSLTG | 8 | 0.05% |
| 97 | TLLWW-----YWYWTGNNWPYV-----ARLTG | 8 | 0.05% |
| 98 | TLRWW-----YWYWTGNNWPYV-----GRLTG | 8 | 0.05% |
| 99 | TLLVFH-----YWYWTGNNWPYV-----SEWYGTG | 8 | 0.05% |
| 100 | TLLWW-----YWYWTGNNWPYV-----YSLTG | 7 | 0.05% |
| | *total reads (top 100)* | 15081 | |

| | | | |
|---|---|---|---|
| 73 | TLMWW-----YWYWTGNNWPYV-----SSLTG | 16 | 0.04% |
| 74 | TLDLSWYWYWTGNNWLYVTTGQGTG | 16 | 0.04% |
| 75 | TLLWW-----YWYWTGNNWPYV-----SELTG | 15 | 0.04% |
| 76 | TLMWW-----YWYWTGNNWPYV-----NSLTG | 13 | 0.03% |
| 77 | TLCWW-----YWYWTGNNWPYV-----NQLTG | 13 | 0.03% |
| 78 | TLRWW-----YWYWTGNNWPYV-----GSLTG | 13 | 0.03% |
| 79 | TLCWW-----YWYWTGNNWPYV-----ETLTG | 12 | 0.03% |
| 80 | TLLWW-----YWYWTGNNWPYV-----NSLTS | 11 | 0.03% |
| 81 | TLCEYCYWYWTGNNWPYWSVWGPV | 11 | 0.03% |
| 82 | TLLWWYWYWTGKNWPYVSHLTG | 10 | 0.02% |
| 83 | TLRWW-----YWYWTGNNWPYV-----ETLTG | 10 | 0.02% |
| 84 | TLCWW-----YWYWTGNNWPYV-----DSLTG | 10 | 0.02% |
| 85 | TLCWW-----YWYWTGNNWPYV-----HTLTG | 10 | 0.02% |
| 86 | TLYSV-----YWYWTGNNWPYV-----LNWPWTG | 9 | 0.02% |
| 87 | TLVLQP-----YWYWTGNNWPYV-----LWNWKTG | 9 | 0.02% |
| 88 | TLCWW-----YWYWTGNNWPYV-----YSLDP | 9 | 0.02% |
| 89 | TLRSLFEYWYWTGNNWQYVNTTG | 9 | 0.02% |
| 90 | TLMWW-----YWYWTGNNWPYV-----GKLTG | 9 | 0.02% |
| 91 | TLLWW-----YWYWTGNNWPYV-----GDLTG | 9 | 0.02% |
| 92 | TLLWWYWYWTGNNWLYVTTGRGTG | 9 | 0.02% |
| 93 | TLCWW-----YWYWTGNNWPYV-----SQLTG | 9 | 0.02% |
| 94 | TLDISWYWYWTGNNWLYVTTGRGTG | 8 | 0.02% |
| 95 | TLFVWV-----YWYWTGNNWPYV-----ENWPWTG | 8 | 0.02% |
| 96 | TLFLFNKYWYWTGNNWLYVPPITTS | 8 | 0.02% |
| 97 | TLLWWYWYWTSNNWPYVNTLTG | 7 | 0.02% |
| 98 | TLFLFNK-----YWYWTGNNWPYV-----NTLTG | 7 | 0.02% |
| 99 | TLVSGGWYRYWTGNNWPYVWVTTG | 7 | 0.02% |
| 100 | TLLWW-----YWYWTGNNWPYV-----AKLTG | 7 | 0.02% |
| | *total reads (top 100)* | 40577 | |

### Supporting Information Table S1

#### Page 3 - Lasso-graft library LG2 against ROR1

Top 100 sequences from Lasso-Graft library LG2 against ROR1 in different selection rounds, with NGS read frequency. Bold entries indicate the sequences selected for further analysis.
The grafted sequence is surrounded by dashes for clarity. Red text indicates mutations occurred in the grafted sequence.
The Ubiquitin core residues around the randomized loop 1 insert are ommitted for clarity. They span Ub1-6 and Ub11-76. Ub7-8 (TL) and Ub9-10 (TG) are included in the shown sequences, and are encoded by default in the library design.

**Round 3**

| Rank | Loop 1 | read count | read frequency | U-body name |
|---|---|---|---|---|
| **1** | **TLCI-----YVMYWYWTGNNW-----VDCTG** | **2001** | **9.92%** | **LG2-1** |
| 2 | TLVG-----YVMYWYWTGNNW-----IPLWTG | 1222 | 6.06% |  |
| 3 | TLCI-----YVMYWYWTGNNW-----IDCTG | 1034 | 5.13% |  |
| 4 | TLCL-----YVMYWYWTGNNW-----VDCTG | 856 | 4.25% |  |
| 5 | TLVYE-----YVMYWYWTGNNW-----IPMRTG | 702 | 3.48% |  |
| 6 | TLCM-----YVMYWYWTGNNW-----VDCHTG | 566 | 2.81% |  |
| 7 | TLCM-----YVMYWYWTGNNW-----VGCWPTG | 563 | 2.79% |  |
| 8 | TLVG-----YVMYWYWTGNNW-----IPMLTG | 534 | 2.65% |  |
| 9 | TLCM-----YVMYWYWTGNNW-----VDCTG | 435 | 2.16% |  |
| 10 | TLCL-----YVMYWYWTGNNW-----FDCHTG | 358 | 1.78% |  |
| 11 | TLCI-----YVMYWYWTGNNW-----IDCSGTG | 334 | 1.66% |  |
| 12 | TLCV-----YVMYWYWTGNNW-----IACQGTG | 325 | 1.61% |  |
| 13 | TLCI-----YVMYWYWTGNNW-----IDCHTG | 323 | 1.60% |  |
| 14 | TLCL-----YVMYWYWTGNNW-----VDCWTG | 309 | 1.53% |  |
| 15 | TLCL-----YVMYWYWTGNNW-----VDCYTG | 290 | 1.44% |  |
| 16 | TLCL-----YVMYWYWTGNNW-----VDCHTG | 290 | 1.44% |  |
| **17** | **TLYH-----YVMYWYWTGNNW-----LLGNRTG** | **272** | **1.35%** | **LG2-4** |
| 18 | TLVG-----YVMYWYWTGNNW-----IPMWTG | 260 | 1.29% |  |
| 19 | TLCM-----YVMYWYWTGNNW-----VDCHGTG | 255 | 1.26% |  |
| 20 | TLCL-----YVMYWYWTGNNW-----IDCTG | 234 | 1.16% |  |
| 21 | TLFDN-----YVMYWYWTGNNW-----IPITTG | 231 | 1.15% |  |
| 22 | TLCM-----YVMYWYWTGNNW-----TDCT | 221 | 1.10% |  |
| **23** | **TLCL-----YVMYWYWTGNNW-----SDACTG** | **212** | **1.05%** | **LG2-2** |
| 24 | TLCM-----YVMYWYWTGNNW-----VPCT | 205 | 1.02% |  |
| 25 | TLCL-----YVMYWYWTGNNW-----IDCWTG | 204 | 1.01% |  |
| 26 | TLCM-----YVMYWYWTGNNW-----IDCTG | 201 | 1.00% |  |
| 27 | TLVMP-----YVMYWYWTGNNW-----IPMRTG | 200 | 0.99% |  |
| 28 | TLCM-----YVMYWYWTGNNW-----VPCHTG | 199 | 0.99% |  |
| 29 | TLCL-----YVMYWYWTGNNW-----THCQGTG | 192 | 0.95% |  |
| 30 | TLDG-----YVMYWYWTGNNW-----IPLWTG | 189 | 0.94% |  |
| 31 | TLCL-----YVMYWYWTGNNW-----IDCFTG | 178 | 0.88% |  |
| 32 | TLCL-----YVMYWYWTGNNW-----VDCATG | 175 | 0.87% |  |
| 33 | TLFDN-----YVMYWYWTGNNW-----FPVTTG | 172 | 0.85% |  |
| 34 | TLCI-----YVMYWYWTGNNW-----MDCTG | 172 | 0.85% |  |
| 35 | TLCL-----YVMYWYWTGNNW-----VPCHGTG | 167 | 0.83% |  |
| 36 | TLCL-----YVMYWYWTGNNW-----TGCYPTG | 163 | 0.81% |  |
| 37 | TLFH-----YVMYWYWTGNNW-----VLGNRTG | 158 | 0.78% |  |
| 38 | TLCV-----YVMYWYWTGNNW-----VQCQGTG | 156 | 0.77% |  |
| 39 | TLHCSPC-----YVMYWYWTGNNW-----AFTG | 148 | 0.73% |  |
| 40 | TLCV-----YVMYWYWTGNNW-----THCQGTG | 142 | 0.70% |  |
| 41 | TLCI-----YVMYWYWTGNNW-----VDCHTG | 137 | 0.68% |  |
| 42 | TLCV-----YVMYWYWTGNNW-----IPCQGTG | 136 | 0.67% |  |
| **43** | **TLCLFTLYWYWTGNNWVECHHTG** | **130** | **0.64%** | **LG2-3** |
| 44 | TLCL-----YVMYWYWTGNNW-----VPCNGTG | 129 | 0.64% |  |
| 45 | TLL-----YVMYWYWTGNNW-----IPIMRTG | 128 | 0.63% |  |
| 46 | TLCM-----YVMYWYWTGNNW-----VPCHETG | 127 | 0.63% |  |
| 47 | TLVG-----YVMYWYWTGNNW-----FPMWTG | 122 | 0.61% |  |
| 48 | TLWSLP-----YVMYWYWTGNNW-----LITRNTG | 120 | 0.60% |  |
| 49 | TLCM-----YVMYWYWTGNNW-----FDCHTG | 120 | 0.60% |  |
| 50 | TLCL-----YVMYWYWTGNNW-----VDCADTG | 120 | 0.60% |  |
| 51 | TLCV-----YVMYWYWTGNNW-----VECQGTG | 119 | 0.59% |  |
| 52 | TLCL-----YVMYWYWTGNNW-----VPCQGTG | 118 | 0.59% |  |
| 53 | TLQCYFN-----YVMYWYWTGNNW-----VS | 116 | 0.58% |  |
| 54 | TLYTRSL-----YVMYWYWTGNNW-----MGSTG | 116 | 0.58% |  |
| 55 | TLYH-----YVMYWYWTGNNW-----IKGNRTG | 115 | 0.57% |  |
| 56 | TLW-----YVMYWYWTGNNW-----INGIHTG | 109 | 0.54% |  |
| 57 | TLGPP-----YVMYWYWTGNNW-----FGPTG | 107 | 0.53% |  |
| 58 | TLCL-----YVMYWYWTGNNW-----VDCYGTG | 106 | 0.53% |  |
| 59 | TLCM-----YVMYWYWTGNNW-----TDVCTG | 100 | 0.50% |  |
| 60 | TLVNW-----YVMYWYWTGNNW-----ALTG | 100 | 0.50% |  |
| 61 | TLW-----YVMYWYWTGNNW-----ITGLHTG | 99 | 0.49% |  |
| 62 | TLCV-----YVMYWYWTGNNW-----VDCTG | 96 | 0.48% |  |
| 63 | TLL-----YVMYWYWTGNNW-----LPIFRTG | 95 | 0.47% |  |
| 64 | TLCL-----YVMYWYWTGNNW-----IDCYTG | 94 | 0.47% |  |
| 65 | TLCI-----YVMYWYWTGNNW-----SDICTG | 93 | 0.46% |  |
| 66 | TLCL-----YVMYWYWTGNNW-----VDCSGTG | 92 | 0.46% |  |
| 67 | TLYDAGW-----YVMYWYWTGNNW-----IPMKNTG | 89 | 0.44% |  |
| 68 | TLCY-----YVMYWYWTGNNW-----VDCYTG | 88 | 0.44% |  |
| **69** | **TLW-----YVMYWYWTGNNW-----ITGIHTG** | **87** | **0.43%** | **LG2-5** |
| 70 | TLCM-----YVMYWYWTGNNW-----FDCTG | 87 | 0.43% |  |

**Round 4**

| Rank | Loop 1 | read count | read frequency | U-body name |
|---|---|---|---|---|
| **1** | **TLCI-----YVMYWYWTGNNW-----VDCTG** | **1904** | **11.66%** | **LG2-1** |
| 2 | TLCI-----YVMYWYWTGNNW-----IDCTG | 877 | 5.37% |  |
| 3 | TLCL-----YVMYWYWTGNNW-----VDCTG | 794 | 4.86% |  |
| 4 | TLVG-----YVMYWYWTGNNW-----IPLWTG | 613 | 3.76% |  |
| **5** | **TLCL-----YVMYWYWTGNNW-----SDACTG** | **565** | **3.46%** | **LG2-2** |
| 6 | TLVYE-----YVMYWYWTGNNW-----IPMRTG | 551 | 3.38% |  |
| 7 | TLCM-----YVMYWYWTGNNW-----VGCWPTG | 527 | 3.23% |  |
| 8 | TLCV-----YVMYWYWTGNNW-----IACQGTG | 506 | 3.10% |  |
| 9 | TLCM-----YVMYWYWTGNNW-----VDCHTG | 476 | 2.92% |  |
| 10 | TLCI-----YVMYWYWTGNNW-----IDCHTG | 392 | 2.40% |  |
| 11 | TLCL-----YVMYWYWTGNNW-----VDCYTG | 382 | 2.34% |  |
| 12 | TLCM-----YVMYWYWTGNNW-----VPCT | 366 | 2.24% |  |
| 13 | TLCL-----YVMYWYWTGNNW-----VDCWTG | 361 | 2.21% |  |
| 14 | TLCL-----YVMYWYWTGNNW-----FDCHTG | 324 | 1.98% |  |
| 15 | TLCI-----YVMYWYWTGNNW-----MDCTG | 300 | 1.84% |  |
| **16** | **TLCLFTLYWYWTGNNWVECHHTG** | **275** | **1.68%** | **LG2-3** |
| 17 | TLCI-----YVMYWYWTGNNW-----IDCSGTG | 269 | 1.65% |  |
| 18 | TLCL-----YVMYWYWTGNNW-----THCQGTG | 262 | 1.61% |  |
| 19 | TLCL-----YVMYWYWTGNNW-----IDCWTG | 251 | 1.54% |  |
| 20 | TLCM-----YVMYWYWTGNNW-----VDCHGTG | 244 | 1.49% |  |
| 21 | TLCM-----YVMYWYWTGNNW-----VDCTG | 220 | 1.35% |  |
| 22 | TLCM-----YVMYWYWTGNNW-----TDCT | 214 | 1.31% |  |
| 23 | TLCL-----YVMYWYWTGNNW-----IDCFTG | 206 | 1.26% |  |
| **24** | **TLYH-----YVMYWYWTGNNW-----LLGNRTG** | **190** | **1.16%** | **LG2-4** |
| 25 | TLFDN-----YVMYWYWTGNNW-----FPVTTG | 188 | 1.15% |  |
| 26 | TLCL-----YVMYWYWTGNNW-----VPCQGTG | 181 | 1.11% |  |
| 27 | TLCV-----YVMYWYWTGNNW-----IPCQGTG | 178 | 1.09% |  |
| 28 | TLVG-----YVMYWYWTGNNW-----IPMLTG | 178 | 1.09% |  |
| 29 | TLCL-----YVMYWYWTGNNW-----VDCHTG | 165 | 1.01% |  |
| 30 | TLQCYFN-----YVMYWYWTGNNW-----VS | 151 | 0.93% |  |
| 31 | TLCM-----YVMYWYWTGNNW-----VPCHTG | 138 | 0.85% |  |
| 32 | TLCL-----YVMYWYWTGNNW-----IDCTG | 136 | 0.83% |  |
| 33 | TLCI-----YVMYWYWTGNNW-----VNCTG | 131 | 0.80% |  |
| 34 | TLFDN-----YVMYWYWTGNNW-----IPITTG | 130 | 0.80% |  |
| 35 | TLCM-----YVMYWYWTGNNW-----VPCHETG | 127 | 0.78% |  |
| 36 | TLCL-----YVMYWYWTGNNW-----TGCYPTG | 124 | 0.76% |  |
| 37 | TLVG-----YVMYWYWTGNNW-----IPMWTG | 124 | 0.76% |  |
| 38 | TLCL-----YVMYWYWTGNNW-----VDCATG | 122 | 0.75% |  |
| 39 | TLCL-----YVMYWYWTGNNW-----IDCYTG | 115 | 0.70% |  |
| 40 | TLVMP-----YVMYWYWTGNNW-----IPMRTG | 115 | 0.70% |  |
| 41 | TLCV-----YVMYWYWTGNNW-----VQCQGTG | 113 | 0.69% |  |
| 42 | TLCI-----YVMYWYWTGNNW-----VDCHTG | 109 | 0.67% |  |
| 43 | TLCL-----YVMYWYWTGNNW-----VPCHGTG | 106 | 0.65% |  |
| 44 | TLCL-----YVMYWYWTGNNW-----VPCNGTG | 103 | 0.63% |  |
| 45 | TLCM-----YVMYWYWTGNNW-----FDCHTG | 101 | 0.62% |  |
| 46 | TLHCSPC-----YVMYWYWTGNNW-----AFTG | 94 | 0.58% |  |
| 47 | TLCV-----YVMYWYWTGNNW-----THCQGTG | 90 | 0.55% |  |
| 48 | TLCL-----YVMYWYWTGNNW-----VDCYGTG | 89 | 0.55% |  |
| 49 | TLW-----YVMYWYWTGNNW-----ITGLHTG | 72 | 0.44% |  |
| 50 | TLW-----YVMYWYWTGNNW-----ITGMHTG | 71 | 0.43% |  |
| **51** | **TLW-----YVMYWYWTGNNW-----ITGIHTG** | **71** | **0.43%** | **LG2-5** |
| 52 | TLCL-----YVMYWYWTGNNW-----VDCSGTG | 68 | 0.42% |  |
| 53 | TLW-----YVMYWYWTGNNW-----INGIHTG | 61 | 0.37% |  |
| 54 | TLCM-----YVMYWYWTGNNW-----IDCTG | 60 | 0.37% |  |
| 55 | TLFH-----YVMYWYWTGNNW-----VLGNRTG | 58 | 0.36% |  |
| 56 | TLCV-----YVMYWYWTGNNW-----VECQGTG | 57 | 0.35% |  |
| 57 | TLDG-----YVMYWYWTGNNW-----IPLWTG | 55 | 0.34% |  |
| 58 | TLCI-----YVMYWYWTGNNW-----VDCSGTG | 55 | 0.34% |  |
| 59 | TLCL-----YVMYWYWTGNNW-----VDCWNTG | 53 | 0.32% |  |
| 60 | TLCY-----YVMYWYWTGNNW-----VDCYTG | 48 | 0.29% |  |
| 61 | TLCM-----YVMYWYWTGNNW-----FDCTG | 48 | 0.29% |  |
| 62 | TLVSE-----YVMYWYWTGNNW-----IPMRTG | 47 | 0.29% |  |
| 63 | TLGPP-----YVMYWYWTGNNW-----FGPTG | 47 | 0.29% |  |
| 64 | TLCL-----YVMYWYWTGNNW-----VDCADTG | 47 | 0.29% |  |
| 65 | TLCM-----YVMYWYWTGNNW-----TDVCTG | 46 | 0.28% |  |
| 66 | TLCL-----YVMYWYWTGNNW-----VDCFTG | 45 | 0.28% |  |
| 67 | TLCL-----YVMYWYWTGNNW-----FDCTG | 45 | 0.28% |  |
| 68 | TLCV-----YVMYWYWTGNNW-----VDCNTG | 44 | 0.27% |  |
| 69 | TLCM-----YVMYWYWTGNNW-----IDCFTG | 43 | 0.26% |  |
| 70 | TLCV-----YVMYWYWTGNNW-----VDCTG | 42 | 0.26% |  |

**Round 5**

| Rank | Loop 1 | read count | read frequency | U-body name |
|---|---|---|---|---|
| **1** | **TLCI-----YVMYWYWTGNNW-----VDCTG** | **2953** | **12.11%** | **LG2-1** |
| **2** | **TLCL-----YVMYWYWTGNNW-----SDACTG** | **2288** | **9.39%** | **LG2-2** |
| **3** | **TLCLFTLYWYWTGNNWVECHHTG** | **2158** | **8.85%** | **LG2-3** |
| 4 | TLCI-----YVMYWYWTGNNW-----IDCTG | 1134 | 4.65% |  |
| 5 | TLCM-----YVMYWYWTGNNW-----VPCT | 1121 | 4.60% |  |
| 6 | TLCL-----YVMYWYWTGNNW-----VDCTG | 964 | 3.95% |  |
| 7 | TLCM-----YVMYWYWTGNNW-----VDCHTG | 764 | 3.13% |  |
| 8 | TLCM-----YVMYWYWTGNNW-----VGCWPTG | 697 | 2.86% |  |
| 9 | TLCL-----YVMYWYWTGNNW-----VDCYTG | 694 | 2.85% |  |
| 10 | TLCI-----YVMYWYWTGNNW-----MDCTG | 616 | 2.53% |  |
| 11 | TLCI-----YVMYWYWTGNNW-----IDCHTG | 609 | 2.50% |  |
| 12 | TLCL-----YVMYWYWTGNNW-----VDCWTG | 607 | 2.49% |  |
| 13 | TLCM-----YVMYWYWTGNNW-----TDCT | 416 | 1.71% |  |
| 14 | TLCV-----YVMYWYWTGNNW-----IACQGTG | 390 | 1.60% |  |
| 15 | TLCI-----YVMYWYWTGNNW-----IDCSGTG | 364 | 1.49% |  |
| 16 | TLCL-----YVMYWYWTGNNW-----FDCHTG | 357 | 1.46% |  |
| 17 | TLCV-----YVMYWYWTGNNW-----IPCQGTG | 352 | 1.44% |  |
| 18 | TLCL-----YVMYWYWTGNNW-----THCQGTG | 327 | 1.34% |  |
| 19 | TLVYE-----YVMYWYWTGNNW-----IPMRTG | 312 | 1.28% |  |
| 20 | TLCM-----YVMYWYWTGNNW-----VDCHGTG | 303 | 1.24% |  |
| 21 | TLCL-----YVMYWYWTGNNW-----VPCQGTG | 299 | 1.23% |  |
| 22 | TLCL-----YVMYWYWTGNNW-----IDCWTG | 287 | 1.18% |  |
| 23 | TLCI-----YVMYWYWTGNNW-----VNCTG | 252 | 1.03% |  |
| **24** | **TLYH-----YVMYWYWTGNNW-----LLGNRTG** | **246** | **1.01%** | **LG2-4** |
| 25 | TLCL-----YVMYWYWTGNNW-----IDCFTG | 238 | 0.98% |  |
| **26** | **TLW-----YVMYWYWTGNNW-----ITGIHTG** | **235** | **0.96%** | **LG2-5** |
| 27 | TLW-----YVMYWYWTGNNW-----ITGMHTG | 229 | 0.94% |  |
| 28 | TLCM-----YVMYWYWTGNNW-----VDCTG | 223 | 0.91% |  |
| 29 | TLW-----YVMYWYWTGNNW-----ITGLHTG | 220 | 0.90% |  |
| 30 | TLCM-----YVMYWYWTGNNW-----VPCHTG | 178 | 0.73% |  |
| 31 | TLCM-----YVMYWYWTGNNW-----FDCHTG | 169 | 0.69% |  |
| 32 | TLCM-----YVMYWYWTGNNW-----VPCHETG | 167 | 0.69% |  |
| 33 | TLCL-----YVMYWYWTGNNW-----IDCYTG | 161 | 0.66% |  |
| 34 | TLWPDGS-----YVMYWYWTGNNW-----LLTTG | 151 | 0.62% |  |
| 35 | TLCI-----YVMYWYWTGNNW-----VDCHTG | 150 | 0.62% |  |
| 36 | TLFDN-----YVMYWYWTGNNW-----FPVTTG | 141 | 0.58% |  |
| 37 | TLCL-----YVMYWYWTGNNW-----VDCHTG | 140 | 0.57% |  |
| 38 | TLDG-----YVMYWYWTGNNW-----IPLWTG | 126 | 0.52% |  |
| 39 | TLSGC-----YVMYWYWTGNNW-----IPMCTG | 122 | 0.50% |  |
| 40 | TLQCYFN-----YVMYWYWTGNNW-----VS | 119 | 0.49% |  |
| 41 | TLYDN-----YVMYWYWTGNNW-----SVQTTG | 118 | 0.48% |  |
| 42 | TLCL-----YVMYWYWTGNNW-----VDCSGTG | 113 | 0.46% |  |
| 43 | TLVG-----YVMYWYWTGNNW-----IPLWTG | 111 | 0.46% |  |
| 44 | TLCL-----YVMYWYWTGNNW-----IDCTG | 107 | 0.44% |  |
| 45 | TLCL-----YVMYWYWTGNNW-----TDLCTG | 107 | 0.44% |  |
| 46 | TLCL-----YVMYWYWTGNNW-----VPCNGTG | 106 | 0.43% |  |
| 47 | TLCL-----YVMYWYWTGNNW-----TGCYPTG | 101 | 0.41% |  |
| 48 | TLCL-----YVMYWYWTGNNW-----VDCYGTG | 98 | 0.40% |  |
| 49 | TLCI-----YVMYWYWTGNNW-----VDCSGTG | 94 | 0.39% |  |
| 50 | TLVMP-----YVMYWYWTGNNW-----IPMRTG | 88 | 0.36% |  |
| 51 | TLFDN-----YVMYWYWTGNNW-----IPITTG | 85 | 0.35% |  |
| 52 | TLCM-----YVMYWYWTGNNW-----TDVCQTG | 79 | 0.32% |  |
| 53 | TLCL-----YVMYWYWTGNNW-----VDCATG | 79 | 0.32% |  |
| 54 | TLWSLP-----YVMYWYWTGNNW-----LITRNTG | 74 | 0.30% |  |
| 55 | TLCV-----YVMYWYWTGNNW-----VQCQGTG | 73 | 0.30% |  |
| 56 | TLCL-----YVMYWYWTGNNW-----VDCFTG | 73 | 0.30% |  |
| 57 | TLCL-----YVMYWYWTGNNW-----VDCADTG | 70 | 0.29% |  |
| 58 | TLCI-----YVMYWYWTGNNW-----IDCFTG | 68 | 0.28% |  |
| 59 | TLCM-----YVMYWYWTGNNW-----SDVCTG | 62 | 0.25% |  |
| 60 | TLYDAGW-----YVMYWYWTGNNW-----IPMKNTG | 61 | 0.25% |  |
| 61 | TLCL-----YVMYWYWTGNNW-----VPCT | 60 | 0.25% |  |
| 62 | TLCL-----YVMYWYWTGNNW-----VPCHGTG | 56 | 0.23% |  |
| 63 | TLW-----YVMYWYWTGNNW-----ISGLHTG | 52 | 0.21% |  |
| 64 | TLCL-----YVMYWYWTGNNW-----APCQGTG | 51 | 0.21% |  |
| 65 | TLCI-----YVMYWYWTGNNW-----IDCYTG | 49 | 0.20% |  |
| 66 | TLCM-----YVMYWYWTGNNW-----IDCTG | 45 | 0.18% |  |
| 67 | TLCV-----YVMYWYWTGNNW-----THCQGTG | 45 | 0.18% |  |
| 68 | TLCM-----YVMYWYWTGNNW-----VECHATG | 43 | 0.18% |  |
| 69 | TLCL-----YVMYWYWTGNNW-----VDCWNTG | 42 | 0.17% |  |
| 70 | TLVG-----YVMYWYWTGNNW-----IPMWTG | 39 | 0.16% |  |

| | | | |
|---|---|---|---|
| 71 | TLCL-----YVMYWYWTGNNW-----FDCTG | 85 | 0.42% |
| 72 | TLCM-----YVMYWYWTGNNW-----TDVCQTG | 84 | 0.42% |
| 73 | TLDVMYWYWTGNNWRPATG | 83 | 0.41% |
| 74 | TLCL-----YVMYWYWTGNNW-----VDCWNTG | 80 | 0.40% |
| 75 | TLCV-----YVMYWYWTGNNW-----VDCYTG | 79 | 0.39% |
| 76 | TLF-----YVMYWYWTGNNW-----ITGIHTG | 77 | 0.38% |
| 77 | TLCV-----YVMYWYWTGNNW-----AKCT | 76 | 0.38% |
| 78 | TLVSE-----YVMYWYWTGNNW-----IPMRTG | 76 | 0.38% |
| 79 | TLCM-----YVMYWYWTGNNW-----SDVCTG | 74 | 0.37% |
| 80 | TLCM-----YVMYWYWTGNNW-----VQCHATG | 71 | 0.35% |
| 81 | TLFH-----YVMYWYWTGNNW-----ILGNRTG | 70 | 0.35% |
| 82 | TLCI-----YVMYWYWTGNNW-----VNCTG | 69 | 0.34% |
| 83 | TLHCC-----YVMYWYWTGNNW-----IPMWTG | 68 | 0.34% |
| 84 | TLVG-----YVMYWYWTGNNW-----IPMRTG | 67 | 0.33% |
| 85 | TLW-----YVMYWYWTGNNW-----ITGMHTG | 67 | 0.33% |
| 86 | TLHQ-----YVMYWYWTGNNW-----LPMWTG | 67 | 0.33% |
| 87 | TLCY-----YVMYWYWTGNNW-----VDCTG | 67 | 0.33% |
| 88 | TLCM-----YVMYWYWTGNNW-----FNCRT | 66 | 0.33% |
| 89 | TLHQ-----YVMYWYWTGNNW-----IPLWTG | 65 | 0.32% |
| 90 | TLMQ-----YVMYWYWTGNNW-----IPMWTG | 63 | 0.31% |
| 91 | TLWSLP-----YVMYWYWTGNNW-----IKTRETG | 61 | 0.30% |
| 92 | TLCL-----YVMYWYWTGNNW-----VPCHTG | 61 | 0.30% |
| 93 | TLFH-----YVMYWYWTGNNW-----LLGNRTG | 58 | 0.29% |
| 94 | TLYDN-----YVMYWYWTGNNW-----CPITTG | 57 | 0.28% |
| 95 | TSFVYD-----YVMYWYWTGNNW-----ILMPTG | 56 | 0.28% |
| 96 | TLCM-----YVMYWYWTGNNW-----IDCWTG | 56 | 0.28% |
| 97 | TLCI-----YVMYWYWTGNNW-----FDCTG | 56 | 0.28% |
| 98 | TLCI-----YVMYWYWTGNNW-----SDVCTG | 54 | 0.27% |
| 99 | TLCV-----YVMYWYWTGNNW-----TSCQGTG | 53 | 0.26% |
| 100 | TLSQ-----YVMYWYWTGNNW-----MPMWTG | 53 | 0.26% |
| | *total reads (top 100)* | 20162 | |

| | | | |
|---|---|---|---|
| 71 | TLCI-----YVMYWYWTGNNW-----IDCFTG | 42 | 0.26% |
| 72 | TLCM-----YVMYWYWTGNNW-----TDVCQTG | 41 | 0.25% |
| 73 | TLCM-----YVMYWYWTGNNW-----SDVCTG | 41 | 0.25% |
| 74 | TLCL-----YVMYWYWTGNNW-----TDLCTG | 40 | 0.25% |
| 75 | TLCI-----YVMYWYWTGNNW-----FDCTG | 39 | 0.24% |
| 76 | TLYTRSL-----YVMYWYWTGNNW-----MGSTG | 38 | 0.23% |
| 77 | TLCI-----YVMYWYWTGNNW-----SDICTG | 38 | 0.23% |
| 78 | TLYDAGW-----YVMYWYWTGNNW-----IPMKNTG | 38 | 0.23% |
| 79 | TLWSLP-----YVMYWYWTGNNW-----LITRNTG | 38 | 0.23% |
| 80 | TLCL-----YVMYWYWTGNNW-----APCQGTG | 36 | 0.22% |
| 81 | TLVG-----YVMYWYWTGNNW-----FPMWTG | 36 | 0.22% |
| 82 | TLCM-----YVMYWYWTGNNW-----IDCWTG | 35 | 0.21% |
| 83 | TLCI-----YVMYWYWTGNNW-----IDCYTG | 33 | 0.20% |
| 84 | TLCV-----YVMYWYWTGNNW-----TSCQGTG | 33 | 0.20% |
| 85 | TLCV-----YVMYWYWTGNNW-----VDCYTG | 32 | 0.20% |
| 86 | TLCT-----YVMYWYWTGNNW-----VPCQGTG | 32 | 0.20% |
| 87 | TLW-----YVMYWYWTGNNW-----ISGLHTG | 32 | 0.20% |
| 88 | TLSGC-----YVMYWYWTGNNW-----IPMCTG | 31 | 0.19% |
| 89 | TLL-----YVMYWYWTGNNW-----LPIFRTG | 29 | 0.18% |
| 90 | TLCL-----YVMYWYWTGNNW-----VPCHNTG | 28 | 0.17% |
| 91 | TLVNW-----YVMYWYWTGNNW-----ALTG | 28 | 0.17% |
| 92 | TLCM-----YVMYWYWTGNNW-----VQCHATG | 27 | 0.17% |
| 93 | TLCI-----YVMYWYWTGNNW-----SDVCTG | 26 | 0.16% |
| 94 | TLCV-----YVMYWYWTGNNW-----VPCQATG | 25 | 0.15% |
| 95 | TLCM-----YVMYWYWTGNNW-----VECHATG | 25 | 0.15% |
| 96 | TLCM-----YVMYWYWTGNNW-----FNCRT | 24 | 0.15% |
| 97 | TLYP-----YVMYWYWTGNNW-----LLGNRTG | 24 | 0.15% |
| 98 | TLYDN-----YVMYWYWTGNNW-----EIITTG | 24 | 0.15% |
| 99 | TLCM-----YVMYWYWTGNNW-----IPCHHTG | 24 | 0.15% |
| 100 | TLWPDGS-----YVMYWYWTGNNW-----LLTTG | 24 | 0.15% |
| | *total reads (top 100)* | 16323 | |

| | | | |
|---|---|---|---|
| 71 | TLCV-----YVMYWYWTGNNW-----VDCNTG | 38 | 0.16% |
| 72 | TLW-----YVMYWYWTGNNW-----IQGIHTG | 38 | 0.16% |
| 73 | TLW-----YVMYWYWTGNNW-----IEGLHTG | 37 | 0.15% |
| 74 | TLDDSH-----YVMYWYWTGNNW-----IPMRTG | 35 | 0.14% |
| 75 | TLCM-----YVMYWYWTGNNW-----VGCHATG | 34 | 0.14% |
| 76 | TLCI-----YVMYWYWTGNNW-----SDVCTG | 34 | 0.14% |
| 77 | TLCM-----YVMYWYWTGNNW-----IDCFTG | 31 | 0.13% |
| 78 | TLVSE-----YVMYWYWTGNNW-----IPMRTG | 31 | 0.13% |
| 79 | TLCL-----YVMYWYWTGNNW-----FDCTG | 29 | 0.12% |
| 80 | TLVG-----YVMYWYWTGNNW-----IPMLTG | 29 | 0.12% |
| 81 | TLWSLP-----YVMYWYWTGNNW-----IKTRETG | 28 | 0.11% |
| 82 | TLCI-----YVMYWYWTGNNW-----SDICTG | 28 | 0.11% |
| 83 | TLWSVMYWYWTGNNWITGMHTG | 27 | 0.11% |
| 84 | TLYP-----YVMYWYWTGNNW-----LLGNRTG | 27 | 0.11% |
| 85 | TLCM-----YVMYWYWTGNNW-----VQCHATG | 27 | 0.11% |
| 86 | TLWSVMYWYWTGNNWITGLHTG | 27 | 0.11% |
| 87 | TLSQ-----YVMYWYWTGNNW-----MPMWTG | 26 | 0.11% |
| 88 | TLCL-----YVMYWYWTGNNW-----MDCTG | 26 | 0.11% |
| 89 | TLWPDFS-----YVMYWYWTGNNW-----FPTTG | 26 | 0.11% |
| 90 | TLCL-----YVMYWYWTGNNW-----VGCWPTG | 26 | 0.11% |
| 91 | TLW-----YVMYWYWTGNNW-----INGIHTG | 26 | 0.11% |
| 92 | TLHCSPC-----YVMYWYWTGNNW-----AFTG | 25 | 0.10% |
| 93 | TLCI-----YVMYWYWTGNNW-----FDCTG | 25 | 0.10% |
| 94 | TLHQ-----YVMYWYWTGNNW-----LPMWTG | 25 | 0.10% |
| 95 | TLCM-----YVMYWYWTGNNW-----IDCWTG | 25 | 0.10% |
| 96 | TLCM-----YVMYWYWTGNNW-----FDCHGTG | 24 | 0.10% |
| 97 | TLFH-----YVMYWYWTGNNW-----VLGNRTG | 24 | 0.10% |
| 98 | TLCI-----YVMYWYWTGNNW-----FDCHTG | 24 | 0.10% |
| 99 | TLCM-----YVMYWYWTGNNW-----IPCHHTG | 24 | 0.10% |
| 100 | TLCM-----YVMYWYWTGNNW-----TDVCTG | 23 | 0.09% |
| | *total reads (top 100)* | 24377 | |

### Supporting Information Table S1

#### Page 4 - ZeroStart library against GαS

Top 100 sequences from ZeroStart library against GαS in different selection rounds, with NGS read frequency. Bold entries indicate the sequences selected for further analysis.
The Ubiquitin core residues around the randomized loop 1 insert are ommitted for clarity. They span Ub1-6 and Ub11-76

| Round 3 | | | | |
|---|---|---|---|---|
| Rank | Loop 1 | read count | read frequency | U-body name |
| **1** | **IGAFVLVWPWSPG** | **3263** | **30.77%** | **RL2-1** |
| **2** | **GVDLFCGVYWDCM** | **889** | **8.38%** | **RL2-3** |
| **3** | **LGRLWLPYPFDLE** | **614** | **5.79%** | **RL2-2** |
| **4** | **WLVECVGFKTCFVF** | **548** | **5.17%** | **RL2-8** |
| **5** | **AASWVFIYPSK** | **500** | **4.71%** | **RL2-6** |
| **6** | **LEWWAYFWGF** | **491** | **4.63%** | **RL2-5** |
| **7** | **FLVPIRRFRNALVF** | **490** | **4.62%** | **RL2-4** |
| **8** | **SLVKCTGFRWCLTW** | **434** | **4.09%** | **RL2-7** |
| **9** | **VPPLVLMWPWDSG** | **229** | **2.16%** | **RL2-10** |
| 10 | MYLTWFGVYAP | 204 | 1.92% | |
| 11 | AQRWVFVWSGSWR | 166 | 1.57% | |
| 12 | ASHGWPWLSLPGS | 153 | 1.44% | |
| **13** | **WIFWAWVLDVLGR** | **143** | **1.35%** | **RL2-9** |
| 14 | WLDSSWIWAAVGIVV | 132 | 1.24% | |
| 15 | LLFPDMWKFILFW | 127 | 1.20% | |
| 16 | DVLGLDEYAYFVV | 123 | 1.16% | |
| 17 | YLVTVVGFLYSLVF | 121 | 1.14% | |
| 18 | LWYHSFVHCLCD | 97 | 0.91% | |
| 19 | LWLSMDTFMLIFDRV | 87 | 0.82% | |
| 20 | WFFPWFLVRSW | 84 | 0.79% | |
| 21 | WSFWIWVATFVST | 76 | 0.72% | |
| 22 | VECICIWPFMNG | 67 | 0.63% | |
| 23 | SWPFYFLLV | 63 | 0.59% | |
| 24 | MWECGVVTPIWGCAD | 59 | 0.56% | |
| 25 | FLVPCSYFRWCLVY | 56 | 0.53% | |
| 26 | VYLLRLPGLVYRLT | 49 | 0.46% | |
| 27 | ARFTFGGVWAP | 49 | 0.46% | |
| 28 | LLPCGWDWIVFT | 48 | 0.45% | |
| 29 | GEAASESWGWTVRYL | 45 | 0.42% | |
| 30 | VRSYVLVWDGVWR | 42 | 0.40% | |
| 31 | PIPYVLVHPH | 41 | 0.39% | |
| 32 | PALLSYYGIWMSGWF | 40 | 0.38% | |
| 33 | PSRFILIWPF | 36 | 0.34% | |
| 34 | PCFLRVGAR | 35 | 0.33% | |
| 35 | PDVYWTGGFTPFFWW | 31 | 0.29% | |
| 36 | IMPYGVWPWLWW | 30 | 0.28% | |
| 37 | RCWLSVGVFG | 27 | 0.25% | |
| 38 | LLERCWTLIVFP | 20 | 0.19% | |
| 39 | AIRWVLIYPW | 20 | 0.19% | |
| 40 | LCDFRMWWLLLFM | 19 | 0.18% | |
| 41 | WNRAHRFP | 19 | 0.18% | |
| 42 | YFLRLYFESVYAIW | 18 | 0.17% | |
| 43 | ALGVGWWLPA | 18 | 0.17% | |
| 44 | LVYCVVYWFRWESSG | 18 | 0.17% | |
| 45 | QALFTLLQLLSGA | 18 | 0.17% | |
| 46 | SSYLCFRLATLFQS | 18 | 0.17% | |
| 47 | RWTTHWFLMLF | 17 | 0.16% | |
| 48 | VVLADVASLIGSGC | 17 | 0.16% | |
| 49 | SSIIVFSGHIYA | 17 | 0.16% | |
| 50 | FALGRYGACVLPRQ | 16 | 0.15% | |
| 51 | VLGWQFCEWCT | 16 | 0.15% | |
| 52 | RMRSHERNVTGC | 15 | 0.14% | |
| 53 | RGSCSFVSCCMC | 15 | 0.14% | |
| 54 | LMRLDFHRVSM | 15 | 0.14% | |
| 55 | LLWRAVLMRAS | 15 | 0.14% | |
| 56 | PATRGWHLILFG | 15 | 0.14% | |
| 57 | TGRHSCCLCLWRR | 15 | 0.14% | |
| 58 | YCPHVGNCM | 15 | 0.14% | |
| 59 | ILVEKYFE | 15 | 0.14% | |
| 60 | GVFFLPRREL | 15 | 0.14% | |
| 61 | VAVLLLVICWPRS | 15 | 0.14% | |
| 62 | CVSCPWFLY | 15 | 0.14% | |

| Round 4 | | | | |
|---|---|---|---|---|
| Rank | Loop 1 | read count | read frequency | U-body name |
| **1** | **IGAFVLVWPWSPG** | **44136** | **50.30%** | **RL2-1** |
| **2** | **GVDLFCGVYWDCM** | **10783** | **12.29%** | **RL2-3** |
| **3** | **LGRLWLPYPFDLE** | **7403** | **8.44%** | **RL2-2** |
| **4** | **LEWWAYFWGF** | **4008** | **4.57%** | **RL2-5** |
| **5** | **FLVPIRRFRNALVF** | **3623** | **4.13%** | **RL2-4** |
| **6** | **AASWVFIYPSK** | **3212** | **3.66%** | **RL2-6** |
| **7** | **SLVKCTGFRWCLTW** | **2576** | **2.94%** | **RL2-7** |
| **8** | **WLVECVGFKTCFVF** | **2540** | **2.89%** | **RL2-8** |
| **9** | **WIFWAWVLDVLGR** | **1119** | **1.28%** | **RL2-9** |
| 10 | WLDSSWIWAAVGIVV | 914 | 1.04% | |
| 11 | MYLTWFGVYAP | 767 | 0.87% | |
| 12 | ASHGWPWLSLPGS | 765 | 0.87% | |
| 13 | DVLGLDEYAYFVV | 711 | 0.81% | |
| 14 | AQRWVFVWSGSWR | 676 | 0.77% | |
| 15 | LLFPDMWKFILFW | 671 | 0.76% | |
| **16** | **VPPLVLMWPWDSG** | **607** | **0.69%** | **RL2-10** |
| 17 | LWYHSFVHCLCD | 303 | 0.35% | |
| 18 | WSFWIWVATFVST | 271 | 0.31% | |
| 19 | YLVTVVGFLYSLVF | 246 | 0.28% | |
| 20 | WFFPWFLVRSW | 246 | 0.28% | |
| 21 | VECICIWPFMNG | 197 | 0.22% | |
| 22 | LLPCGWDWIVFT | 158 | 0.18% | |
| 23 | MWECGVVTPIWGCAD | 154 | 0.18% | |
| 24 | LWLSMDTFMLIFDRV | 138 | 0.16% | |
| 25 | FLVPCSYFRWCLVY | 112 | 0.13% | |
| 26 | VRSYVLVWDGVWR | 85 | 0.10% | |
| 27 | VYLLRLPGLVYRLT | 81 | 0.09% | |
| 28 | PSRFILIWPF | 64 | 0.07% | |
| 29 | IMPYGVWPWLWW | 50 | 0.06% | |
| 30 | RCWLSVGVFG | 48 | 0.05% | |
| 31 | PDVYWTGGFTPFFWW | 41 | 0.05% | |
| 32 | IRAFVLVWPWSPG | 39 | 0.04% | |
| 33 | PCFLRVGAR | 38 | 0.04% | |
| 34 | IGAFVLIWPWSPG | 37 | 0.04% | |
| 35 | GEAASESWGWTVRYL | 37 | 0.04% | |
| 36 | ARFTFGGVWAP | 36 | 0.04% | |
| 37 | PALLSYYGIWMSGWF | 35 | 0.04% | |
| 38 | PIPYVLVHPH | 34 | 0.04% | |
| 39 | VPSRVLLYPHDSG | 34 | 0.04% | |
| 40 | IGAFVLVWPWSPS | 33 | 0.04% | |
| 41 | VLGWQFCEWCT | 29 | 0.03% | |
| 42 | IEAFVLVWPWSPG | 28 | 0.03% | |
| 43 | IGVFVLVWPWSPG | 27 | 0.03% | |
| 44 | LCDFRMWWLLLFM | 26 | 0.03% | |
| 45 | IGALVLVWPWSPG | 24 | 0.03% | |
| 46 | IGAFVLVWPWSPD | 23 | 0.03% | |
| 47 | IGAFILVWPWSPG | 22 | 0.03% | |
| 48 | IGAFVLVWPRSPG | 22 | 0.03% | |
| 49 | IGAFVLVWPW | 21 | 0.02% | |
| 50 | LERGWTCIFFW | 19 | 0.02% | |
| 51 | LLERCWTLIVFP | 18 | 0.02% | |
| 52 | LRYPRSWLWILFW | 18 | 0.02% | |
| 53 | IGTFVLVWPWSPG | 18 | 0.02% | |
| 54 | IGAFVLVWTWSPG | 15 | 0.02% | |
| 55 | IPGLVSPYPFDRA | 15 | 0.02% | |
| 56 | AIRWVLIYPW | 14 | 0.02% | |
| 57 | IGAFVLVCPWSPG | 14 | 0.02% | |
| 58 | IGRLWLPYPFDLE | 13 | 0.01% | |
| 59 | IGAFVLVWLWSPG | 13 | 0.01% | |
| 60 | IGAFVLVWPWYPG | 12 | 0.01% | |
| 61 | SVDLFCGVYWDCM | 11 | 0.01% | |
| 62 | CLWMCMLCGLYLSV | 11 | 0.01% | |

| Round 5 | | | | |
|---|---|---|---|---|
| Rank | Loop 1 | read count | read frequency | U-body name |
| **1** | **IGAFVLVWPWSPG** | **46834** | **62.34%** | **RL2-1** |
| **2** | **LGRLWLPYPFDLE** | **8902** | **11.85%** | **RL2-2** |
| **3** | **GVDLFCGVYWDCM** | **7932** | **10.56%** | **RL2-3** |
| **4** | **FLVPIRRFRNALVF** | **2677** | **3.56%** | **RL2-4** |
| **5** | **LEWWAYFWGF** | **1696** | **2.26%** | **RL2-5** |
| **6** | **AASWVFIYPSK** | **1402** | **1.87%** | **RL2-6** |
| **7** | **SLVKCTGFRWCLTW** | **1134** | **1.51%** | **RL2-7** |
| **8** | **WLVECVGFKTCFVF** | **1017** | **1.35%** | **RL2-8** |
| **9** | **WIFWAWVLDVLGR** | **604** | **0.80%** | **RL2-9** |
| **10** | **VPPLVLMWPWDSG** | **387** | **0.52%** | **RL2-10** |
| 11 | WLDSSWIWAAVGIVV | 339 | 0.45% | |
| 12 | LLFPDMWKFILFW | 291 | 0.39% | |
| 13 | DVLGLDEYAYFVV | 283 | 0.38% | |
| 14 | ASHGWPWLSLPGS | 255 | 0.34% | |
| 15 | AQRWVFVWSGSWR | 152 | 0.20% | |
| 16 | MYLTWFGVYAP | 134 | 0.18% | |
| 17 | LWYHSFVHCLCD | 68 | 0.09% | |
| 18 | WSFWIWVATFVST | 62 | 0.08% | |
| 19 | WFFPWFLVRSW | 54 | 0.07% | |
| 20 | IGAFILVWPWSPG | 47 | 0.06% | |
| 21 | IGAFVLIWPWSPG | 41 | 0.05% | |
| 22 | YLVTVVGFLYSLVF | 38 | 0.05% | |
| 23 | MWECGVVTPIWGCAD | 36 | 0.05% | |
| 24 | VECICIWPFMNG | 35 | 0.05% | |
| 25 | IGTFVLVWPWSPG | 31 | 0.04% | |
| 26 | IEAFVLVWPWSPG | 28 | 0.04% | |
| 27 | IGAFVLVWPWSPS | 24 | 0.03% | |
| 28 | LLPCGWDWIVFT | 24 | 0.03% | |
| 29 | FLVPCSYFRWCLVY | 22 | 0.03% | |
| 30 | IGVFVLVWPWSPG | 21 | 0.03% | |
| 31 | IRAFVLVWPWSPG | 19 | 0.03% | |
| 32 | IGAFVLVWPWSPD | 19 | 0.03% | |
| 33 | LWLSMDTFMLIFDRV | 16 | 0.02% | |
| 34 | VGAFVLVWPWSPG | 16 | 0.02% | |
| 35 | IGAFVLVWPWSSG | 16 | 0.02% | |
| 36 | PSRFILIWPF | 15 | 0.02% | |
| 37 | GVDLFCGVYWDCV | 14 | 0.02% | |
| 38 | VYLLRLPGLVYRLT | 13 | 0.02% | |
| 39 | IGSFVLVWPWSPG | 13 | 0.02% | |
| 40 | IGAFVLVWPRSPG | 13 | 0.02% | |
| 41 | IGAFVFVWPWSPG | 13 | 0.02% | |
| 42 | IGALVLVWPWSPG | 12 | 0.02% | |
| 43 | IGAFVLVCPWSPG | 12 | 0.02% | |
| 44 | IGAFVLVWPW | 11 | 0.01% | |
| 45 | IGAFVLVW | 11 | 0.01% | |
| 46 | GVDLFCGVYWDCI | 11 | 0.01% | |
| 47 | IGAFVLVWSWSPG | 11 | 0.01% | |
| 48 | VRSYVLVWDGVWR | 10 | 0.01% | |
| 49 | LGRLWLPYPFDLG | 10 | 0.01% | |
| 50 | IMPYGVWPWLWW | 9 | 0.01% | |
| 51 | IGDFVLVWPWSPG | 9 | 0.01% | |
| 52 | IGAFVLVWPLSPG | 9 | 0.01% | |
| 53 | IGAFVLVWTWSPG | 8 | 0.01% | |
| 54 | IGAFVLVWPWSLG | 8 | 0.01% | |
| 55 | IVAFVLVWPWSPG | 8 | 0.01% | |
| 56 | GVDLFCGIYWDCM | 8 | 0.01% | |
| 57 | LGQLWLPYPFDLE | 8 | 0.01% | |
| 58 | IGAFVLVRPWSPG | 7 | 0.01% | |
| 59 | GVDLFCGVYWDCL | 7 | 0.01% | |
| 60 | IGAFVLVWPWSPV | 7 | 0.01% | |
| 61 | VPSRVLLYPHDSG | 7 | 0.01% | |
| 62 | IGAFVLVWPWSTG | 7 | 0.01% | |

| | | | |
|---|---|---|---|
| 63 | RVSWLKAC | 14 | 0.13% |
| 64 | STLVSYFESVYAIW | 14 | 0.13% |
| 65 | MAFRLYFESV | 14 | 0.13% |
| 66 | CRCVSRWFFDALCW | 14 | 0.13% |
| 67 | SFLATYFESVTQFG | 14 | 0.13% |
| 68 | VPCFSSAVLL | 14 | 0.13% |
| 69 | LRHVLRMFWRY | 14 | 0.13% |
| 70 | YVEDLMRYCWL | 14 | 0.13% |
| 71 | VLLYSYPPSSR | 14 | 0.13% |
| 72 | LPCFCIERLDASV | 14 | 0.13% |
| 73 | LFFDRGRLLLVC | 14 | 0.13% |
| 74 | LATFWLGKLCGT | 14 | 0.13% |
| 75 | WAHSAYFES | 13 | 0.12% |
| 76 | CSFGFSSL | 13 | 0.12% |
| 77 | GKSVVVLWKLCHFRC | 13 | 0.12% |
| 78 | SSPFRWVMSSL | 13 | 0.12% |
| 79 | GGGIVLKLSSYNLRL | 13 | 0.12% |
| 80 | IRCFTPRDR | 13 | 0.12% |
| 81 | LFFPGPYVASGFLV | 13 | 0.12% |
| 82 | AHWIALRLLTC | 13 | 0.12% |
| 83 | LPHLVEASCF | 13 | 0.12% |
| 84 | RSWFRALRAV | 13 | 0.12% |
| 85 | DYLRFGILYAGV | 13 | 0.12% |
| 86 | RWSCVYFESVYAIW | 13 | 0.12% |
| 87 | WLGPLMVR | 13 | 0.12% |
| 88 | FFACWLRWFL | 13 | 0.12% |
| 89 | ENVHLWYLSDML | 13 | 0.12% |
| 90 | PAWLVMCMG | 13 | 0.12% |
| 91 | WASSAQADEFF | 13 | 0.12% |
| 92 | CGCSEVSVSLGL | 13 | 0.12% |
| 93 | IGPIYYFESVY | 13 | 0.12% |
| 94 | IFWWFLVGP | 13 | 0.12% |
| 95 | LAPGLSPC | 12 | 0.11% |
| 96 | RLHCIVYFRM | 12 | 0.11% |
| 97 | VSDKWYFESVY | 12 | 0.11% |
| 98 | SRLLVVTRF | 12 | 0.11% |
| 99 | LRGKAERVRS | 12 | 0.11% |
| 100 | SQCAPVFPLLRGVVA | 12 | 0.11% |
| | *total reads (top 100)* | 10605 | |

| | | | |
|---|---|---|---|
| 63 | LGAFVLVWPWSPG | 11 | 0.01% |
| 64 | IFWWFLVGP | 11 | 0.01% |
| 65 | IGAFVLVWPWSSG | 11 | 0.01% |
| 66 | IVAFVLVWPWSPG | 11 | 0.01% |
| 67 | GVDLFCGVYWDCI | 10 | 0.01% |
| 68 | IGAFVLVWPWSLG | 10 | 0.01% |
| 69 | IGAFVLVW | 10 | 0.01% |
| 70 | IGAFVLVWPCSPG | 10 | 0.01% |
| 71 | IGAFVLVWSWSPG | 10 | 0.01% |
| 72 | IGAFVLVWPWFPG | 10 | 0.01% |
| 73 | IGDFVLVWPWSPG | 10 | 0.01% |
| 74 | IGAFVLVWHWSPG | 9 | 0.01% |
| 75 | GVDLFCGVYWDYM | 9 | 0.01% |
| 76 | IGAFALVWPWSPG | 9 | 0.01% |
| 77 | GVDLFCGIYWDCM | 9 | 0.01% |
| 78 | TSEWVLIYRGERG | 8 | 0.01% |
| 79 | VGAFVLVWPWSPG | 8 | 0.01% |
| 80 | FGAFVLVWPWSPG | 8 | 0.01% |
| 81 | GMDLFCGVYWDCM | 7 | 0.01% |
| 82 | GVDLFCGVYWDCV | 7 | 0.01% |
| 83 | IGAFVLVRPWSPG | 7 | 0.01% |
| 84 | DVDLFCGVYWDCM | 7 | 0.01% |
| 85 | LGQLWLPYPFDLE | 7 | 0.01% |
| 86 | IGSFVLVWPWSPG | 7 | 0.01% |
| 87 | IGAFVLVLPWSPG | 7 | 0.01% |
| 88 | LGSSAMWRWLLFF | 7 | 0.01% |
| 89 | IGAFVLFWPWSPG | 6 | 0.01% |
| 90 | TGAFVLVWPWSPG | 6 | 0.01% |
| 91 | GVNLFCGVYWDCM | 6 | 0.01% |
| 92 | HFVTIWWSFSHALGW | 6 | 0.01% |
| 93 | LGRLWLPYPFDLG | 6 | 0.01% |
| 94 | IGAFVLGWPWSPG | 6 | 0.01% |
| 95 | IGAFVLVWPWPPG | 6 | 0.01% |
| 96 | IWAFVLVWPWSPG | 6 | 0.01% |
| 97 | IGAFVLAWPWSPG | 6 | 0.01% |
| 98 | IGAFVFVWPWSPG | 6 | 0.01% |
| 99 | PATRGWHLILFG | 5 | 0.01% |
| 100 | IGAFVLVWPWSP | 5 | 0.01% |
| | *total reads (top 100)* | 87741 | |

| | | | |
|---|---|---|---|
| 63 | IGRLWLPYPFDLE | 7 | 0.01% |
| 64 | PIPYVLVHPH | 7 | 0.01% |
| 65 | IGAFVLFWPWSPG | 7 | 0.01% |
| 66 | PCFLRVGAR | 6 | 0.01% |
| 67 | IGAFVLAWPWSPG | 6 | 0.01% |
| 68 | LSRLWLPYPFDLE | 6 | 0.01% |
| 69 | GVDLFCGVYWDYM | 6 | 0.01% |
| 70 | LGRLWLPYPFDSE | 6 | 0.01% |
| 71 | LERGWTCIFFW | 6 | 0.01% |
| 72 | SLVECTGFRWCLTW | 6 | 0.01% |
| 73 | IGAFVLGWPWSPG | 6 | 0.01% |
| 74 | IGAFVLVLPWSPG | 6 | 0.01% |
| 75 | IGAFVLVWPWFPG | 6 | 0.01% |
| 76 | IGAFVLVWPCSPG | 6 | 0.01% |
| 77 | RCWLSVGVFG | 6 | 0.01% |
| 78 | IGAFVLVWLWSPG | 6 | 0.01% |
| 79 | FLVPIRRFHNALVF | 5 | 0.01% |
| 80 | GVDLFCSVYWDCM | 5 | 0.01% |
| 81 | LGRLWLPYPF | 5 | 0.01% |
| 82 | LGRLWLPYLFDLE | 5 | 0.01% |
| 83 | GVDLFCGVYWNCM | 5 | 0.01% |
| 84 | LGRLWLPY | 5 | 0.01% |
| 85 | ARFTFGGVWAP | 5 | 0.01% |
| 86 | IGAFVLVWPWPPG | 5 | 0.01% |
| 87 | LGRPWLPYPFDLE | 5 | 0.01% |
| 88 | IGAFVLVWPWYPG | 5 | 0.01% |
| 89 | IGAFVLVWHWSPG | 5 | 0.01% |
| 90 | IGAFVLVWPWSHG | 4 | 0.01% |
| 91 | FLVPIRRFRNALVL | 4 | 0.01% |
| 92 | FLVPIRRFRNALMF | 4 | 0.01% |
| 93 | LGRLWLLYPFDLE | 4 | 0.01% |
| 94 | LGAFVLVWPWSPG | 4 | 0.01% |
| 95 | LGRLWLPYPFNLE | 4 | 0.01% |
| 96 | GVDLFCGV | 4 | 0.01% |
| 97 | LGRLWLPYPFDLK | 4 | 0.01% |
| 98 | PALLSYYGIWMSGWF | 4 | 0.01% |
| 99 | IGAYVLVWPWSPG | 4 | 0.01% |
| 100 | IGAFALVWPWSPG | 3 | 0.00% |
| | *total reads (top 100)* | 75124 | |

Supporting Information Table S1

**Page 5 - Lasso-graft library LG3 against GαS**

Top 100 sequences from Lasso-Graft library LG against GαS in different selection rounds, with NGS read frequency. Bold entries indicate the sequences selected for further analysis.
The grafted sequence is surrounded by dashes for clarity. Red text indicates mutations occurred in the grafted sequence.
The Ubiquitin core residues around the randomized loop 1 insert are ommitted for clarity. They span Ub1-6 and Ub11-76. Ub7-8 (TL) and Ub9-10 (TG) are included in the shown sequences, and are encoded by default in the library design.

**Round 3**

| Rank | Loop 1 | read count | read frequency | U-body name |
|---|---|---|---|---|
| **1** | **LDEGE-----YFESVYAIWGTL-----HGFSV** | **32218** | **61.66%** | **LG3-1** |
| **2** | **VEF-----YFESVYAIWGTL-----WPYVT** | **6059** | **11.60%** | **LG3-2** |
| **3** | **VFVEN-----YFESVYAIWGTL-----NPLFW** | **2142** | **4.10%** | **LG3-3** |
| 4 | VFL-----YFESVYAIWGTL-----SPLIL | 2042 | 3.91% |  |
| 5 | LFERS-----YFESVYAIWGTL-----LVPRI | 527 | 1.01% |  |
| 6 | RWAIV-----YFESVYAIWGTL-----WDFVL | 346 | 0.66% |  |
| 7 | PLSCR-----YFESVYAIWGTL-----VLSMV | 326 | 0.62% |  |
| 8 | PLRDD-----YFESVYAIWGTL-----CALSL | 294 | 0.56% |  |
| 9 | RFYRV-----YFESVYAIWGTL-----FHDYV | 292 | 0.56% |  |
| 10 | SSRRV-----YFESVYAIWGTL-----YFDFN | 282 | 0.54% |  |
| 11 | RFWRM-----YFESVYAIWGTL-----YLDLV | 281 | 0.54% |  |
| 12 | CLTTE-----YFESVYAIWGTL-----ESVLK | 223 | 0.43% |  |
| 13 | YGCET-----YFESVYAIWGTL-----FACVF | 207 | 0.40% |  |
| 14 | WMRGSTLKACTQFG | 202 | 0.39% | |
| 15 | FLTTA-----YFESVYAIWGTL-----CVLIR | 174 | 0.33% |  |
| 16 | LIATE-----YFESVYAIWGTL-----MVEAG | 165 | 0.32% |  |
| 17 | RPEGC-----YFESVYAIWGTL-----ANNISK | 150 | 0.29% |  |
| 18 | FFSVI-----YFESVYAIWGTL-----FAFYV | 144 | 0.28% |  |
| 19 | SRSWV-----YFESVYAIWGTL-----NMRGS | 122 | 0.23% |  |
| 20 | YLLVW-----YFESVYAIWGTL-----GFSVV | 114 | 0.22% |  |
| 21 | RYERW-----YFESVYAIWGTL-----WITVV | 114 | 0.22% |  |
| 22 | IYASC-----YFESVYAIWGTL-----VFLVK | 107 | 0.20% |  |
| 23 | GTAVL-----YFESVYAIWGTL-----VSYIY | 105 | 0.20% |  |
| 24 | NIHSK-----YFESVYAIWGTL-----DSLIF | 104 | 0.20% |  |
| 25 | ISSML-----YFESVYAIWGTL-----EMTWF | 95 | 0.18% |  |
| 26 | LRFHG-----YFESVYAIWGTL-----LFMLG | 95 | 0.18% |  |
| 27 | SQLFV-----YFESVYAIWGTL-----SVVFV | 91 | 0.17% |  |
| 28 | WYVWL-----YFESVYAIWGTL-----NYVYV | 90 | 0.17% |  |
| 29 | KGLAL-----YFESVYAIWGTL-----GFSVT | 90 | 0.17% |  |
| 30 | LGGSY-----YFESVYAIWGTL-----FVRFC | 89 | 0.17% |  |
| 31 | RMIGL-----YFESVYAIWGTL-----TSIDE | 88 | 0.17% |  |
| 32 | CFRGQ-----YFESVYAIWGTL-----FLIVA | 88 | 0.17% |  |
| 33 | EGISL-----YFESVYAIWGTL-----WCPRM | 87 | 0.17% |  |
| 34 | LYVDL-----YFESVYAIWGTL-----GGVMI | 82 | 0.16% |  |
| 35 | MWCGS-----YFESVYAIWGTL-----GFVLY | 82 | 0.16% |  |
| 36 | ICTVF-----YFESVYAIWGTL-----IVAVA | 81 | 0.16% |  |
| 37 | DFCLG-----YFESVYAIWGTL-----VLVTL | 80 | 0.15% |  |
| 38 | MCVIG-----YFESVYAIWGTL-----VVWSS | 80 | 0.15% |  |
| 39 | VLFLY-----YFESVYAIWGTL-----QFSGW | 79 | 0.15% |  |
| 40 | EADLI-----YFESVYAIWGTL-----FYCFC | 79 | 0.15% |  |
| 41 | MALVF-----YFESVYAIWGTL-----EWMVQ | 78 | 0.15% |  |
| 42 | TRFHD-----YFESVYAIWGTL-----GIGLG | 78 | 0.15% |  |
| 43 | SLLVW-----YFESVYAIWGTL-----IVVFS | 78 | 0.15% |  |
| 44 | DRCFV-----YFESVYAIWGTL-----VEFFG | 78 | 0.15% |  |
| 45 | HSVVC-----YFESVYAIWGTL-----IKDRL | 77 | 0.15% |  |
| 46 | RIMVW-----YFESVYAIWGTL-----WLLFS | 77 | 0.15% |  |
| 47 | VVSVY-----YFESVYAIWGTL-----VAPWV | 77 | 0.15% |  |
| 48 | SGLFS-----YFESVYAIWGTL-----TGIIV | 76 | 0.15% |  |
| 49 | EAWVH-----YFESVYAIWGTL-----NLRVS | 76 | 0.15% |  |
| 50 | VGLNW-----YFESVYAIWGTL-----YVSSE | 75 | 0.14% |  |
| 51 | WWLWL-----YFESVYAIWGTL-----LLGLG | 75 | 0.14% |  |
| 52 | NSWQN-----YFESVYAIWGTL-----ILVMR | 75 | 0.14% |  |
| 53 | EYRLF-----YFESVYAIWGTL-----VMVFY | 73 | 0.14% |  |
| 54 | CMRLSYFESVYAIW | 73 | 0.14% | |
| 55 | YVRKI-----YFESVYAIWGTL-----YTLHC | 72 | 0.14% |  |
| 56 | VVHSF-----YFESVYAIWGTL-----TTRWW | 72 | 0.14% |  |
| 57 | DEDSL-----YFESVYAIWGTL-----ELLFM | 71 | 0.14% |  |
| 58 | VVIVY-----YFESVYAIWGTL-----ALWQW | 71 | 0.14% |  |
| 59 | CYFYS-----YFESVYAIWGTL-----AFWCC | 71 | 0.14% |  |
| 60 | FMRCV-----YFESVYAIWGTL-----QFENT | 71 | 0.14% |  |
| 61 | SPALLCLLMMYLVL | 69 | 0.13% | |
| 62 | CGSLL-----YFESVYAIWGTL-----AGFWL | 69 | 0.13% |  |
| 63 | WKHSF-----YFESVYAIWGTL-----LVIDE | 69 | 0.13% |  |
| 64 | ALWVC-----YFESVYAIWGTL-----CIICS | 69 | 0.13% |  |
| 65 | FMDYK-----YFESVYAIWGTL-----LGIAY | 69 | 0.13% |  |
| 66 | LDLAL-----YFESVYAIWGTL-----CLTLC | 68 | 0.13% |  |
| 67 | YLLSH-----YFESVYAIWGTL-----KSFPC | 68 | 0.13% |  |
| 68 | GDVLA-----YFESVYAIWGTL-----LWYYC | 68 | 0.13% |  |
| 69 | WAISI-----YFESVYAIWGTL-----GVLFC | 68 | 0.13% |  |
| 70 | YSYHG-----YFESVYAIWGTL-----VLCMA | 68 | 0.13% |  |
| 71 | MRCFC-----YFESVYAIWGTL-----LVGML | 67 | 0.13% |  |
| 72 | VRTLF-----YFESVYAIWGTL-----FFLER | 67 | 0.13% |  |

**Round 4**

| Rank | Loop 1 | read count | read frequency | U-body name |
|---|---|---|---|---|
| **1** | **LDEGE-----YFESVYAIWGTL-----HGFSV** | **151215** | **95.18%** | **LG3-1** |
| **2** | **VEF-----YFESVYAIWGTL-----WPYVT** | **2421** | **1.52%** | **LG3-2** |
| 3 | WMRGSTLKACTQFG | 1208 | **0.76%** | |
| **4** | **VFVEN-----YFESVYAIWGTL-----NPLFW** | **349** | **0.22%** | **LG3-3** |
| **5** | **LDEGG-----YFESVYAIWGTL-----HGFSV** | **246** | **0.15%** | **LG3-4** |
| 6 | VFL-----YFESVYAIWGTL-----SPLIL | 204 | **0.13%** |  |
| 7 | LDEGEYFESMYAIWGTLHGFSV | 184 | **0.12%** | |
| 8 | LDEGEYFESVYTIWGTLHGFSV | 171 | **0.11%** | |
| 9 | LDEGEYFESVYAIW | 136 | **0.09%** | |
| 10 | LDEGEYFESVYAIWDTLHGFSV | 127 | **0.08%** | |
| 11 | LDERE-----YFESVYAIWGTL-----HGFSV | 121 | **0.08%** |  |
| 12 | LDEGEYFESVYAIWSTLHGFSV | 112 | **0.07%** | |
| 13 | LDKGE-----YFESVYAIWGTL-----HGFSV | 109 | **0.07%** |  |
| 14 | LDEGEYFKSVYAIWGTLHGFSV | 102 | **0.06%** | |
| 15 | LDEEE-----YFESVYAIWGTL-----HGFSV | 97 | **0.06%** |  |
| 16 | LDEGE-----YFESVYAIWGTL-----HGFSI | 94 | **0.06%** |  |
| 17 | LDEGE-----YFESVYAIWGTL-----HRFSV | 94 | **0.06%** |  |
| 18 | LDEGE-----YFESVYAIWGTL-----HGFLV | 93 | **0.06%** |  |
| 19 | LDEGE-----YFESVYAIWGTL-----HEFSV | 87 | **0.05%** |  |
| 20 | LNEGE-----YFESVYAIWGTL-----HGFSV | 86 | **0.05%** |  |
| 21 | LDEGEYFDSVYAIWGTLHGFSV | 79 | **0.05%** | |
| 22 | LDEGK-----YFESVYAIWGTL-----HGFSV | 77 | **0.05%** |  |
| 23 | LDEGEYFESVFAIWGTLHGFSV | 62 | **0.04%** | |
| 24 | LDEGEYFESVYVIWGTLHGFSV | 61 | **0.04%** | |
| 25 | LDEGEYLESVYAIWGTLHGFSV | 61 | **0.04%** | |
| 26 | LDEGEYFESVYA | 56 | **0.04%** | |
| 27 | LDEGEYFENVYAIWGTLHGFSV | 53 | **0.03%** | |
| 28 | LDEGE-----YFESVYAIWGTL-----YGFSV | 43 | **0.03%** |  |
| 29 | LDGGE-----YFESVYAIWGTL-----HGFSV | 31 | **0.02%** |  |
| 30 | LDEGSTLKACTQFG | 30 | **0.02%** | |
| 31 | LDEGEYFESVYAIWGTPHGFSV | 29 | **0.02%** | |
| 32 | FDEGE-----YFESVYAIWGTL-----HGFSV | 29 | **0.02%** |  |
| 33 | LDEGEYFESVYEIWGTLHGFSV | 29 | **0.02%** | |
| 34 | LGEGE-----YFESVYAIWGTL-----HGFSV | 28 | **0.02%** |  |
| 35 | LDEGEHFESVYAIWGTLHGFSV | 26 | **0.02%** | |
| 36 | LDEGEYFESVYAVWGTLHGFSV | 26 | **0.02%** | |
| 37 | LDEGEYFESVYAIWGILHGFSV | 26 | **0.02%** | |
| 38 | LDEGE-----YFESVYAIWGTL-----HGLSV | 25 | **0.02%** |  |
| 39 | LDEGEYFGSVYAIWGTLHGFSV | 23 | **0.01%** | |
| 40 | LDEGEYFESVYAIWGNLHGFSV | 23 | **0.01%** | |
| 41 | LDEGE-----YFESVYAIWGTL-----NGFSV | 22 | **0.01%** |  |
| 42 | IGAFVLVWPWSPG | 22 | **0.01%** | |
| 43 | VEF-----YFESVYAIWGTL-----HGFSV | 22 | **0.01%** |  |
| 44 | LDEGEYFESVYAIWGSLHGFSV | 22 | **0.01%** | |
| 45 | LDEGEYFESAYAIWGTLHGFSV | 22 | **0.01%** | |
| 46 | LDEGEYFESVYAIRGTLHGFSV | 20 | **0.01%** | |
| 47 | LDEGEYFESV | 20 | **0.01%** | |
| 48 | VFL-----YFESVYAIWGTL-----HGFSV | 20 | **0.01%** |  |
| 49 | LFERS-----YFESVYAIWGTL-----LVPRI | 19 | **0.01%** |  |
| 50 | LDEGEYFERVYAIWGTLHGFSV | 19 | **0.01%** | |
| 51 | WSFTLKACTQFG | 19 | **0.01%** | |
| 52 | LDEGE-----YFESVYAIWGTL-----RGFSV | 19 | **0.01%** |  |
| 53 | LDEGEYSESVYAIWGTLHGFSV | 19 | **0.01%** | |
| 54 | LDEGE-----YFESVYAIWGTL-----HGSSV | 18 | **0.01%** |  |
| 55 | LDEWE-----YFESVYAIWGTL-----HGFSV | 18 | **0.01%** |  |
| 56 | LDEGE-----YFESVYAIWGTL-----HGFSA | 17 | **0.01%** |  |
| 57 | LDEGEYFESVCAIWGTLHGFSV | 16 | **0.01%** | |
| 58 | PLRDD-----YFESVYAIWGTL-----CALSL | 16 | **0.01%** |  |
| 59 | LDEGEYFESVYAILGTLHGFSV | 16 | **0.01%** | |
| 60 | LDEGV-----YFESVYAIWGTL-----HGFSV | 16 | **0.01%** |  |
| 61 | SDEGE-----YFESVYAIWGTL-----HGFSV | 16 | **0.01%** |  |
| 62 | LDEGEYFESVR | 15 | **0.01%** | |
| 63 | LDEGEYFESVYATWGTLHGFSV | 15 | **0.01%** | |
| 64 | LDEGE-----YFESVYAIWGTL-----HWFSV | 15 | **0.01%** |  |
| 65 | LDEGEYFEGVYAIWGTLHGFSV | 15 | **0.01%** | |
| 66 | LDEGEYLKACTQFG | 14 | **0.01%** | |
| 67 | RWAIV-----YFESVYAIWGTL-----WDFVL | 14 | **0.01%** |  |
| 68 | VFVEN-----YFESVYAIWGTL-----HGFSV | 14 | **0.01%** |  |
| 69 | LDEGEYFESVYAICGTLHGFSV | 14 | **0.01%** | |
| 70 | LDEGEYFESVYAIWGTMHGFSV | 14 | **0.01%** | |
| 71 | LDEGE-----YFESVYAIWGTL-----HGFPV | 14 | **0.01%** |  |
| 72 | LDEGEYFESVYAIWGALHGFSV | 12 | **0.01%** | |

**Round 5**

| Rank | Loop 1 | read count | read frequency | U-body name |
|---|---|---|---|---|
| **1** | **LDEGE-----YFESVYAIWGTL-----HGFSV** | **144445** | **96.88%** | **LG3-1** |
| 2 | WMRGSTLKACTQFG | 1219 | **0.82%** | |
| **3** | **VEF-----YFESVYAIWGTL-----WPYVT** | **302** | **0.20%** | **LG3-2** |
| **4** | **LDEGG-----YFESVYAIWGTL-----HGFSV** | **208** | **0.14%** | **LG3-4** |
| 5 | LDEGEYFESVYAIW | 177 | **0.12%** | |
| 6 | LDEGEYFESVYTIWGTLHGFSV | 170 | **0.11%** | |
| 7 | LDEGEYFDSVYAIWGTLHGFSV | 153 | **0.10%** | |
| 8 | LDEGE-----YFESVYAIWGTL-----HGFSI | 149 | **0.10%** |  |
| 9 | LDEGEYFESMYAIWGTLHGFSV | 149 | **0.10%** | |
| 10 | LDEGEYFESVYAIWDTLHGFSV | 88 | **0.06%** | |
| 11 | LDEGEYLESVYAIWGTLHGFSV | 79 | **0.05%** | |
| 12 | LDERE-----YFESVYAIWGTL-----HGFSV | 77 | **0.05%** |  |
| 13 | LDEGE-----YFESVYAIWGTL-----HRFSV | 71 | **0.05%** |  |
| 14 | LDEEE-----YFESVYAIWGTL-----HGFSV | 67 | **0.04%** |  |
| 15 | LDEGE-----YFESVYAIWGTL-----HGFLV | 66 | **0.04%** |  |
| 16 | LDKGE-----YFESVYAIWGTL-----HGFSV | 65 | **0.04%** |  |
| 17 | LDEGEYFKSVYAIWGTLHGFSV | 62 | **0.04%** | |
| 18 | LDEGE-----YFESVYAIWGTL-----HEFSV | 61 | **0.04%** |  |
| 19 | LDEGEYFESVYA | 55 | **0.04%** | |
| 20 | LDEGEYFESVYAIWSTLHGFSV | 54 | **0.04%** | |
| 21 | LNEGE-----YFESVYAIWGTL-----HGFSV | 53 | **0.04%** |  |
| 22 | LDEGK-----YFESVYAIWGTL-----HGFSV | 52 | **0.03%** |  |
| 23 | LDEGEYFESVFAIWGTLHGFSV | 43 | **0.03%** | |
| 24 | LDEGEYFESVYVIWGTLHGFSV | 42 | **0.03%** | |
| 25 | LDEGEYFESVYAIWGTMHGFSV | 39 | **0.03%** | |
| 26 | LGEGE-----YFESVYAIWGTL-----HGFSV | 38 | **0.03%** |  |
| 27 | LDEGE-----YFESVYAIWGTL-----YGFSV | 38 | **0.03%** |  |
| 28 | LDEGEYFESV | 36 | **0.02%** | |
| 29 | LDEGEYFENVYAIWGTLHGFSV | 35 | **0.02%** | |
| 30 | LDEGEYFESAYAIWGTLHGFSV | 32 | **0.02%** | |
| 31 | LDEGEYFESVYEIWGTLHGFSV | 32 | **0.02%** | |
| 32 | LDEGEYFESVYAVWGTLHGFSV | 30 | **0.02%** | |
| 33 | LDEGEYFESVYAIWGTPHGFSV | 26 | **0.02%** | |
| 34 | LDEGEYFESVYAILGTLHGFSV | 25 | **0.02%** | |
| 35 | LDGGE-----YFESVYAIWGTL-----HGFSV | 25 | **0.02%** |  |
| 36 | LDEGEYFERVYAIWGTLHGFSV | 24 | **0.02%** | |
| 37 | LDEGEYFESVYAIWGNLHGFSV | 24 | **0.02%** | |
| 38 | LDEGSTLKACTQFG | 24 | **0.02%** | |
| 39 | FDEGE-----YFESVYAIWGTL-----HGFSV | 23 | **0.02%** |  |
| 40 | LDEGE-----YFESVYAIWGTL-----NGFSV | 23 | **0.02%** |  |
| 41 | LDEGEHFESVYAIWGTLHGFSV | 23 | **0.02%** | |
| 42 | LDEGEYFESVYAIRGTLHGFSV | 23 | **0.02%** | |
| 43 | LDEGEYLKACTQFG | 22 | **0.01%** | |
| 44 | LDEGEYFGSVYAIWGTLHGFSV | 22 | **0.01%** | |
| 45 | LDEGE-----YFESVYAIWGTL-----HGLSV | 21 | **0.01%** |  |
| 46 | LDEGEYFESVYAIWGILHGFSV | 21 | **0.01%** | |
| 47 | SDEGE-----YFESVYAIWGTL-----HGFSV | 20 | **0.01%** |  |
| 48 | LDEGE-----YFESVYAIWGTL-----HGSSV | 19 | **0.01%** |  |
| 49 | LDEGE-----YFESVYAIWGTL-----RGFSV | 18 | **0.01%** |  |
| 50 | LDEGEYFESVYAIWGSLHGFSV | 17 | **0.01%** | |
| 51 | LDEGEYSESVYAIWGTLHGFSV | 17 | **0.01%** | |
| 52 | LDEGEYFESVYAICGTLHGFSV | 17 | **0.01%** | |
| 53 | LDEGE-----YFESVYAIWGTL-----HGFPV | 17 | **0.01%** |  |
| 54 | LDEGEYFESVCAIWGTLHGFSV | 17 | **0.01%** | |
| 55 | LDEGEYFESVHAIWGTLHGFSV | 16 | **0.01%** | |
| **56** | **VFVEN-----YFESVYAIWGTL-----NPLFW** | **16** | **0.01%** | **LG3-3** |
| 57 | LDEGEFFESVYAIWGTLHGFSV | 16 | **0.01%** | |
| 58 | LDEGE-----YFESVYAIWGTL-----HGFS | 15 | **0.01%** |  |
| 59 | LDDGE-----YFESVYAIWGTL-----HGFSV | 15 | **0.01%** |  |
| 60 | WMKGSTLKACTQFG | 15 | **0.01%** | |
| 61 | VFL-----YFESVYAIWGTL-----SPLIL | 15 | **0.01%** |  |
| 62 | LEEGE-----YFESVYAIWGTL-----HGFSV | 14 | **0.01%** |  |
| 63 | LDEGE-----YFESVYAIWGTL-----LGFSV | 14 | **0.01%** |  |
| 64 | LDEGEYFESVR | 14 | **0.01%** | |
| 65 | LDEGEYFESVYAIWGALHGFSV | 13 | **0.01%** | |
| 66 | LDEGEYFESVYATWGTLHGFSV | 12 | **0.01%** | |
| 67 | LDEGEYFESEYAIWGTLHGFSV | 12 | **0.01%** | |
| 68 | LDEGE-----YFESVYAIWGTL-----HGFSA | 12 | **0.01%** |  |
| 69 | DEGE-----YFESVYAIWGTL-----HGFSV | 12 | **0.01%** |  |
| 70 | LDEGECFESVYAIWGTLHGFSV | 11 | **0.01%** | |
| 71 | LDEGE-----YFESVYAIWGTL----- | 11 | **0.01%** |  |
| 72 | LDEGV-----YFESVYAIWGTL-----HGFSV | 10 | **0.01%** |  |

| | | | | |
|---|---|---|---|---|
| 73 | LMFAF-----YFESVYAIWGTL-----GGYFI | 67 | 0.13% |  |
| 74 | AMHKL-----YFESVYAIWGTL-----MVFWS | 67 | 0.13% |  |
| 75 | VNLSA-----YFESVYAIWGTL-----ELYFV | 67 | 0.13% |  |
| 76 | ESGRS-----YFESVYAIWGTL-----FGLCV | 67 | 0.13% |  |
| 77 | FVGAVCLALLVFL | 66 | 0.13% | |
| 78 | VSAVW-----YFESVYAIWGTL-----IHFMI | 66 | 0.13% |  |
| 79 | DVVYM-----YFESVYAIWGTL-----RLFYS | 66 | 0.13% |  |
| 80 | WNKSC-----YFESVYAIWGTL-----TGLNG | 66 | 0.13% |  |
| 81 | CVCHF-----YFESVYAIWGTL-----NSLAP | 66 | 0.13% |  |
| 82 | WSPSF-----YFESVYAIWGTL-----LVSMW | 66 | 0.13% |  |
| 83 | VVESQ-----YFESVYAIWGTL-----VWMDF | 66 | 0.13% |  |
| 84 | IDRVT-----YFESVYAIWGTL-----IFGCF | 66 | 0.13% |  |
| 85 | YTLAG-----YFESVYAIWGTL-----MRSVV | 66 | 0.13% |  |
| 86 | ALWVN-----YFESVYAIWGTL-----ICLIS | 65 | 0.12% |  |
| 87 | VICSA-----YFESVYAIWGTL-----FYVQF | 65 | 0.12% |  |
| **88** | **LDEGG-----YFESVYAIWGTL-----HGFSV** | **65** | **0.12%** | **LG3-4** |
| 89 | YRWCL-----YFESVYAIWGTL-----LFLWC | 65 | 0.12% |  |
| 90 | VSLII-----YFESVYAIWGTL-----DIVCG | 65 | 0.12% |  |
| 91 | TFLRA-----YFESVYAIWGTL-----SVWGC | 64 | 0.12% |  |
| 92 | SFCWF-----YFESVYAIWGTL-----PAIYE | 64 | 0.12% |  |
| 93 | EVLSL-----YFESVYAIWGTL-----VWPVI | 64 | 0.12% |  |
| 94 | AGVRC-----YFESVYAIWGTL-----FADFV | 64 | 0.12% |  |
| 95 | WCKLV-----YFESVYAIWGTL-----FSLRV | 64 | 0.12% |  |
| 96 | ALRIR-----YFESVYAIWGTL-----GYGNK | 64 | 0.12% |  |
| 97 | HFGTR-----YFESVYAIWGTL-----CLLAF | 63 | 0.12% |  |
| 98 | KHFML-----YFESVYAIWGTL-----MVMVD | 63 | 0.12% |  |
| 99 | LLCVN-----YFESVYAIWGTL-----IWNWV | 63 | 0.12% |  |
| 100 | GIFLL-----YFESVYAIWGTL-----GL | 63 | 0.12% |  |
| | *total reads (top 100)* | 52252 | | |

| | | | |
|---|---|---|---|
| 73 | LDEGE-----YFESVYAIWGTL-----WPYVT | 12 | **0.01%** |
| 74 | LDDGE-----YFESVYAIWGTL-----HGFSV | 11 | **0.01%** |
| 75 | CLTTE-----YFESVYAIWGTL-----ESVLK | 11 | **0.01%** |
| 76 | LDEGEFFESVYAIWGTLHGFSV | 11 | **0.01%** |
| 77 | LYEGE-----YFESVYAIWGTL-----HGFSV | 11 | **0.01%** |
| 78 | LDEGEYFESVHAIWGTLHGFSV | 11 | **0.01%** |
| 79 | LEEGE-----YFESVYAIWGTL-----HGFSV | 10 | **0.01%** |
| 80 | RFYRV-----YFESVYAIWGTL-----FHDYV | 10 | **0.01%** |
| 81 | LDEGE-----YFESVYAIWGTL-----HGFS | 10 | **0.01%** |
| 82 | LDEGE-----YFESVYAIWGTL-----HGFTV | 9 | **0.01%** |
| 83 | PLSCR-----YFESVYAIWGTL-----VLSMV | 9 | **0.01%** |
| 84 | LDEVE-----YFESVYAIWGTL-----HGFSV | 9 | **0.01%** |
| 85 | LDEGE-----YFESVYAIWGTL-----HVFSV | 9 | **0.01%** |
| 86 | LDEGECFESVYAIWGTLHGFSV | 8 | **0.01%** |
| 87 | LDEGE-----YFESVYAIWGTL-----LGFSV | 8 | **0.01%** |
| 88 | LGGSY-----YFESVYAIWGTL-----FVRFC | 8 | **0.01%** |
| 89 | LDEGE-----YFESVYAIWGTL----- | 8 | **0.01%** |
| 90 | LDEGEYFVSVYAIWGTLHGFSV | 7 | **0.00%** |
| 91 | LDVGE-----YFESVYAIWGTL-----HGFSV | 7 | **0.00%** |
| 92 | RPEGC-----YFESVYAIWGTL-----ANNISK | 7 | **0.00%** |
| 93 | SSRRV-----YFESVYAIWGTL-----YFDFN | 7 | **0.00%** |
| 94 | LDEGEYYESVYAIWGTLHGFSV | 7 | **0.00%** |
| 95 | LDEGE-----YFESVYAIWGTL-----HGVSV | 7 | **0.00%** |
| 96 | LDEGEYFESVNAIWGTLHGFSV | 7 | **0.00%** |
| 97 | MDEGE-----YFESVYAIWGTL-----HGFSV | 6 | **0.00%** |
| 98 | LVEGE-----YFESVYAIWGTL-----HGFSV | 6 | **0.00%** |
| 99 | LMRGSTLKACTQFG | 6 | **0.00%** |
| 100 | QRLEL-----YFESVYAIWGTL-----SWTLE | 6 | **0.00%** |
| | *total reads (top 100)* | 158865 | |

| | | | |
|---|---|---|---|
| 73 | LYEGE-----YFESVYAIWGTL-----HGFSV | 10 | **0.01%** |
| 74 | LDEGEYFEGVYAIWGTLHGFSV | 10 | **0.01%** |
| 75 | LVEGE-----YFESVYAIWGTL-----HGFSV | 10 | **0.01%** |
| 76 | LDEGEYFVSVYAIWGTLHGFSV | 9 | **0.01%** |
| 77 | LDVGE-----YFESVYAIWGTL-----HGFSV | 9 | **0.01%** |
| 78 | LDEGEYFESVYAIWVTLHGFSV | 9 | **0.01%** |
| 79 | LDEGEYFESVYAIWCTLHGFSV | 9 | **0.01%** |
| 80 | LDEGE-----YFESVYAIWGTL-----QGFSV | 8 | **0.01%** |
| 81 | LDEGEYFEIVYAIWGTLHGFSV | 8 | **0.01%** |
| 82 | LDEGEYFESVYAIG | 8 | **0.01%** |
| 83 | LDEGE-----YFESVYAIWGTL-----HWFSV | 7 | **0.00%** |
| 84 | LDEGEYFESVNAIWGTLHGFSV | 7 | **0.00%** |
| 85 | LDEWE-----YFESVYAIWGTL-----HGFSV | 7 | **0.00%** |
| 86 | LDEGGTLKACTQFG | 7 | **0.00%** |
| 87 | LDEGE-----YFESVYAIWGTL-----HGFTV | 7 | **0.00%** |
| 88 | LDEGEYFESVYAIWGTQHGFSV | 7 | **0.00%** |
| 89 | LDEGE-----YFESVYAIWGTL-----HG | 7 | **0.00%** |
| 90 | LDEGEYFESVYAIWRTLHGFSV | 6 | **0.00%** |
| 91 | LDEVE-----YFESVYAIWGTL-----HGFSV | 6 | **0.00%** |
| 92 | LDEGE-----YFESVYAIWGTL-----HGISV | 5 | **0.00%** |
| 93 | VEF-----YFESVYAIWGTL-----HGFSV | 5 | **0.00%** |
| 94 | LDEGE-----YFESVYAIWGTL-----HVFSV | 5 | **0.00%** |
| 95 | LDEAE-----YFESVYAIWGTL-----HGFSV | 5 | **0.00%** |
| 96 | LDEGEYFESVYSIWGTLHGFSV | 5 | **0.00%** |
| 97 | WMRGSTLKACTPFG | 5 | **0.00%** |
| 98 | WMRESTLKACTQFG | 4 | **0.00%** |
| 99 | LDEGEYYESVYAIWGTLHGFSV | 4 | **0.00%** |
| 100 | LDEGEYFESVYPIWGTLHGFSV | 4 | **0.00%** |
| | *total reads (top 100)* | 149091 | |
