## Supplementary Table 2 for "Random *in vitro* Protein single-Loop Engineering (RiPLE) by mRNA display"

List of oligonucleotides used this study

**Library preparation and PCR primers**

| **Name** | **Description** | **Comment** | **DNA sequence** |
|---|---|---|---|
| G10_Ub6_NNK15_Ub14.F102 | RiPLE 1st step extension PCR fwd | Identical designs for NNK8-14 | AGAAGGAGATATACATATGCAGATCTTCGTCAAGNNKNNKNNKNNKNNKNNKNNKNNKNNKNNKNNKNNKNNKNNKNNKAAAACCATAACTCTGGAAGTGGA |
| Ub11-44.R102 | RiPLE 1st step extension PCR rev | | GATCAGACGCTGCTGATCCGGCGGGATGCCTTCTTTATCCTGGATTTTCGCTTTCACGTTTTCGATGGTATCGCTCGGTTCCACTTCCAGAGTTATGGTTTT |
| T7G10MUb6.F63 | 2nd step extension PCR fwd | Also used for qPCR during selection | TAATACGACTCACTATAGGGTTAACTTTAAGAAGGAGATATACATATGCAGATCTTCGTCAAG |
| Ub38-76_SSG2an14.R154 | 2nd step extension PCR rev | | TTTCCGCCCCCCGTCCTAACCGCTAGAGCCAGAGCTACCTCCACGCAGTCTTAAGACCAGATGCAGGGTCGACTCTTTCTGGATGTTATAATCAGACAGaGTGCGaCCATCTTCCAGCTGTTTGCCCGCAAAGATCAGACGCTGCTGATCCGGC |
| LG1-B2-F5.F81 | LG1 1st step extension PCR fwd | Identical designs for NNK1-4 | GAGATATACATATGCAGATCTTCGTCAAGACGTTANNKNNKNNKNNKNNKTATTGGTATTGGACCGGCAACAACTGGCCGT |
| LG1-B2-R5.R144 | LG1 1st step extension PCR rev | Identical designs for NNK1-4 | AGATCAGACGCTGCTGATCCGGCGGGATGCCTTCTTTATCCTGGATTTTCGCTTTCACGTTTTCGATGGTATCGCTCGGTTCCACTTCCAGAGTTATGGTTTTACCGGTMNNMNNMNNMNNMNNCACATACGGCCAGTTGTTGC |
| LG2-B2-F5.F83 | LG2 1st step extension PCR fwd | Identical designs for NNK1-4 | GAGATATACATATGCAGATCTTCGTCAAGACGTTANNKNNKNNKNNKNNKTATGTGATGTATTGGTATTGGACCGGCAACAAC |
| LG2-B2-R5.R147 | LG2 1st step extension PCR rev | Identical designs for NNK1-4 | AGATCAGACGCTGCTGATCCGGCGGGATGCCTTCTTTATCCTGGATTTTCGCTTTCACGTTTTCGATGGTATCGCTCGGTTCCACTTCCAGAGTTATGGTTTTACCGGTMNNMNNMNNMNNMNNCCAGTTGTTGCCGGTCCAATACC |
| LG3-GN13-F5-R5.F116 | LG3 1st step extension PCR fwd | Identical designs for NNK1-4 | GATATACATATGCAGATCTTCGTCAAGNNKNNKNNKNNKNNKTACTTTGAAAGCGTGTACGCAATTTGGGGCACCCTGNNKNNKNNKNNKNNKAAAACCATAACTCTGGAAGTGGA |
| | *For LG3, Ub11-44.R102 used for 1st step extension* | | |
| SSG2an13.R39 | Reverse primer for qPCR during selection | | TTTCCGCCCCCCGTCCTAACCGCTAGAGCCAGAGCT |

**Sequencing primer**

| **Name** | **Description** | **Comment** | **DNA sequence** |
|---|---|---|---|
| Forward, Rd1T7g10M.F70 | Forward extension primer for NGS | | CACTCTTTCCCTACACGACGCTCTTCCGATCTTAATACGACTCACTATAGGGTTAACTTTAAGAAGGAGA |
| Reverse, an13Rd2.R49 | Reverse extension primer for NGS | | AGGACGGGGGGCGGAAAAGATCGGAAGAGCACACGTCTGAACTCCAGTC |
| P5S5**Rd1.F57 | Forward barcoding primer for NGS | "N" & length change with barcode | AATGATACGGCGACCACCGAGATCTACACNNNNNNNNACACTCTTTCCCTACACGAC |
| Rd2N7**P7.R52 | Reverse barcoding primer for NGS | "N" & length change with barcode | CAAGCAGAAGACGGCATACGAGATNNNNNNNNGTGACTGGAGTTCAGACGTG |
